# Epigenetic mechanisms controlling human leukemia stem cells and therapy resistance

**DOI:** 10.1101/2022.09.22.509005

**Authors:** Sumiko Takao, Victor Morell, Masahiro Uni, Alicia Slavit, Sophia Rha, Shuyuan Cheng, Laura K Schmalbrock, Fiona C Brown, Sergi Beneyto-Calabuig, Richard P Koche, Lars Velten, Alex Kentsis

**Affiliations:** Molecular Pharmacology Program, Sloan Kettering Institute, Memorial Sloan Kettering Cancer Center, New York, NY, USA; Tow Center for Developmental Oncology, Department of Pediatrics, Memorial Sloan Kettering Cancer Center, NY, USA; Centre for Genomic Regulation (CRG), The Barcelona Institute of Science and Technology, Dr. Aiguader 88, Barcelona 08003, Spain; Universitat Pompeu Fabra (UPF), Barcelona, Spain; Center for Epigenetics Research, Sloan Kettering Institute, New York, NY, USA; Departments of Pediatrics, Pharmacology, and Physiology & Biophysics, Weill Medical College of Cornell University, NY, USA

## Abstract

Many human cancers, including acute myeloid leukemia (AML), arise from mutations of stem and progenitor cells. Immunophenotypic profiling has shown that leukemias develop hierarchically, with mutations in leukemia stem cells associated with disease propagation and relapse^1,2^. Although leukemia initiating cells can be enriched using cell surface markers, their frequency tends to be variable and low, obscuring mechanisms and hindering effective therapies^3,4^. To define AML stem cells in human patients, we performed functional genomic profiling of diverse leukemias using label tracing techniques designed to preserve hematopoietic stem cell (HSC) function in vivo. We found that propagation of human AML is mediated by a rare but distinct quiescent label-retaining cell (LRC) population that evades detection by currently known immunophenotypic markers. We show that human AML LRC quiescence is reversible, sparing genetic clonal competition that maintains its epigenetic inheritance. LRC quiescence is defined by distinct promoter-centered chromatin and gene expression dynamics and controlled by a distinct AP-1/ETS transcription factor network, including JUN in particular, which is associated with disease persistence and chemotherapy resistance in diverse patients. These results enable prospective isolation and functional genetic manipulation of immunophenotypically-varied leukemia stem cells in human patient specimens, as well as establish key functions of epigenetic plasticity in leukemia development and therapy resistance. We anticipate that these findings will lead to the elucidation of essential properties of leukemia stem cell quiescence and the design of therapeutic strategies for their clinical identification and control.

---

Although the treatment of AML continues to improve, most patients develop disease that is refractory to intensive chemotherapy^5–7^. Leukemia development, evolution, and chemotherapy resistance have been attributed to leukemia stem cells, originally identified as rare cells that can propagate human leukemias upon transplantation in immunodeficient mice^1,2^. While leukemia initiating cells can be enriched using cell surface markers, their frequency tends to be variable and low (less than 1 in 1000 cells), obscuring mechanisms and hindering effective therapies^3,4^. Molecular mechanisms of leukemia stem cell development and therapy resistance have been investigated in genetically-engineered leukemias in mice^8–11^, but there remains uncertainty about their relevance to human AML.

## Prospective isolation of human patient AML stem cells

To define molecular mechanisms of human AML stem cells, we used orthotopic transplantation in immunodeficient mice in combination with chemical label tracing and functional genetics of primary human patient leukemia specimens. First, we assessed 48 human patient leukemias obtained at disease diagnosis or relapse by orthotopic transplantation in *NOD.Cg-Prkdc^scid^ Il2rg^tm1Wjl^*/SzJ (NSG) mice. Overall engraftment efficiency was 46% and 70% in primary and secondary transplants, respectively (Extended Data Table 1). Consistent with prior observations^12,13^, serial engraftment and transplantation were higher for specimens obtained from patients with chemotherapy resistance, as compared to those who achieved remission upon standard-of-care chemotherapy treatment (80 versus 60%; Extended Data Table 1).

Based on prior studies using genetic and chemical label tracing approaches for the isolation of quiescent or dormant cells^14–20^, we optimized the carboxyfluorescein succinimidyl ester (CFSE) chemical label tracing technique by maximizing covalent cellular protein labeling while avoiding any measurable effects on the viability of hematopoietic cells (Extended Data Figure 1a-b). When applied to primary hematopoietic cells isolated from the bone marrow of healthy mice, CFSE labeling preserved normal stem cell function, as evident from complete rescue of lethally irradiated wild-type mice upon transplantation of CFSE-labeled hematopoietic stem and progenitor cells (Extended Data Figure 1c). We confirmed that these labeling conditions also preserved primary human leukemia initiating cells in their ability to initiate disease upon orthotopic transplantation in NSG mice by CFSE-labeled bulk patient leukemia cells as compared to their unlabeled controls (Extended Data Figure 1d).

We hypothesized that CFSE label retention would specifically identify leukemia stem cells, given its ability to detect non-dividing cells with quiescent proteome turnover (Figure 1a-b). To test this idea, we selected genetically diverse patient AML specimens that exhibited short latency in serial orthotopic mouse transplants (Extended Data Table 2). First, we confirmed that CFSE labeling and orthotopic transplantation of human patient leukemia cells in NSG mice identified non-proliferating label retaining cells (LRCs), as measured by 5-ethynyl-2’-deoxyuridine (EdU) incorporation and fluorescence-activated cell scanning (Extended Data Figure 1e). Consistent with this, LRCs also exhibited little to no detectable apoptosis, as measured using cleaved caspase 3 intracellular staining, in contrast to proliferating non-LRCs some of which undergo apoptosis (Extended Data Figure 1g). We also analyzed cell cycle status of LRCs using Hoechst 33342 and Pyronin Y (H-Y) staining, which label DNA and RNA content, respectively^21^. Consistent with their quiescence, most LRCs exhibited H-Y staining of G0 cell cycle phase, in contrast to G1 and G2M cell cycle phase H-Y features of non-LRCs (Extended Data Figure 2).

**Figure 1.**
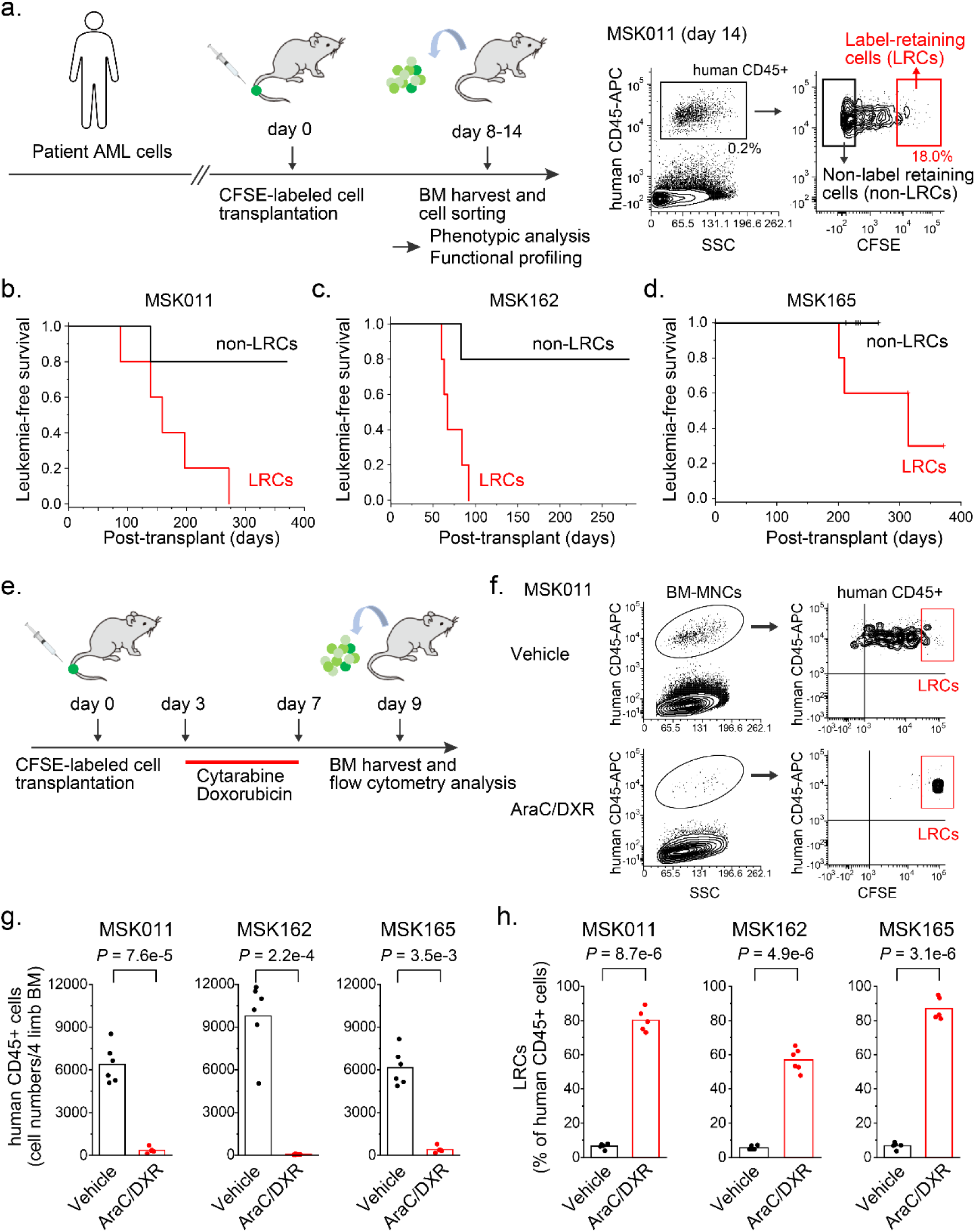
Quiescent human AML patient cells maintain leukemia initiation, propagation and chemotherapy resistance. **a.** Experimental design to prospectively isolate quiescent AML cells from human patients using optimized CFSE labeling and orthotopic transplantation in immunodeficient mice (left panel). Representative flow cytometry analysis of quiescent label-retaining human CD45-positive AML cells (LRCs, red box) with high CFSE fluorescence, as compared to their non-label retaining cells (non-LRCs, black box) that have lost CFSE fluorescence through cell division and proteome turnover (right panel). **b.** Leukemia-free survival of mice secondarily transplanted with equal numbers (900 cells/mouse, 5 mice for each group) of MSK011 patient AML LRCs (red) or non-LRCs (black), where LRCs initiate fully penetrant leukemia, and non-LRCs do not (log-rank *p* = 0.021). **c.** Leukemia-free survival of mice tertiarily transplanted with equal numbers (1,000 cells/mouse, 5 mice for each group) of MSK162 patient AML LRCs (red) or non-LRCs (black), where LRCs propagate and initiate leukemia, whereas non-LRCs largely do not (log-rank *p* = 0.013). **d.** Leukemia-free survival of mice tertiarily transplanted with equal numbers (360 cells/mouse, 5 mice for each group) of MSK165 patient AML LRCs (red) or non-LRCs (black), where LRCs propagate and initiate leukemia in 60% of mice, whereas non-LRCs do not (limiting dilution analysis (LDA) *p* = 0.019). **e.** Experimental design for the analysis of mice transplanted with CFSE-labeled human patient AMLs and treated with cytarabine (AraC) and doxorubicin (DXR). **f.** Representative flow cytometry plots to analyze LRC frequencies in bone marrow human leukemia cells isolated from mice transplanted with CFSE-labeled MSK011 patient AML cells and treated with AraC and DXR chemotherapy or vehicle (refer to Extended Data Figure 6a-b for MSK162 and MSK165). **g.** Combined AraC and DXR chemotherapy treatment reduces bone marrow disease burden of human CD45-positive MSK011 (left panel), MSK162 (center panel) and MSK165 (right panel) AML cell numbers in mice (t-test *p* = 7.6 x 10^-5^, 2.2 x 10^-4^, and 3.5 x 10^-3^, respectively). Bars represent mean values of 6 biological replicates. **h.** LRC frequencies of MSK011 (left panel), MSK162 (center panel) and MSK165 (right panel) patient AML cells are significantly increased upon AraC and DXR chemotherapy (red) as compared to vehicle-treated controls (black), exhibiting that these patient AML LRCs are resistant to AraC and DXR chemotherapy (t-test *p* = 8.7 x 10^-6^, 4.9 x 10^-6^ and 3.1 x 10^-6^, respectively), in contrast to non-LRCs that are largely eradicated by chemotherapy treatment (refer to Extended Data Figure 6e). Bars represent mean values of measurement of 6 biological replicates.

## Initiation of human patient leukemias is mediated by quiescent and chemoresistant LRCs

To determine whether patient LRCs have leukemia stem cell properties, we transplanted equal numbers of LRCs and non-LRCs into secondary recipient mice using three diverse patient AMLs, MSK011, MSK162 and MSK165 (Figure 1a, Extended Data Figure 3a-b and Extended Data Table 2). All mice transplanted with MSK011 LRCs developed leukemias (900 LRCs/mouse), which was confirmed by human CD45-specific staining using fluorescence-activated cell scanning (Extended Data Figure 3c), while most non-LRC transplanted mice remained disease free (log-rank *p* = 0.021; Figure 1b). Despite limiting cell numbers (360 cells/mouse) for MSK165, 60% of LRC-transplanted mice also developed leukemia, while non-LRC transplanted mice did not (limiting dilution analysis (LDA) *p* = 0.019; Figure 1d). Whereas both MSK162 LRCs and non-LRCs (800 cells/mouse) caused disease initially, only LRCs caused leukemias in tertiary recipients and non-LRCs did not (log-rank *p* = 0.013; Figure 1c and Extended Data Figure 3d).

Although MSK011 LRCs showed relatively high and low expression of CD34 and CD38, respectively, in agreement with prior observations^2^, no established surface marker combinations, including lymphoid-primed multipotent progenitor (LMPP)-like (CD34+CD38-CD45RA+) and granulocyte-monocyte progenitor (GMP)-like (CD34+CD38+CD123+CD45RA+), could be used to reliably discriminate LRCs from non-LRCs (Extended Data Figure 4a and 4d-e). We found similar results with MSK162 and MSK165 leukemias; in fact, MSK162 leukemia exhibited essentially no measurable CD34+CD38- cells (Extended Data Figure 4b-d).

Since leukemia stem cells have been reported to exhibit low reactive oxygen species (ROS) levels, we also measured ROS levels using the oxidation-sensitive fluorogenic probe CellROX^22,23^. MSK165 LRCs exhibited significantly lower CellROX fluorescence as compared to its non-LRCs (mean fluorescence intensity of 1744 vs 4864, respectively; t-test *p* = 5 x 10^-3^; Extended Data Figure 5), but other patient leukemias showed more variable differences, and MSK162 LRCs and non-LRCs had similar CellROX fluorescence activity (mean fluorescence intensity of 5535 vs 5427, respectively; t-test *p* = 0.92; Extended Data Figure 5). Thus, initiation and maintenance of diverse human patient leukemias are mediated by a rare but distinct quiescent LRC population with varied cell surface markers and ROS activity.

Are LRCs chemotherapy resistant? To determine this, we used a combined cytarabine (AraC) and doxorubicin (DXR) treatment regimen of NSG mice developed to model induction chemotherapy used clinically for human patients^24^. Analysis of bone marrow of AraC- and DXR-treated mice transplanted with CFSE-labeled human patient AMLs showed that while chemotherapy significantly reduced total leukemia disease burden, more than 60-80% of residual chemotherapy-resistant cells were comprised of LRCs (t-test *p* = 8.7 x 10^-6^, 4.9 x 10^-6^, and 3.1 x 10^-6^ for MSK011, MSK162 and MSK165 patient specimens, respectively; Figure 1e-h and Extended Data Figure 6a-e), in agreement with other studies^19,20^. Notably, we also found similar results when CFSE-labeled patient leukemia cell-transplanted mice were treated with the CBP/p300 acetyltransferase inhibitor A-485 (Extended Data Figure 6f-i). Thus, LRCs comprise a therapy-resistant reservoir, consistent with their cellular quiescence.

## Human patient AML cell quiescence is reversible

We next investigated whether patient LRC quiescence is associated with specific genetic clones. First, we identified leukemia disease-defining mutations for MSK011, MSK162, and MSK165 patient leukemias using high-coverage DNA sequencing of 585 genes recurrently mutated in hematologic malignancies (Extended Data Table 2)^25^. This analysis also identified 18, 21, and 52 single nucleotide and short insertion and deletion variants with varied allele frequencies (VAF), reflecting the respective specific clonal architectures of these leukemias (Extended Data Table 3; Figure 2a-c). These genetic variants included pathogenically cooperating mutations, such as *KRAS G12C* and *KMT2A-*translocation in MSK011, *KAT6A*-translocation and *FLT3 D835Y* mutation in MSK162, and *WT1 S381\** and *FLT3-ITD E604_F605ins* mutations in MSK165 leukemias. We reasoned that if LRC quiescence is caused by genetic clonal evolution, then specific subclonal mutations should segregate in LRCs versus non-LRCs, as measured by their relative VAFs (Extended Data Table 3; Figure 2a-c). No subclonal mutations exhibited significant association with patient LRCs. In contrast, several subclones were significantly enriched in non-LRCs, such as *FLT3 D835Y* in MSK162 and *FLT3*-*ITD E604_F605ins* in MSK165 patient leukemias (non-LRC 76% versus 0%, and 66% versus 31%, respectively; Extended Data Table 3; Figure 2a-c). This explains the tendency of *FLT3*-mutant subclones to be depleted by chemotherapy, and is consistent with recent measurements of clonal evolution using single-cell sequencing^26–28^. Thus, genetic clonal evolution is not required for LRC quiescence in transplantation studies.

**Figure 2.**
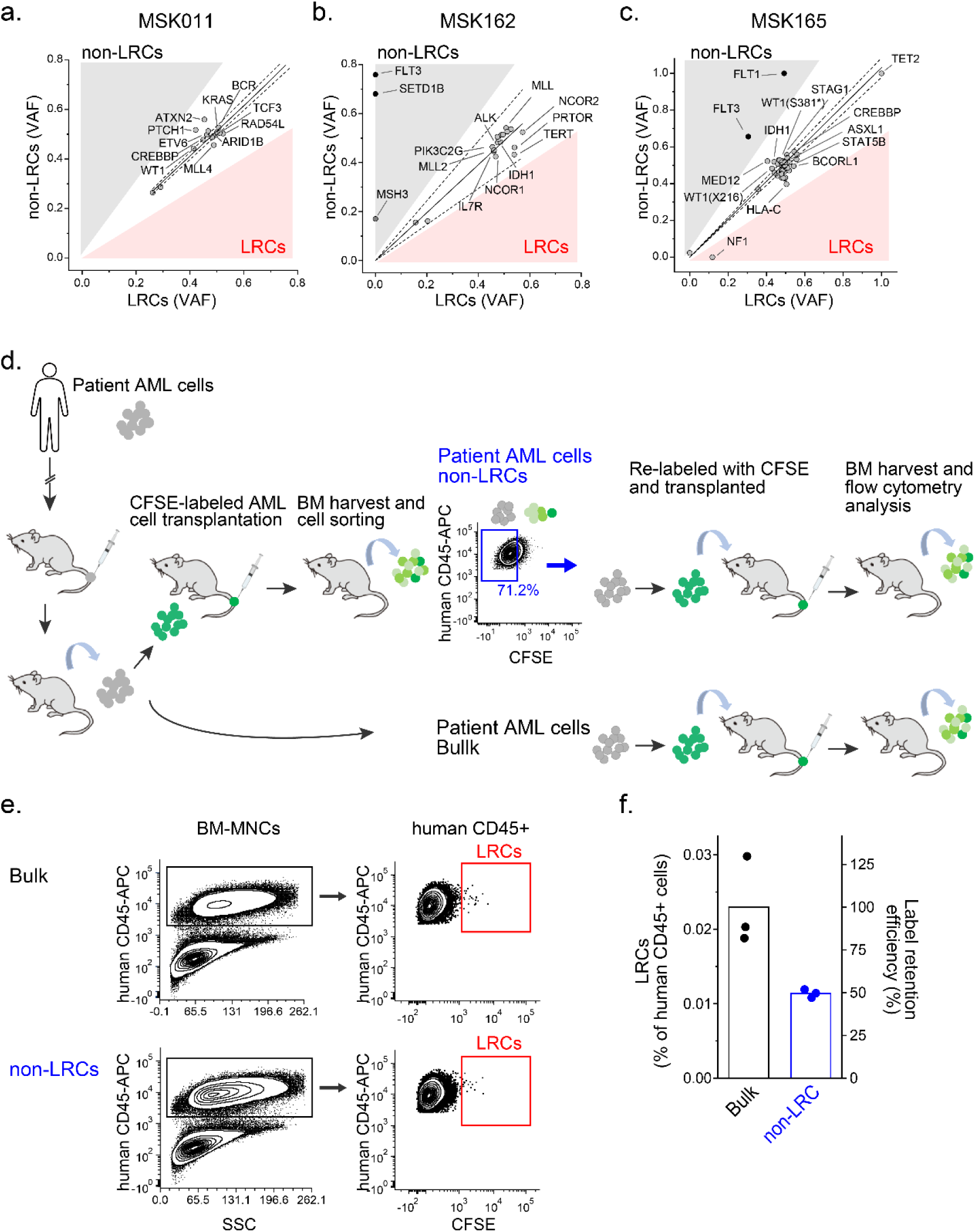
Human patient AML quiescence is reversible evading genetic clonal evolution that maintains disease propagation in serial orthografts in vivo. **a-c.** Comparison of variant allele frequencies (VAF) of genomic DNA sequencing of LRCs (red) versus non-LRCs (gray) in MSK011 (a), MSK162 (b) and MSK165 (c) human patient AMLs, demonstrating several genetic subclones that are enriched in non-LRCs as compared to LRCs. Dashed lines mark 95% confidence intervals. **d.** Experimental design to examine the reversibility of a quiescent state in patient AML cells. Bulk leukemia cells and purified non-LRCs (blue) isolated from the primary recipient mice are re-labeled and sequentially transplanted into secondary recipient mice. **e.** Representative flow cytometry plots of human leukemia cells isolated from bone marrow of secondary recipient mice transplanted with bulk leukemia cells (upper panels) or non-LRCs (lower panels) demonstrate gating strategies to measure LRC frequencies. **f.** Initially proliferating, non-LRC exhibit significant ability to acquire LRC quiescence (blue), and LRC frequencies generated from non-LRCs (blue) are approximately half as compared to parental bulk leukemia cells (black; t-test *p* = 0.0016). Data show LRC frequencies of human leukemia cells (left x-axis) and efficiencies of generated LRCs from non-LRCs relative to bulk cells (left y-axis). Bars represent mean values of measurement of biological triplicates.

To directly investigate whether non-LRCs retain the ability to enter a quiescent state and become LRCs, we conducted re-labeling experiments. Upon isolating non-LRCs from the primary recipient mice transplanted with CFSE-labeled patient AML cells, we re-labeled these cells with CFSE and transplanted them into secondary recipient mice. Simultaneously, parental bulk leukemia cells, which contained ancestors of both LRCs and non-LRCs in primary transplants, were CFSE-labeled and transplanted as controls. We then compared the apparent LRC frequency in the secondary transplants of LRCs to bulk leukemia cells to evaluate the ability to develop LRCs from non-LRCs (Figure 2d). As compared to parental bulk leukemia cells, the initially proliferating non-LRCs were found to acquire LRC quiescence upon secondary transplantation at approximately half the frequency of parental bulk leukemia cells (Figure 2e-f), consistent with other studies^19,20^. Thus, human patient AML cell quiescence is a cell state which is reversibly accessible by leukemia cells. Furthermore, there was indirect evidence that reversibly acquired quiescence involves functional leukemia-initiating cells, as evident by leukemia development in tertiary transplants of non-LRCs (revised Extended Data Figure 3e).

## Distinct promoter-centered chromatin and gene expression dynamics of human LRCs

We reasoned that the reversibility of LRC quiescence can be regulated epigenetically. To elucidate this, we used the recently developed low-input assay for transposase-accessible chromatin with high-throughput sequencing (ATAC-seq) to define LRC-regulated chromatin regions. This identified 777, 3437, and 2036 differentially accessible regions in LRCs of MSK011, MSK162, and MSK165 patient leukemias, respectively (Extended Data Figure 7a-f), including commonly shared loci (Extended Data Figure 7g). Whereas most non-LRC chromatin dynamics were distributed across the genome relatively uniformly, changes in LRC chromatin accessibility were concentrated near promoter regions (Figure 3a-d and Extended Data Figure 7h). We observed specific transcription factor DNA binding sequence motifs in LRC-accessible chromatin regions (Figure 3e-j, Extended Data Figure 8a-h), including those corresponding to ETS, KLF, CCAAT, and AP-1 (e.g. JUN) binding motifs^29–34^, some of which were shared across specimens (Figure 3k).

**Figure 3.**
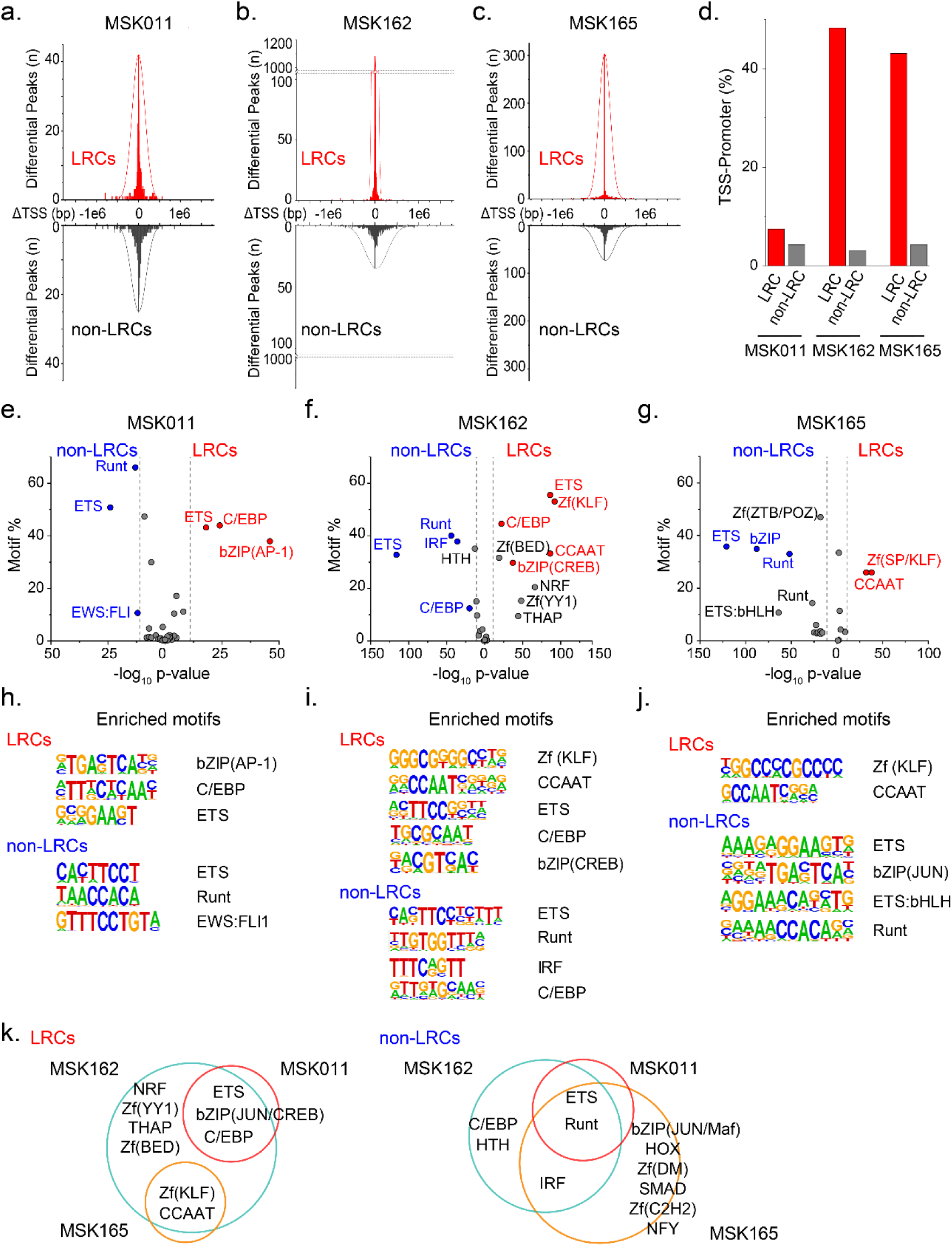
Human patient AML quiescence is associated with promoter-centered chromatin accessibility dynamics. **a-c**. Histograms of differentially accessible chromatin regions in quiescent LRCs (red) as compared to non-LRCs (black) as a function of their distance from transcription start sites (ΔTSS) in MSK011 (a), MSK162 (b), and MSK165 (c) human patient leukemias. **d.** Human quiescent LRCs (red) exhibit increased chromatin accessibility of transcriptional start promoter regions as compared to non-LRCs (gray) in MSK011, MSK162, and MSK165 leukemias (Fisher’s exact test *p* = 6.9 x 10^-2^, 2.2 x 10^-16^, and 2.2 x 10^-16^, respectively). Data represent measurement triplicates. **e-j.** Transcription factor binding sequence motifs enriched in differentially accessible chromatin regions in LRCs (red) as compared to non-LRCs (blue) in MSK011 (e), MSK162 (f) and MSK165 (g), as a function of their statistical significance of enrichment, with specific motif sequences shown (h, i and j, respectively). Dashed lines mark *p*-values of 1.0 x 10^-11^. **k.** Venn diagrams represent overlapping transcription factor binding sequence motifs enriched in differentially accessible chromatin regions in LRCs (left) and non-LRCs (right) across 3 patient AMLs.

Using RNA sequencing, we also identified 749, 567, and 489 significantly differentially LRC-expressed genes in MSK011, MSK162, and MSK165 patient leukemias, respectively, including commonly dysregulated genes (Figure 4a-b, Extended Data Figure 6a-f, Extended Data Table 4 and 5). This included *JUN*, *FLI1*, *ETS1* and *KLF2*, consistent with the sequence motif analysis of differentially accessible LRC chromatin (Figure 4c). We also identified several other potential LRC regulators such as *ZFP36L1* (Figure 4b), which encodes a RNA-binding protein and whose paralog *ZFP36L2* was recently identified as a regulator of AML cell differentiation^35^. Moreover, differentially regulated cell surface protein-coding genes included hematopoietic and leukemia stem cell-related genes such as *ANGPT1*^36,37^, whose expression is associated with inferior patient survival, as assessed in the cohort of 172 AML patients in TCGA AML dataset^38,39^ (log-rank *p* = 0.0038; Extended Data Figure 9g). In all, these findings defined distinct promoter-centered chromatin and gene expression dynamics of human LRCs.

**Figure 4.**
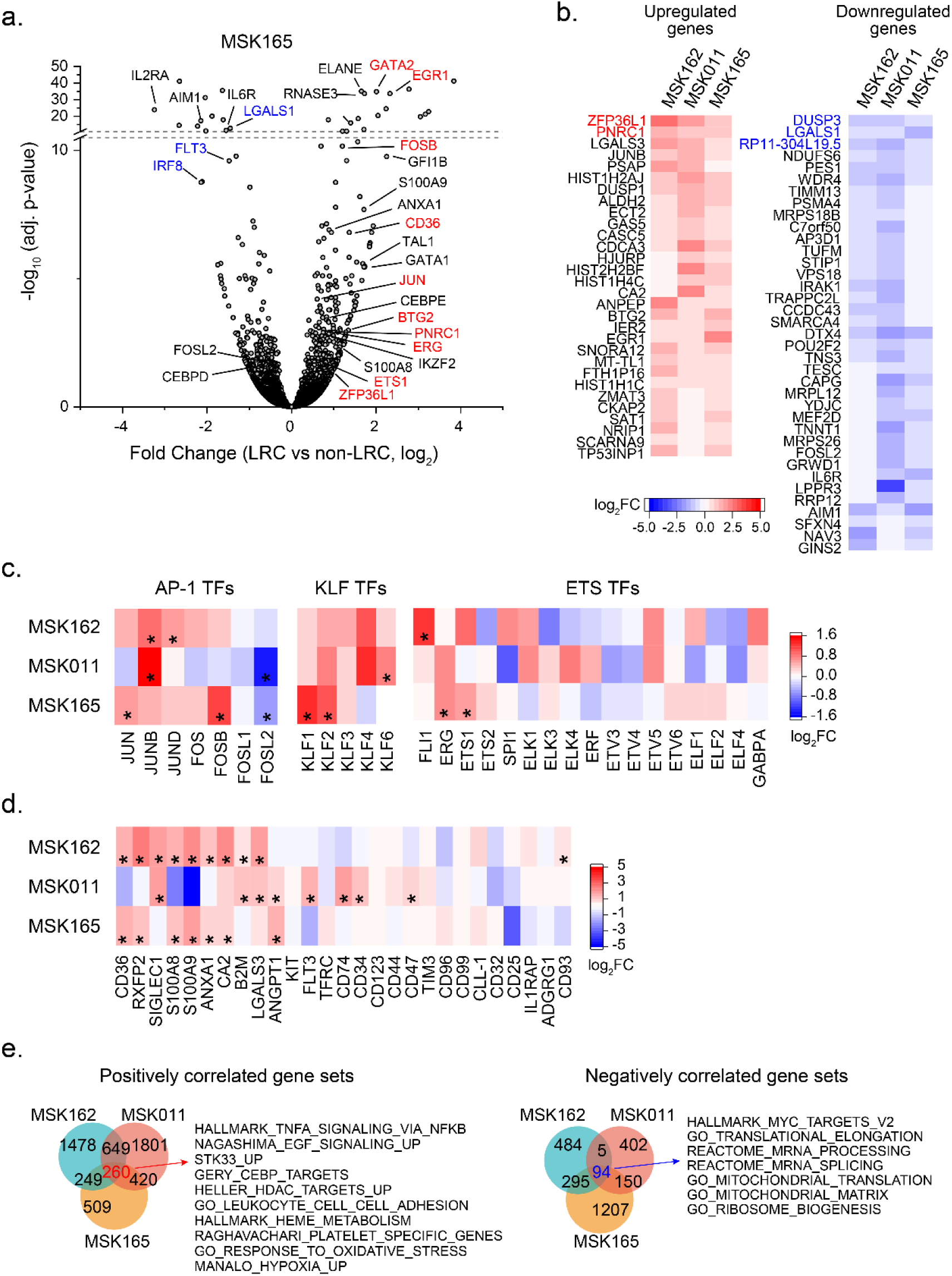
Coherent gene expression dysregulation in quiescent patient AML LRC cells. **a.** Representative gene expression of MSK165 human patient AML cells with statistical significance of measurement triplicates as a function of differential gene expression of LRCs versus non-LRCs. Notable upregulated and downregulated genes are labeled in red and blue, respectively. Results of MSK011 and MSK162 LRC gene expression analysis are described in Extended Data Figure 9d-e. **b.** Heatmaps showing commonly differentially regulated genes between LRCs and non-LRCs which are significantly upregulated (left panel) and downregulated (right panel) in at least 2 human patient leukemias (adjusted *p* < 0.1, fold change > 1.5), with blue to red color gradient representing relative decrease and increase of fold change of gene expression, respectively. **c.** Heatmap showing differential gene expression of specific AP-1, KLF and ETS transcription factors (TFs) between LRCs and non-LRCs in MSK011, MSK162, and MSK165 human patient leukemias with blue to red color gradient representing relative decrease and increase of fold change of gene expression, respectively (* denotes adjusted *p* < 0.1, fold change > 1.5). **d.** Heatmap showing differential gene expression of cell surface proteins between LRCs and non-LRCs in MSK011, MSK162, and MSK165 human patient leukemias with blue to red color gradient representing relative decrease and increase of fold change of gene expression, respectively (* denotes adjusted *p* < 0.1, fold change > 1.5). **e.** Gene set enrichment analysis for differentially expressed genes in MSK011, MSK162, and MSK165 LRCs versus non-LRCs, showing commonly dysregulated expression of genes regulating specific cellular signaling pathways. Venn diagrams show the numbers of significantly positively correlated (left) and negatively correlated (right) gene sets in LRCs (*p* < 0.01, false discovery rate < 0.25). Selected significantly enriched gene sets are listed.

In addition, these data provided an opportunity to determine whether LRC quiescence contributes to therapy resistance and disease persistence in patients with AML. Gene set enrichment analysis (GSEA) of patient LRCs revealed 260 significantly upregulated and 94 significantly depleted gene sets in LRCs (Figure 4e and Extended Data Table 6). For example, we detected downregulation of MYC activity and ribosome biogenesis, similar to prior observations of dormant hematopoietic stem cells^40–43^. We combined GSEA with recent single-cell RNA sequencing (scRNA-seq) of bone marrow cells isolated from AML patients before and after chemotherapy treatment (Figure 5a-b, Extended Data Table 7), leveraging simultaneous detection of differentially expressed and mutant genes to specifically identify specific differential gene expression in leukemia cells^44^. Supervised comparison of genetically varied leukemia cells upon chemotherapy treatment of two different patients, AML329 and AML707B, showed multiple gene sets significantly upregulated and downregulated in residual leukemia cells shared with those specifically expressed in human LRCs (Figure 5b-d). Among them, we identified TNF signaling and hypoxia response pathways for example, which have recently been nominated in AML stem cell pathogenesis^45–48^, and multiple other pathways which provide important hypotheses for future studies (Figure 5b-d). LRCs and residual leukemia cells after chemotherapy also showed similarity to some, but not all, of the previously reported AML stem cell signatures identified by cell surface marker enrichment^49–52^, as also reported recently^45–47^. In all, these findings suggest that gene expression programs induced by LRC quiescence contribute to therapy resistance and leukemia persistence in patients.

**Figure 5.**
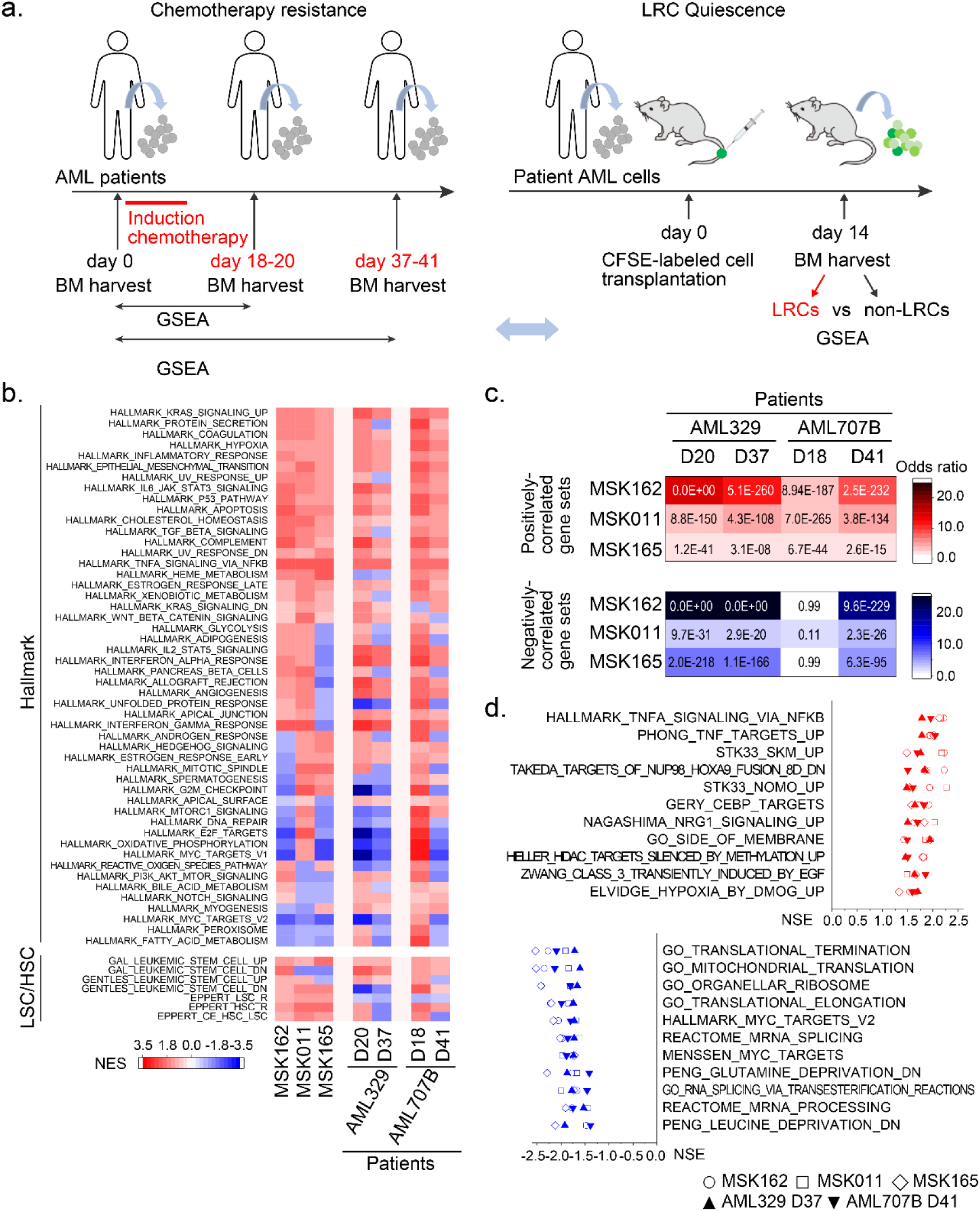
Shared gene expression dynamics associated with chemotherapy resistance and quiescence in diverse human AML patient specimens. **a.** Schematic of comparative gene expression analyses of AML cells isolated from the bone marrow (BM) of patients before and after treatment with induction chemotherapy (left) on indicated days^37^, and LRC quiescence of CFSE label retention in mouse orthografts (right). **b.** Heatmaps showing specific enrichment of distinct gene sets between LRCs in MSK162, MSK011, and MSK165 specimens and chemotherapy-resistant cells in AML329 and AML707B specimens analyzed after induction chemotherapy treatment. Red to blue color gradient represents positive and negative normalized enrichment scores (NES), respectively. **c.** Heatmaps of significance of similarity in gene expression between MSK162, MSK011, and MSK165 LRCs and chemotherapy-resistant cells in AML329 and AML707B patient specimens (numbers indicate hypergeometric test *p* values). White to red and white to blue color gradients represent positive and negative odds ratios, respectively. **d.** Commonly dysregulated gene sets shared by both LRCs and residual leukemia cells after chemotherapy were listed. For example, TNF-alpha and NRG1 signaling pathways were positively correlated, whereas MYC, RNA processing and mitochondrial biogenesis pathways were negatively correlated in both LRCs and residual leukemia cells after chemotherapy.

## AML LRC quiescence is controlled by a distinct transcription factor network

We sought to develop a functional genetic system to identify regulators of human patient LRC quiescence. To enable this, we designed a lentiviral vector for the production of ultrahigh virus titers necessary for the transduction of primary human leukemia cells, encoding mCherry fluorescent protein and doxycycline-inducible cDNAs marked by specific barcode sequences (Figure 6a, Extended Data Table 8). We selected 20 transcription factors with differential expression in diverse LRCs with concordant markers of chromatin motif accessibility at LRC-regulated genes (Figures 3-4, Extended Data Table 9). We confirmed appropriate doxycycline-inducible cDNA expression by Western immunoblotting (Extended Data Figure 10a), prepared equimolar cDNA plasmid pools (Extended Data Figure 10b), transduced primary human MSK162 patient leukemia cells to achieve single-copy cDNA integration as assessed by mCherry expression by fluorescence-activated cell scanning (<20% cells; Extended Data Figure 10c), and confirmed their stable representation of at least 5,000 cells/cDNA by quantitative DNA barcode sequencing after primary transplantation in NSG mice (10^6^ cells/mouse) in the absence of doxycycline (Extended Data Figure 10d-e).

**Figure 6.**
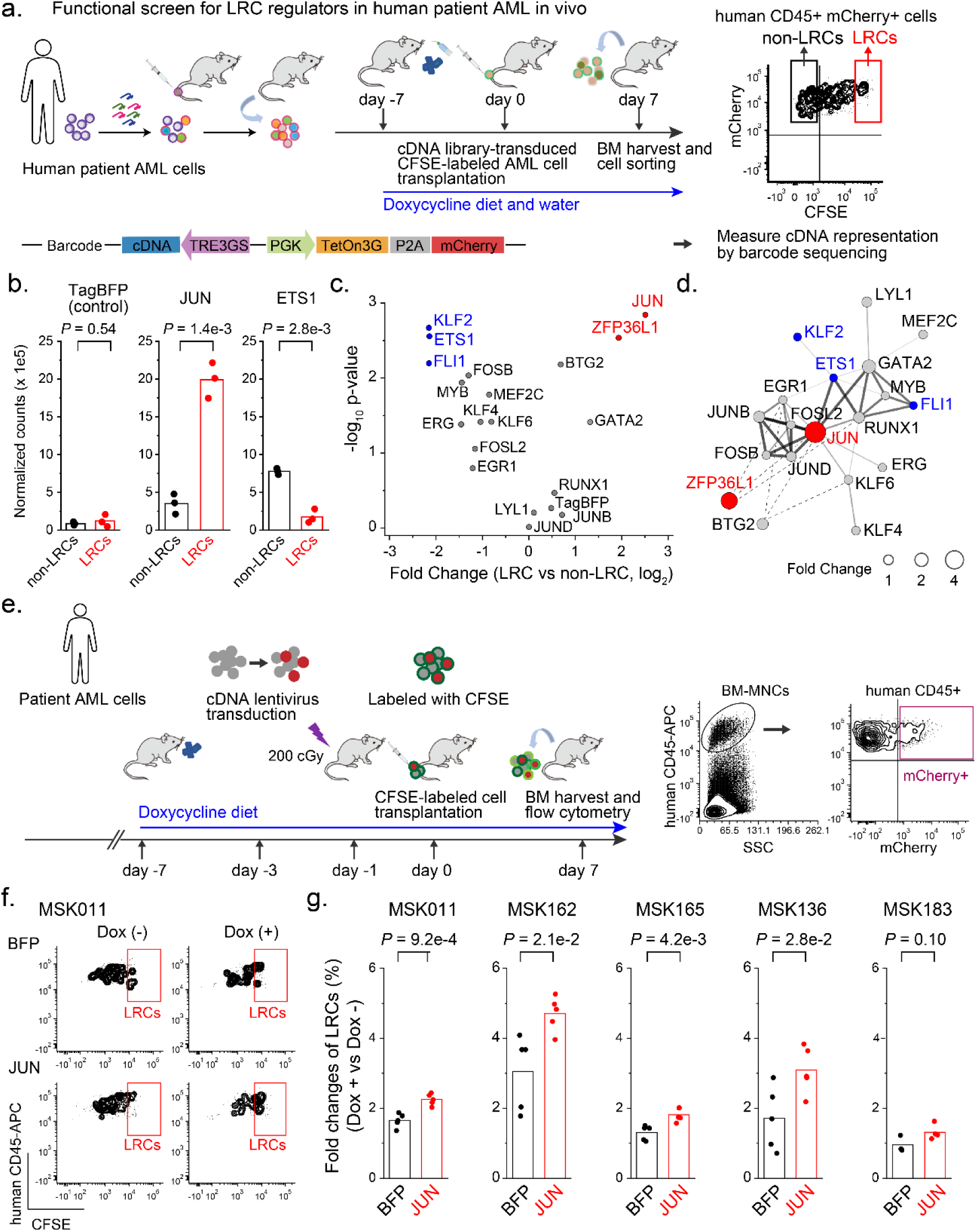
AML stem cell quiescence is controlled by a distinct transcription factor network. **a.** Experimental design to identify regulatory factors controlling AML stem cell quiescence using lentiviral transduction of doxycycline-inducible cDNA library in human patient AML cells, followed by CFSE labeling to isolate quiescent LRCs (red) and proliferating non-LRCs (black). Days of induction of cDNA expression using doxycycline treatment are indicated. Each lentiviral doxycycline-inducible cDNA vector includes a specific barcode sequence, enabling quantitative identification by DNA sequencing (bottom). **b.** Enforced expression of *JUN* (center panel) or *ETS1* (right panel) in MSK162 patient AML specimen induces or depletes quiescent LRCs (red) as compared to non-LRCs (black), respectively (two-tailed Welch’s t-test *p* = 1.4 x 10-3 and 2.8 x 10-3, respectively). Enforced TagBFP expression serves as negative control (left panel, two-tailed Welch’s t-test *p* = 0.54). Bars represent mean values of normalized read counts of biological triplicates. **c.** Volcano plot showing genes whose enforced expression induces (red) or depletes (blue) quiescent LRCs in MSK162 patient AML specimen. TagBFP serves as negative control. **d.** Schematic of the LRC regulatory interaction network. Red and blue circles denote LRC activators and repressors, respectively, with circle size proportional to effect size. Solid lines indicate physical interactions with line thickness corresponding to STRING confidence scores. **e.** Experimental design to investigate the function of *JUN* as a regulator for LRC quiescence in diverse patient AMLs. Patient AML cells are transduced with mCherry-expressing doxycycline-inducible *JUN* or TagBFP lentivirus vectors, and after labeled with CFSE, transplanted into NSG mice with or without doxycycline diet *in vivo* (left panel). Representative flow cytometry plots show gating strategies to analyze LRC frequencies in *JUN* or TagBFP-transduced patient leukemia cells by gating on human CD45-positive, mCherry-expressing cells (right panel). **f.** Representative flow cytometry plots exhibit LRC distribution of CD45-positive, mCherry-expressing MSK011 patient leukemia cells, *JUN* (lower panels) versus TagBFP (upper panels) with (right panels) or without doxycycline (left panels). **g.** LRC frequencies of human CD45-positive, mCherry-expressing cells are measured in five different patient AMLs (refer to Extended Data Figure 10g). Dot plots demonstrate fold changes of LRC frequencies in cells with versus without doxycycline induction in *JUN* (red) or TagBFP (black) transduced cells. LRC frequencies are significantly increased in *JUN*-transduced cells upon doxycycline induction compared to TagBFP-transduced cells in 4 of 5 patient AML cells, exhibiting that enforced JUN expression induces LRC quiescence. (t-test *p* = 9.2 x 10^-4^, 2.1 x 10^-2^, 4.2 x 10^-3^, 2.8 x 10^-2^ and 0.01 for MSK011, MSK162, MSK165, MSK136 and MSK183, respectively). Bars represent mean values of measurement of 3 to 5 biological replicates.

To identify regulators of human LRC quiescence, we CFSE-labeled MSK162 patient leukemia cells genetically-modified with the LRC regulator cDNA library, and transplanted them into NSG mice treated with doxycycline to enforce specific cDNA expression in individual leukemia cells (Figure 6a and Extended Data Figure 10f). Analysis of abundance of cDNA barcodes in LRCs versus non-LRCs revealed positive and negative regulators of LRC quiescence, in contrast to control TagBFP which showed no significant effect (t-test *p* = 0.54; Figure 6b-c). For example, enforced expression of *JUN* or *ZFP36L1* caused a nearly 4-fold increase in LRCs (t-test *p* = 1.4 x 10^-3^ and 2.9 x 10^-3^, respectively), whereas enforced expression of *ETS1*, *FLI1* or *KLF2* suppressed LRC retention by more than 4-fold (t-test *p* = 2.8 x 10^-3^, 6.4 x 10^-3^, and 2.1 x 10^-3^, respectively; Figure 6b-c). Since many of the identified LRC regulators can interact physically^53^, they may comprise an integrated LRC regulatory transcription factor network, which is at least in part centered on JUN (Figure 6d).

## Fine-tuned expression of LRC regulators is required for leukemia progression

The cooperative regulation of LRC quiescence by distinct transcription factors including JUN suggests that specific levels of expression and activity are required for LRC quiescence and leukemia stem cell function. To investigate the function of *JUN* as a LRC regulator in diverse patient leukemias, we engineered five different patient AML cells to enforce JUN expression using the mCherry-expressing doxycycline-inducible lentivirus vector system, and after labeling with CFSE, transplanted them into NSG mice with or without doxycycline diet *in vivo* to induce expression of JUN versus TagBFP control (Figure 6e). We used fluorescence-activated cell scanning to identify human CD45-positive transduced mCherry-expressing cells and then quantified LRC distribution in doxycycline-induced JUN or TagBFP-expressing cells (Figure 6e). This analysis revealed that LRCs were significantly increased in JUN-expressing cells as compared to control TagBFP upon induction with doxycycline in four out of five patient AMLs studied (t-test *p* = 9.2 x 10^-4^, 2.1 x 10^-2^, 4.2 x 10^-3^, and 2.8 x 10^-2^ for MSK011, MSK162, MSK165, and MSK136, respectively; Figure 6f-g and Extended Data Figure 10g). Thus, enforced JUN expression is sufficient to promote LRC quiescence in multiple diverse patient leukemias.

Compelled by these findings, we sought to determine whether JUN-induced LRC quiescence is also associated with altered leukemia stem cell function and disease progression *in vivo*. Thus, we performed competitive transplantation experiments of JUN-overexpressing patient leukemia cells. We separately engineered doxycycline-inducible JUN or TagBFP-expressing MSK165, MSK011, and MSK162 cells, isolated transduced cells by their mCherry expression using fluorescence-activated cell sorting, and transplanted equal cell numbers of JUN and TagBFP-transduced cells into NSG mice with or without doxycycline diet *in vivo* (Figure 7a). Recipient mice developed disease with variable latency and penetrance with and without doxycycline (Figure 7b and Extended Data Figure 11a-b). However, genomic DNA analysis of engrafted leukemia cells revealed that doxycycline induction was associated with the relative loss of JUN-expressing clones (decrease from 29% to 0%, 33% to 0%, and 63% to 33% for MSK165, MSK011, and MSK162 leukemias, respectively; Figure 7c-e). Thus, in addition to promoting LRC quiescence, enforced JUN expression also impairs leukemia progression, presumably via effects on leukemia stem cell engraftment and/or proliferation.

**Figure 7.**
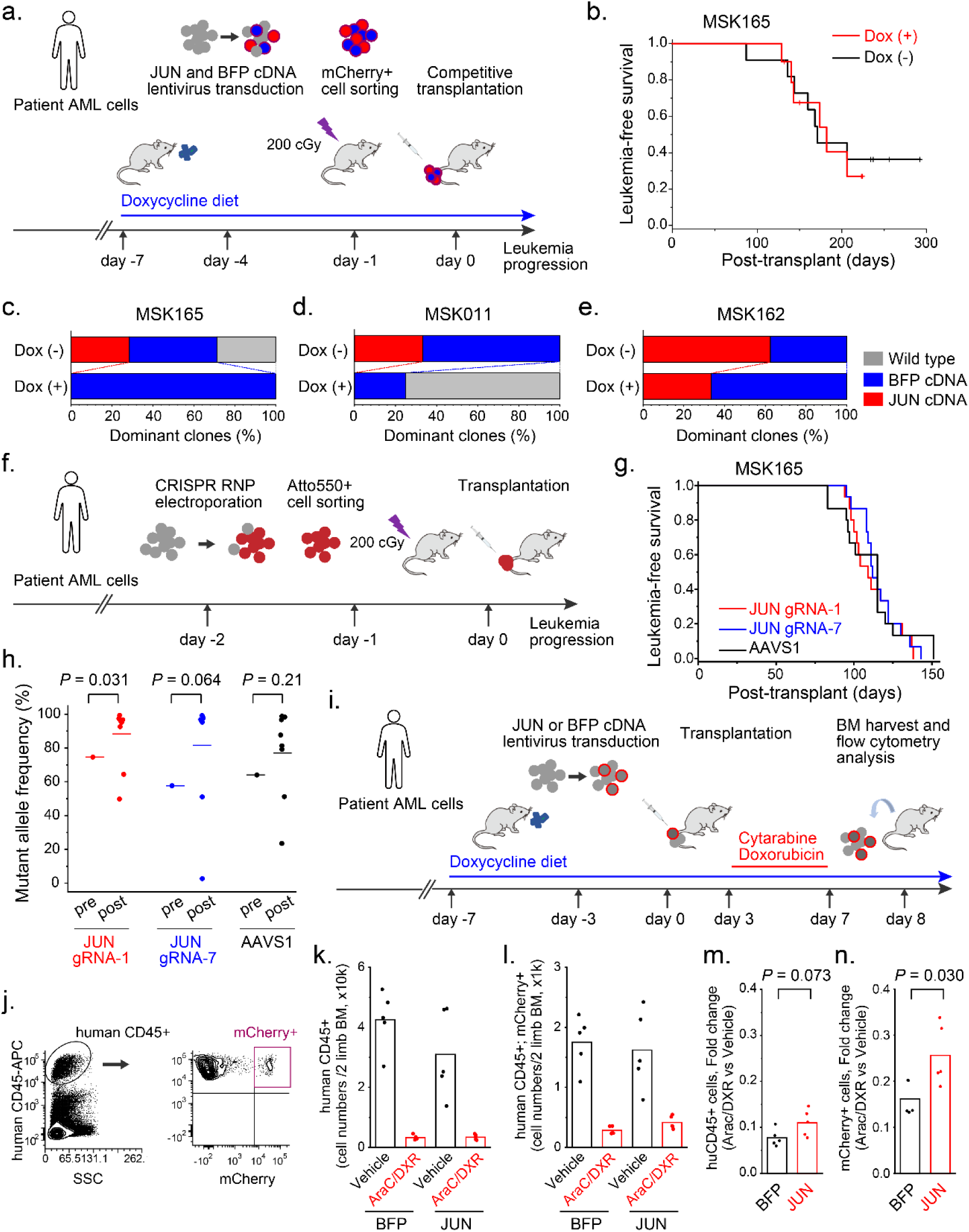
Fine-tuned expression of LRC regulators is required for leukemia progression and chemotherapy resistance. **a.** Experimental design for competitive transplantation of JUN-overexpressing-patient leukemia cells. Doxycycline-inducible *JUN* or TagBFP-transduced, mCherry-expressing patient leukemia cells are separately engineered and isolated by fluorescence-activated cell sorting. Equal cell numbers of *JUN* and TagBFP-transduced cells are transplanted into NSG mice with or without doxycycline diet *in vivo*. **b.** Leukemia-free survival of mice transplanted with a mixture of *JUN* and TagBFP-transduced MSK165 patient leukemia cells with or without doxycycline diet is shown (25,000 cells/mouse, 11 mice for each group). (refer to Extended Data Figure 11a-b for MSK011 and MSK162) **c-e.** Genomic DNA of engrafted patient leukemia cells is analyzed to determine the dominant clones of propagated leukemia cells in each mouse (refer to Extended Data Figure 11c-e). Stacked bar charts represent the proportion of dominant clones in each group with (bottom bar) or without (upper bar) doxycycline induction in MSK165 (left panel), MSK011 (center panel) and MSK162 (right panel). *JUN*-transduced clones are relatively reduced upon doxycycline induction, exhibiting enforced JUN expression impairs leukemia progression (28.6% to 0%%, 33.3% to 0%, and 62.5% to 33.3% for MSK165, MSK011, and MSK162 leukemias, respectively). **f.** Experimental design to investigate leukemia progressing property of *JUN*-knockout patient leukemia cells, which are generated using Cas9 crRNA;ATT0550-labeled-tracrRNA RNP electroporation, isolated by fluorescence-activated cell sorting, and transplanted into NSG mice. **g.** Leukemia-free survival of mice transplanted with *JUN*-knockout or control *AAVS1*-targeted MSK165 patient leukemia cells (30,000 cells/mouse, 15 mice for each group), where *JUN*-knockout cells propagate leukemia with similar kinetics as control *AAVS1*-targeted cells (log-rank *p* = 0.74 and 0.85 for *JUN* gRNA-1 and gRNA-7 versus *AAVS1*, respectively). (refer to Extended Data Figure 11b and 11d for MSK011 and MSK162). **h.** Mutant allele frequencies are analyzed using the Tracing of Indels by Decomposition (TIDE) method before and after transplantation of *JUN*-knockout or control *AAVS1*-targeted MSK165 patient leukemia cells. *JUN*-knockout clones are enriched upon leukemia progression *in vivo*, whereas *AAVS1*-targeted cells are not (t-test *p* = 0.031, 0.064 and 0.21 for *JUN* gRNA-1, gRNA-7 and *AAVS1*, respectively). **i.** Experimental design to investigate whether enforced JUN expression confers chemotherapy resistance on patient leukemia cells. mCherry-expressing, doxycycline-inducible JUN or TagBFP-transduced MSK011 patient leukemia cells are labeled with CFSE and transplanted into NSG mice under doxycycline diet *in vivo*, followed by combined AraC and DXR chemotherapy *in vivo*. **j.** Representative flow cytometry plots to analyze bone marrow human leukemia cells isolated from mice transplanted with MSK011 patient leukemia cells containing mCherry-expressing, JUN or TagBFP-transduced cells. **k-l.** Combined AraC and DXR chemotherapy treatment reduces total human CD45-positive (j) and mCherry expressing (k) human leukemia cell numbers in mouse bone marrow, regardless of JUN or TagBFP transduction (t-test *p* = 7.9 x 10^-4^ and 1.3 x 10^-2^ for CD45-positive cell numbers of TagBFP and JUN; 1.5 x 10^-3^ and 1.4 x 10^-2^ for mCherry-positive cell numbers of TagBFP and JUN, respectively). Bars represent mean values of 5 biological replicates. **m-n.** There is no significant difference in fold reduction of total human leukemia cell numbers in AraC/DXR-treated mice relative to vehicle-treated mice between JUN versus TagBFP transduction group (t-test *p* = 0.074; l), whereas mCherry-positive JUN-expressing cells exhibit increased resistance to AraC/DXR treatment compared to mCherry-positive TagBFP-expressing cells (t-test *p* = 0.030). Bars represent mean values of 5 biological replicates.

JUN is an AP-1 family transcription factor and its enforced expression may regulate diverse AP-1 target genes by heterodimerization with other AP-1 transcription factors. To determine the requirements of JUN for LRC quiescence and leukemia stem cell function and propagation, we used CRISPR genome editing to engineer loss-of-function mutations of *JUN* in patient leukemia cells (Figure 7f and Extended Data Figure 12b). We identified two specific gRNAs that produced bi-allelic loss-of-function mutations of *JUN* and loss of measurable JUN protein expression upon electroporation of Cas9 crRNA:tracrRNA ribonucleoprotein (RNP) complex (Extended Data Figures 12a and 13a). By using tracrRNAs labeled with the ATTO550 fluorophore, we used fluorescence-activated cell sorting to isolate electroporated cells before CFSE labeling and transplantation into NSG mice.

Using this approach, we transplanted equal numbers of CFSE-labeled *JUN*-knockout patient AML cells, which did not show substantial differences in LRC quiescence as compared to control *AAVS1*-targeted cells (Extended Data Figure 12c). Similarly, we did not observe significant differences in leukemia-free survival of mice transplanted with *JUN*-knockout patient AML cells as compared to control *AAVS1*-targeted cells (Figure 7g, Extended Data Figure 13b and 13d). Although with some variability among individual mice, genomic DNA analysis of leukemia cell mutant allele frequencies before and after transplantation using the tracking of indels by decomposition (TIDE) method^54^ showed significant enrichment of *JUN*-knockout but not *AAVS1*-targeted clones upon leukemia progression *in vivo* as compared to cells before transplantation (t-test *p* = 0.031 and 0.064 versus 0.21 for *JUN* gRNA-1 and gRNA-7 versus *AAVS1*, respectively; Figure 7h). Similar results were seen in limiting dilution transplant experiments (t-test *p* = 0.019 and 0.32 for MSK011 *JUN* gRNA-1 versus *AAVS1*, respectively; Extended Data Figure 13b-e). Thus, loss of JUN can promote leukemia progression, presumably via effects on leukemia stem cell engraftment and/or proliferation. In all, these results indicate that fine-tuned expression of LRC regulators such as *JUN* can control human leukemia quiescence, stem cell function and disease progression.

## AP-1 activity is required for chemotherapy resistance

Our findings indicate that LRCs comprise a chemotherapy-resistant reservoir. To investigate whether JUN activity can contribute to chemotherapy resistance, we transplanted patient AML cells containing doxycycline-inducible *JUN* or *TagBFP*-transduced cells into NSG mice, and treated them with the combined AraC and DXR treatment regimen developed to model induction chemotherapy used clinically for human patients^24^, combined with doxycycline diet *in vivo* (Figure 7i-j). As expected, total mouse bone marrow mononuclear cells and human CD45+ leukemia cells were markedly reduced in AraC/DXR-treated mice as compared to vehicle control treated mice, regardless of expression of JUN versus TagBFP (Figure 7k-l and Extended Data Figure 11g). However, transduced JUN-expressing cells were significantly enriched upon chemotherapy treatment as compared to TagBFP-transduced cells (t-test *p* = 0.029; Figure 8m-n). Thus, in addition to regulating LRC quiescence and leukemia stem cell function and propagation, enforced JUN expression confers chemotherapy resistance *in vivo*.

**Figure 8.**
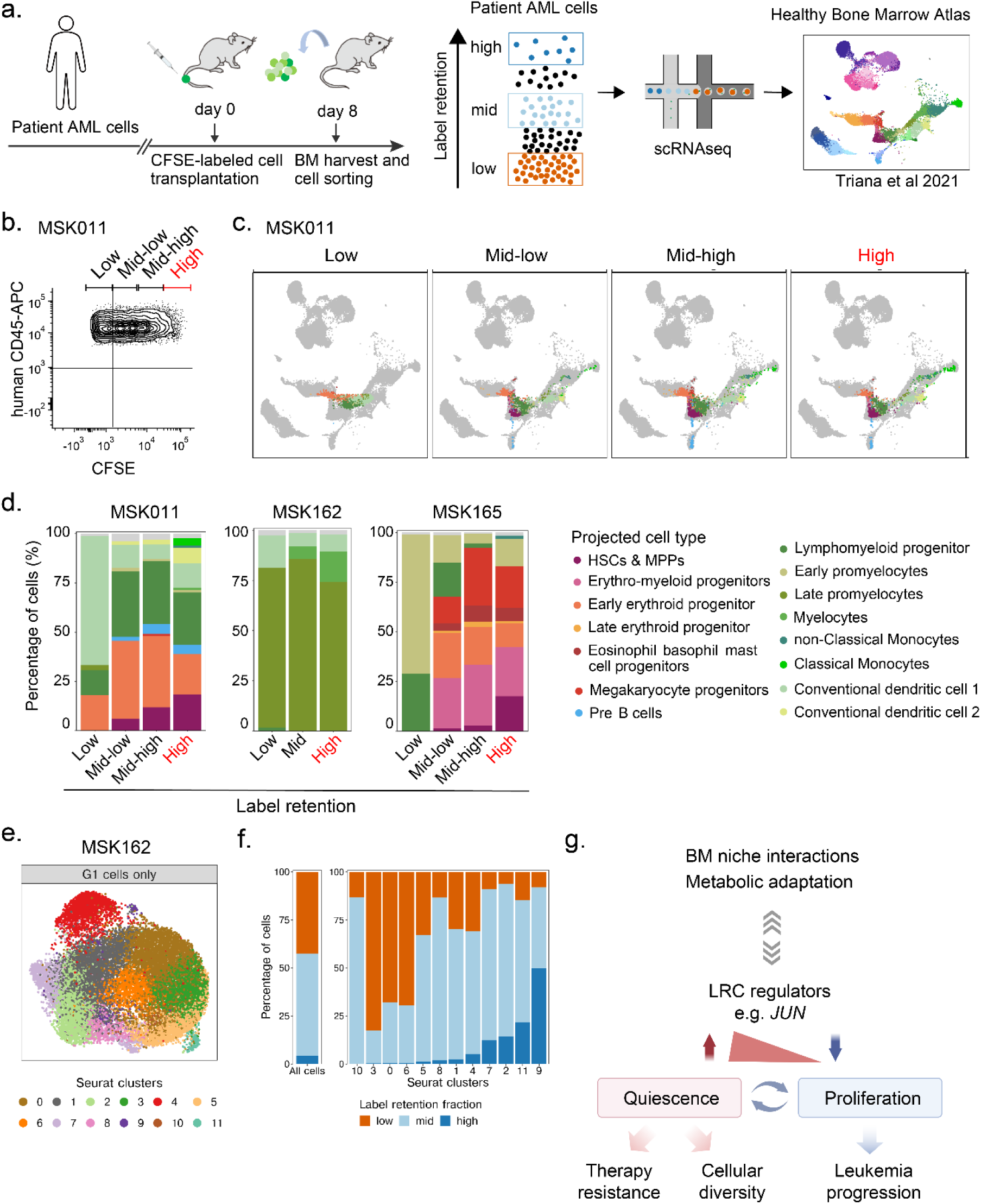
Single-cell analysis of LRC quiescence reveals shared and distinct gene expression programs. **a.** Experimental design for single-cell RNA sequencing of patient LRCs versus non-LRCs performed using three different patient AMLs. Human leukemia cells isolated from mice transplanted with CFSE-labeled patient leukemia cells are separately collected based on CFSE-label retention levels of high, middle and low using fluorescence-activated cell sorting. Gene expression profiles are mapped to a reference atlas of healthy human bone marrow hematopoiesis. **b.** Representative flow cytometry plots to isolate human patient leukemia cells based on the levels of CFSE-label retention in MSK011. (Flow cytometry plots of MSK162 and MSK165 are described in Extended Data Figure 16c and 16e.) **c.** Uniform manifold approximation and projection (uMAP) for high, middle-high, middle-low and low-label retaining cells of MSK011 exhibit that higher label retaining cell fractions are comprised of more diverse cell types, including HSC- and MPP-like cells. (uMAPs for MSK162 and MSK165 are described in Extended Data Figure 16d and 16f.) **d.** Stacked bar charts represent projected cell type composition of indicated label-retaining cell fractions of MSK011 (left panel), MSK162 (center panel) and MSK165 (right panel) patient leukemia cells. High label-retaining cells exhibit shared and distinct gene expression programs among three different patient AMLs. **e-f.** uMAP depicts unsupervised clustering of G1 MSK162 cells, which identifies 12 distinct clusters (e). Stacked bar charts represent cell status of label retention in each 12 cluster (h), exhibiting cells with high label retention are enriched in cluster 9 and 11. **g.** Schematic of the mechanisms controlling leukemia cell quiescence and progression.

To explore whether this applies to human patients, we analyzed expression of LRC regulatory factors in individual AML cells in diverse human patients before and after clinical chemotherapy treatment (Extended Data Figure 14a-d, Extended Data Table 7)^44^. We found *JUN* and/or its AP-1 cofactor *FOS* to be highly expressed in persistent AML cells upon induction chemotherapy in patients AML707B (*JUN* and *FOS*) and AML329 (*FOS*) (*p* = 6.2 x 10^-3^ and 8.6 x 10^-6^ for AML 707B and 1.5 x 10^-8^ for AML 329, respectively; Extended Data Figure 14a-d). AML870 also showed *JUN* upregulation in persistent chemotherapy-resistant leukemias cells, especially in those with features of hematopoietic stem (HSC-like) gene expression (Fold Change = 3.5, *p* = 0.0051 for AML707B; Fold change = 5.0, *p* = 0.13 for AML870; Extended Data Figure 15a-b). High *JUN* expression in HSC-like persistent chemotherapy-resistant cells was also maintained in patients with monocytic AML (AML420B and AML556; Extended Data Figure 15d-e). Interestingly, in patient AML707B, in addition to AP-1 TFs, other LRC regulatory factors, such as *GATA2* and *ZFP36L1*, were also upregulated in residual leukemia cells after chemotherapy (Extended Data Figure 14c), consistent with their observed functions in LRC quiescence (Figure 6c). These factors were also preferentially co-expressed in individual leukemia cells, consistent with their cooperative regulation of cellular quiescence and drug resistance (Extended Data Figure 14e). In all, these results suggest that quiescence regulatory factors, including AP-1 factors specifically, can control chemotherapy resistance, quiescence, and ultimate leukemia relapse in patients.

## Single-cell analysis of LRC quiescence reveals shared and distinct gene expression programs

Is cell quiescence of various human leukemias associated with uniform or varied cell states? To explore this, we performed single-cell RNA sequencing (scRNA-seq) of human patient LRCs versus non-LRCs based on high, middle, and low levels of CFSE label retention using three diverse patient AMLs (Figure 8a). Human leukemia cells harvested from mouse bone marrow were separated into 3-4 fractions based on CFSE signal intensity using fluorescence-activated cell sorting (Figure 8b, Extended Data Figure 16c, and 16e), and analyzed using droplet-based Chromium 10x Genomics single-cell gene expression profiling. We obtained between 182-16400 cells per fraction from each patient leukemia after stringent quality control filters (Extended Data Figure 16a). As expected, cells with low label retention had higher S and G2M cell cycle phase Seurat gene expression scores^55^ as compared to cells with middle and high label retention (Extended Data Figure 16b).

To define cell states, we mapped the observed leukemia cell gene expression to a reference atlas of human bone marrow hematopoiesis^56,57^ (Figure 8a). This analysis revealed that LRCs were comprised of cells with varied transcriptional states, but distinct from non-LRCs (Figure 8c-d, and Extended Data Figure 16e-f). MSK011 and MSK165 contained cells with HSC- or MPP-like gene expression programs, which were enriched for high-LRCs, suggesting that some LRCs share transcriptional features of primitive LSCs (p-LSCs), also identified in a recent study^58^. On the other hand, most cells from MSK162 exhibited myeloid differentiated features, and no cells displayed HSC- or MPP-like gene expression program.

To further define gene expression features of quiescent LRCs, we carried out unsupervised clustering of G1-phase MSK162 cells. This identified twelve distinct clusters, with clusters 9 and 11 preferentially enriched in high label-retaining LRCs (Figure 8e-h). Cluster 11 cells expressed high levels of *BCAT1* and exhibited a high monocytic leukemic stem cell score (mLSC score) which was recently found to be associated with venetoclax-resistant monocytic leukemic stem cells (Extended Data Figure 16g)^58^. Thus, LRC quiescence was associated with distinct but varied transcriptional cell states among different patient leukemias.

We also analyzed the observed scRNA-seq cell states using single-cell regulatory network inference and clustering (SCENIC) gene regulatory network inference^59,60^. This revealed specific regulons, including distinct apparent activities of AP-1 and ETS transcription factors, such as FOS in MSK011, JUND, RUNX1, and NR3C1 in MSK162, ELF2 and ZNF148 in MSK165 (Extended Data Figure 17a, 17c, and 17e). Thus, quiescent LRCs share common and distinct gene expression programs in diverse human patient leukemias, including distinct AP-1 and ETS transcription factor networks associated with specific hematopoietic developmental lineages.

## Discussion

Cancer relapse occurs in the majority of patients after chemotherapy and frequently signifies incurable disease. In acute myeloid leukemia, a common blood cancer that affects both children and adults, disease relapse is largely due to the persistence of leukemia initiating stem cells. Here, we implemented label tracing functional genomic techniques to human patient AML specimens. This unbiased approach enables molecular and functional analyses of diverse leukemia initiating and quiescent LRC stem cells that evade detection by currently known cell surface markers, as also proposed by others^19,20,61^.

Recent observations of altered protein homeostasis in AML and hematopoietic stem cells^62–64^, as at least in part induced by dysregulation of mRNA translation^65^, may explain the specific ability of chemical protein label retention to identify functional leukemia initiating and disease persisting cells. Future studies will be important to define the molecular mechanisms of LRC proteome quiescence.

Importantly, our findings explain the frequent and seemingly paradoxical observations of how AML relapse in patients can occur both from rare leukemia stem cells as well as immunophenotypically committed or differentiated subclones^8,66^. Our findings also indicate that drug resistance is a consequence of dormancy^67^. Prior work by Bhatia and colleagues and Melnick and colleagues have implicated chemotherapy, and cytarabine in particular, as an inducer of altered and senescent-like state responsible for AML relapse^45,46^. Our work indicates that many of the features of quiescence and dormancy observed in this state are already accessible to leukemia cells as LRCs. The similarity of gene expression profiles of LRCs, combined with the apparent kinetics of their growth upon transplantation in mice, suggest that this population exists before the onset of therapy and not because of treatment.

Do LRCs occupy different bone marrow niches from normal HSCs? Upregulation of cell adherent molecules or hypoxic response in LRCs suggest that LRCs may function in the context of a distinct bone marrow niche (Figure 4e), as previously observed for endosteal cells^68^. The interactions of LRCs with tissue niches might represent new therapeutic targets. For example, our work helps to corroborate distinct metabolic requirements of leukemia stem cells^69^, including the function of fatty acid oxidation in the promotion of LRC quiescence and therapy resistance, as evident from the high LRC expression of the fatty acid receptor CD36^47,70^ (Figure 4a, 4d, and Extended Data Table 5). LRC-expressed molecules, such as CD36, which may be dispensable for healthy hematopoietic stem and progenitor cells, represent compelling therapeutic targets to selectively eradicate LRCs.

The organization of the LRC transcription factor network suggests that acquisition of LRC quiescence may be accomplished by the coordinated dysregulation of cell differentiation and dormancy. The apparently divergent functions of JUN and ETS1 in controlling LRC quiescence, while both being required for normal hematopoietic stem and progenitor cell functions (Expanded Data Figure 9), suggest that the LRC transcription factor network is aberrantly organized, as recently also observed for other aberrantly organized transcription factor complexes in genetically diverse human leukemias^71,72^.

In addition, this transcription factor network appears dynamic, as enforced JUN expression promotes quiescence and chemotherapy resistance while inhibiting leukemia proliferation, and JUN downregulation enhances leukemia stem cell function and disease progression (Figure 8g). Human quiescence appears to have shared features among molecularly diverse AML subtypes (Figure 4e), but its transcriptional heterogeneity will need further studies (Figure 8d and Extended Data Figure 17), particularly for the investigation of new therapeutic strategies to eradicate quiescent cells that are responsible for disease relapse. While our studies indicate that LRC quiescence is reversible, sparing genetic competition that maintains its epigenetic inheritance, clonal genetic evolution is also an important part of leukemia development in patients, given its contributions to pre-leukemia clonal hematopoiesis and leukemia therapy resistance^26^.

Finally, our study is limited by transplantation in immunodeficient mice, with its distinct cytokine and cellular environments, which may incompletely model human physiology and thereby obscure additional mechanisms of human leukemia stem cells. We also cannot exclude the possibility of LRC genetic mutations that are not captured by target gene sequencing, even though we targeted most known recurrently mutated genes in hematologic malignancies^25^. We anticipate that similar studies of pre-malignant and cancer cells will provide essential insights into the mechanisms of cancer initiation, evolution, and therapeutic targeting. This should lead to new therapeutic strategies aimed at restoring normal cell development and therapeutic control of cancer cell quiescence.

## Methods

### Human AML specimens

Primary AML specimens were obtained from the bone marrow of patients upon obtaining written informed consent and approval by the Institutional Review Board of Memorial Sloan Kettering Cancer Center, MD Anderson Cancer Center and the Children’s Oncology Group in accordance with the Declaration of Helsinki.

### CFSE-labeling

5(6)-Carboxyfluorescein diacetate N-hydroxysuccinimidyl ester (CFSE, Abcam) was dissolved in DMSO at 5 mM and stored at -20 °C. To prepare CFSE-labeled cells, cells were washed in PBS once and incubated with 1 *μ*M CFSE in PBS supplemented with 1% FBS at 37 °C for 5 minutes, unless otherwise indicated. After the reaction was quenched by adding 15 ml of Iscove’s Modified Dulbecco’s Medium (IMDM) supplemented with 15% FBS, cells were washed twice with IMDM supplemented with 15% FBS using centrifugation.

### Detection of fluorescently-labeled proteins

Cells were lysed in RIPA buffer (140 mM NaCl, 0.4% SDS, 0.1% sodium deoxycholate, 1% Triton X-100 and 20 mM Tris-HCl) with sonication using the Covaris S220 instrument (Covaris). Lysates were cleared by centrifugation at 14,000 x g for 15 minutes at 4 °C, and the collected lysates were denatured in Laemmli sample buffer supplemented with 50 mM DTT at 95 °C for 5 minutes. Lysates were separated by sodium dodecyl sulfate polyacrylamide gels electrophoresis (SDS-PAGE, Novex) at 120 V for 12 minutes. Gels were fixed in 45% methanol and 10% acetic acid in water and imaged using Typhoon laser-scanning fluorescence imager (Cytiva).

### Patient-derived AML mouse xenografts

All mouse studies were conducted with approval from the Memorial Sloan Kettering Cancer Center Institutional Animal Care and Use Committee. NOD.Cg-Prkdc^scid^ Il2rg^tm1Wjl^/SzJ (NSG, Jackson Laboratory) mice were used for transplantation of primary human patient AML cells. NSG mice were irradiated with 200 cGy and transplanted with 1,000,000 cells per mouse via tail vein injection. For analysis, bone marrow cells were isolated by dissecting and crushing femoral and humeral bones using mortar and pestle in PBS supplemented with 2.5% FBS, and isolated by filtration through 70-μm mesh, followed by lysis of red cells using the RBC Lysis Buffer (BioLegend). In the case of subsequently performing fluorescence-activated cell sorting or analysis, bone marrow mononuclear cells were purified by density gradient centrifugation using Ficoll-Hypaque Plus, according to manufacturer’s instructions (GE Healthcare). For serial transplantation, purified cells were incubated overnight in StemSpan (STEMCELL Technologies) supplemented with 100 ng/*μ*l each of human SCF, FLT3 ligand, and TPO (PeproTech) at 37 °C with 5% CO_2_. Subsequently, cells were transplanted into 200 cGy-irradiated NSG mice with 1,000 cells or indicated cells numbers per mouse via tail vein injection. Transplanted mice were fed with Sulfatrim-supplemented chow.

### Mouse bone marrow transplantation

C57BL/6J mice were used as donors (The Jackson Laboratory). Upon dissection and crushing of femoral and humeral bones, bone marrow was filtered through 70-μm mesh, and red cells were lysed using the RBC Lysis Buffer (BioLegend). Hematopoietic stem and progenitor cells were isolated using magnetic purification with the mouse Lineage Cell Depletion Kit, according to the manufacturer’s instructions (Miltenyi Biotec). Recipient C57BL/6J mice were irradiated with 900 cGy and transplanted with isolated bone marrow hematopoietic stem and progenitor cells with or without CFSE-labeling using 500,000 cells per mouse via tail vein injection. Transplanted mice were fed with Sulfatrim-supplemented chow.

### Fluorescence-activated cell sorting and scanning

Isolated bone marrow cells were purified by density gradient centrifugation using Ficoll-Hypaque Plus, according to manufacturer’s instructions (GE Healthcare). Purified mononuclear cells were suspended in PBS with supplemented with 10 *μ*g/ml mouse gamma globulin (Jackson ImmunoResearch) and incubated on ice for 20 minutes. Subsequently, cells were resuspended in MACS buffer (PBS containing 2 mM ETDA and 2.5% FBS) supplemented with human FcR Blocking Reagent (Miltenyi Biotec) at 1:5 dilution and anti-human CD45-APC (BioLegend 304012) at 1:10 dilution, and incubated on ice for 20 minutes. After washing 2 times in MACS buffer, cells were resuspended in MACS buffer containing 1 *μ*M SYTOX Blue (Invitrogen) and processed by fluorescence-activated cell analyzer or sorter using LSRFortessa cell analyzer or FACSAria II cell sorter (BD Biosciences). Unless otherwise indicated, cells were gated by selecting SYTOX Blue-negative and human CD45-positive cells. For multi-color staining for cell surface markers, the following antibodies were used additionally: PerCP/Cy5.5 anti-human CD34 (BioLegend 343612), PE/Cy7 anti-human CD38 (BIoLegend 303516), APC/Cy7 human-CD45RA (BioLegend 304128), PE anti-human CD90 (BioLegend 328110), PE anti-human CD123 (BioLegend 306006), PE anti-human CD117 (BioLegend 313204), and APC/Cy7 anti-human CD244 (BioLegend 329518). Raw data were processed and quantified using FCS Express 7 (De Novo).

### Cell cycle profiling in vivo

5-ethynyl-2’-deoxyuridine (EdU) was obtained from baseclick (Munich, Germany), and dissolved in PBS at 5 mg/ml and stored at -20 °C. Twenty four hours before analysis, mice were treated with 50 mg/kg EdU in PBS via intraperitoneal injection. Upon isolation of bone marrow mononuclear cells as described above, cells were fluorescently labeled using EdU 647 Kit, according to the manufacturer’s instructions with the following modifications (baseclick). Cells were stained with Fixable Viability Dye eFluor 780 (eBiocience) in PBS on ice for 30 minutes. After washing cells with PBS supplemented with 1% BSA, cells were fixed using 4% paraformaldehyde in PBS, and incubated at room temperature for 15 minutes. After washing with PBS supplemented with 1% BSA, cells were permeabilized using Saponin-based permeabilization buffer at room temperature for 20 minutes. Fluorophore labeling was performed using Eterneon-Red 645 azide and catalyst solution for 30 minutes at room temperature, according to manufacturer’s instructions (baseclick). Labeled cells were subsequently stained with anti-human CD45-PE (BioLegend 304008) or anti-human CD45-APC (BioLegend 304012) and anti-cleaved caspase-3-PE (BD) at 4 °C for 20 minutes.

### Hoechst 33342 and Pyronin Y staining for cell cycle analysis

Cells were suspended in StemSpan media containing 10 μg/ml Hoechst 33342 (Invitrogen) and incubated for 45 minutes at 37 °C with 5% CO_2_. Pyronin Y (Sigma-Aldrich, P9172) was subsequently added to cell suspension at a final concentration of 3 μg/ml, and cells were incubated another 15 minutes at 37 °C with 5% CO_2_. Labeled cells were stained with anti-human CD45-PE/Cy7 (BioLegend 304016) in MACS buffer containing10 μg/ml Hoechst 33342 and 3 μg/ml Pyronin Y at 4 °C for 20 minutes as described above, and resuspended in 1.7 nM SYTOX Red (Invitrogen)-containing MACS buffer before processing by fluorescence-activated cell analyzer.

### Reactive Oxygen Species (ROS) level measurement

Cells were suspended in StemSpan media and seeded on 6-well plates. CellROX orange (Invitrogen) was added to each well at a final concentration of 0.5 μM, and cells were incubated for 30 minutes at 37 °C with 5% CO_2_. Collected cells were washed with MACS buffer twice and stained with anti-human CD45-APC at 4 °C for 20 minutes as described above. Cells were resuspended in MACS buffer containing 1 μM SYTOX blue before processing by fluorescence-activated cell analyzer.

### Chemotherapy treatment

Cytosine β-D-arabinofuranoside hydrochloride (AraC, Sigma) and doxorubicin hydrochloride (DXR, Sigma) were dissolved in PBS at 20 mg/ml and 0.6 mg/ml, respectively, and stored at - 20 °C. Mice were treated with 100 mg/kg AraC in PBS daily for 5 days and 3 mg/kg DXR in PBS daily for 3 days via intraperitoneal injection.

A-485 (Tocris) was dissolved in DMSO at 200 mg/ml and stored at -20 °C. Working solution for mouse studies was constituted of 10% A-485 200 mg/ml in DMSO, 45% PEG300, 5% Tween80 (Sigma-Aldrich) and 40% Saline, and used on the same day. Mice were treated with 100mg/kg/dose A-485 twice per day on the first day and once a day for subsequent 4 days via intraperitoneal injection.

### Genomic DNA sequencing analysis

DNA isolation, sequencing library preparation, and paired-end sequencing were performed using HiSeq and NovaSeq (Illumina) as described previously^57^. Hybridization capture and variant allele analysis were performed using the MSK-HemePACT panel of 585 genes recurrently mutated in hematological malignancies, as previously described^22^.

### Chromatin accessibility analysis

Purified cells (3,000 cells per sample) were lysed by incubation in 10 mM Tris pH 7.4, 10 mM NaCl, 3 mM MgCl_2_, 0.1% NP-40 for 2 minutes at 4 °C, followed by sedimentation at 1500 g to isolate nuclei. Tagmentation was performed using Nextera DNA sample prep kit (Illumina) at 37 °C for 30 minutes and subsequently stopped by addition of SDS to a final concentration of 0.2%. Tagmented DNA was purified using Agencourt AMPure XP beads (Beckman Coulter), and barcoded libraries were generated using the NEBNext Q5 Hot Start HiFi PCR Master Mix (New England Biolabs) and Nextera index primers (Illumina). Paired-end 50-bp sequencing (50 million reads per sample) was performed using HiSeq (Illumina). Sequencing reads were filtered for Q>15 and trimmed of adapter sequences using TrimGalore (v0.4.5), and aligned to hg19 using bowtie2 (v2.2.2). Peak calling was performed using MACS2 and filtered for blacklisted regions (http://mitra.stanford.edu/kundaje/akundaje/release/blacklists/). Signal was sequencing depth normalized and motif signatures were identified using Homer (‘findMotifsGenome.pl’).

### RNA sequencing

RNA from 5,000 cells per sample was extracted using the Quick-RNA MicroPrep kit, according to manufacturer’s instructions (Zymo Research). Barcoded libraries were constructed using QuantSeq 3’mRNA-Seq Library Prep Kit FWD for Illumina (Lexogen) with ERCC RNA spike-in Mix, according to the manufacturer’s instructions (Thermo Fisher). Single-end 50-bp sequencing was performed using HiSeq (Illumina) with 40 million reads per sample. Sequencing reads were filtered for Q>15 and adapter trimmed using TrimGalore (v0.4.5) before aligning to human assembly hg19 with STAR v2.5 using the default parameters. The raw counts matrix was built using HTSeq v0.6.1 and normalization and differential gene expression were performed using DESeq2 with default parameters.

### Single cell RNA-seq data analysis

Raw read counts were downloaded from the Gene Expression Omnibus (https://www.ncbi.nlm.nih.gov/geo/, accession GSE116256) together with metadata, cell classification and clustering scores, as published^37^. Total read counts per cell were normalized to 10,000 reads per cell. Normalized data were log-transformed using Seurat. Differential gene expression was performed using DESeq2 with the default parameters.

### Gene set enrichment analysis

Gene set enrichment analysis was performed using GSEAPreranked v4.4.0 with default parameters using DESeq2 output and MSigDB gene set version 6.0 (https://www.gsea-msigdb.org/gsea/msigdb/).

### Lentiviral cDNA library preparation

Doxycycline-inducible cDNA-expressing barcoded lentiviral plasmids (V191, Extended Data Table 8) were designed and synthesized by Transomic Technologies (Huntsville, Alabama). (One was synthesized by Custom DNA Constructs, Islandia, NY). Individual plasmids were sequence verified and quantified, and then pooled for lentivirus production. In total, 1.8 billion HEK293T cells grown in CellStack-5 (Corning) (30 million cells per CellStack-5) were transfected with pooled plasmids (600 μg per CellStack-5) and packaging plasmids, pMD2.G, and psPAX2 (300 μg and 300 μg per CellStack-5, respectively), using PEIpro according to the manufacturer’s instructions (Polyplus). Collected virus supernatant (7.5 liters) was 0.45-μm mesh filtered, and concentrated using Lenti-X concentrators, according to manufacturer’s instructions (Takara). Lentivirus titers were determined by infection of HEK293T cells as biological infectious units. Individual cDNA-expressing lentivirus vector was also prepared the same as above.

### Lentiviral transduction of human patient leukemia cells

Cells were infected with lentivirus preparations at multiplicity of infection of less than 0.2 by spinoculation at 800 g for 90 min at room temperature in StemSpan media supplemented with 100 ng/*μ*l each of human SCF, FLT3 ligand and TPO and 12.5 μl of LentiBOOST (SIRION Biotech) per well of 12-well plates, followed by incubation overnight at 37°C with 5% CO_2_. Upon replacement of media with fresh StemSpan supplemented with 100 ng/*μ*l each of human SCF, FLT3 ligand and TPO, cells were cultured for 3 days. Transduced cells were isolated using fluorescence-activated cell sorting by gating on SYTOX Blue-negative, human CD45- and mCherry-positive cells, and transplanted as described above.

### Inducible cDNA expression in human patient leukemia cells in vivo

Recipient NSG mice were fed with 625 mg/kg doxycycline hyclate chow (Envigo) or 625 mg/kg doxycycline hyclate chow and water containing 2 mg/ml doxycycline hyclate (Sigma) for one week before transplantation, followed by the same treatment upon transplantation. Transduced cells (1 million cells per mouse, unless otherwise indicated) were transplanted into NSG mice irradiated with 200 cGy via tail vein injection.

### Lentiviral cDNA library screening

Each experiment used 45 NSG mice divided into three groups of 15 mice as biological triplicates. Bone marrow cells were isolated and pooled from 15 mice per group and purified using fluorescence-activated cell sorting as described above. Genomic DNA was extracted from each sample using Quick-DNA Microprep Plus kit, according to manufacturer’s instructions (Zymo Research). Plasmid cDNA barcode sequences were amplified using primers containing indexed Illumina sequencing adaptors (Extended Data Table 10) using KOD Hot Start DNA Polymerase (Novagen). Amplicons were purified using Agencourt AMPure XP beads (Beckman Coulter). Single-end 50-bp sequencing was performed using NovaSeq (Illumina) with at least 10 million reads per sample. To quantify the read counts for each library sequence, a custom script was written based on the FASTX toolkit (http://hannonlab.cshl.edu/fastx_toolkit/). Each samples fastq file was processed as follows: The 3’ adapters were clipped and then reads were filtered to have a q value of 20 or more for all bases. The results sequences were then compared to the library sequence file to retain only sequences that match the library. Finally, the matching sequences were counted to give a count per library sequence. Total read counts were normalized to 10 million reads per sample for comparison.

### PCR amplification of cDNA-encoding regions

In competitive transplantation of JUN and TagBFP-transduced patient leukemia cells, genomic DNA was extracted from patient leukemia cells isolated from moribund mice using PureLink Genomic DNA Mini Kit (Invitrogen). Exogeneous JUN or TagBFP-encoding regions integrated to genome DNA were amplified with primers specific for TRE3GS promoter (5’-TTATGTAAACCAGGGCGCCT) and SV40 polyA (5’-AGCAGAGATCCAGTTTATCGACT) using Platinum SuperFi II PCR Master Mix (Thermo Fisher). Amplicons were resolved by gel electrophoresis using E-Gel EX Agarose Gels, 2% (Invitrogen).

### CRISPR genome editing

To engineer loss-of-function mutations of JUN in patient leukemia cells, Cas9 crRNA:tracrRNA Ribonucleoprotein (RNP) complex electroporation was performed using Alt-R CRISPR-Cas9 System (Integrated DNA Technologies) according to the manufacturer’s instructions. Briefly, equimolar crRNA and ATTO550-labeled tracrRNA were mixed in IDTE Duplex Buffer, heated at 95 °C for 5 minutes, and ramped down to room temperature for 10 minutes. Alt-R S.p. HiFi Cas9 Nuclease V3 (Integrated DNA Technologies) and crRNA:tracrRNA duplex were mixed and incubated at room temperature for 20 minutes. Cells were washed with PBS and resuspended in Resuspension Buffer T (Neon, Invitrogen). A mixture of cells, Cas9:crRNA:tracrRNA complex and Alt-R Electroporation enhancer (Integrated DNA Technologies) were prepared, and electroporation was performed at a condition of 1600V, 10 msec, 3 pulses using Neon Transfection System (Invitrogen). gRNA sequences used for JUN knockout or control AAVS1 were listed in Extended Data Table 11.

### Tracing of Indels by Decomposition (TIDE) analysis

Genomic DNA was extracted from electroporated cells using PureLink Genomic DNA Mini Kit. Each gRNA-targeting genomic region was amplified with specific primer pairs (Extended Data Table 11) using Platinum SuperFi II PCR Master Mix. PCR products were purified using PureLink PCR Purification Kit (Invitrogen) and Sanger sequenced (Eton Bioscience). Sequencing data is analyzed using TIDE^54^, and genome editing efficiencies were estimated.

### Single-cell RNA sequencing sample preparation

Human patient leukemia cells harvested from mouse bone marrow were separated into 3 or 4 fractions based on CFSE signal intensity using fluorescence-activated cell sorting. Single cell suspensions were loaded onto Chromium Next GEM Chip G (10X Genomics PN 1000120) and GEM generation, cDNA synthesis, cDNA amplification, and library preparation of an expected 10,000 cells proceeded using the Chromium Next GEM Single Cell 3’ Kit v3.1 (10X Genomics PN 1000268) according to the manufacturer’s protocol. cDNA amplification included 11 cycles and sequencing libraries were prepared with 12-14 cycles of PCR. Indexed libraries were pooled equimolar and sequenced on a NovaSeq 6000 in a PE28/88 paired end run using the NovaSeq 6000 S4 Reagent Kit (200 cycles) (Illumina).

### Single-cell RNA sequencing data processing and analysis

Raw count matrices were generated with Cell Ranger v7.1.0 with the option to include introns set to True. Cells with <2500 genes detected or >7.5% mitochondrial reads were removed. Cell cycle scores were computed using default Seurat method (v4.3)^55^. We considered cells to be in G1 if they had scores < 0 for S and G2-M scores. For the automated annotation of leukemia samples, single cell RNA-seq data was projected onto a reference atlas of human healthy bone marrow hematopoiesis as had been described in previous work^56,57^ using a workflow based on scmap^73^. We obtained uMAP coordinates, cell type labels, and myelocyte pseudotime, where applicable. For the unsupervised analysis of G1 cells from MSK162, we followed the default Seurat implementation for dimensionality reduction and unsupervised clustering. 50 principal components were used to construct the k-nearest neighbor graph and to obtain the uMAP embedding. Monocytic LSC score (mLSC) was computed using the function AddModuleScore() from Seurat. The list of genes was derived from MLL-rearranged leukemia samples^74^, as shown in previous work^58^. Intra-patient differential expression analysis was computed using MAST^75^ and Bonferroni correction was applied to account for multiple hypothesis testing. FDR=0.1 was used to consider genes differentially expressed. Single-cell regulon activities were computed with single-cell regulatory network inference and clustering (SCENIC)^59,60^ following the steps described in ref. 60. In brief, gene regulatory networks were inferred from the raw count matrix using the method grn2boost, then regulon prediction was carried out using a cisTarget database with 10kb regulatory regions around the TSS. Single-cell regulon activities were computed using the AUCell package.

### Western blot analysis

Cells were lysed in Laemmli sample buffer (BioRad) supplemented with cOmplete protease inhibitors (Roche) and 100 mM DTT (BioRad) at a ratio of 100 μL sample buffer per 1 million cells. Cell suspensions were incubated at 95 °C for 7 min, and lysates were clarified by centrifugation for 5 min at 1500 rpm. Fourteen μL of clarified lysates were resolved by sodium dodecyl sulfate-polyacrylamide gel electrophoresis (SDS-PAGE) using 10% polyacrylamide Bis-Tris gels (Invitrogen) and transferred onto Immobilon FL PVDF or P PVDF membranes (Millipore, Billerica, MA, USA) at 30V for 90 min at 4 °C. For fluorescent western blotting, membranes were blocked using Odyssey Blocking buffer (Li-Cor, Lincoln, Nebraska, USA). Blots were incubated with primary antibodies for JUN (1:1000, CST 60A8), ETS1 (1:1000, CST 808A), beta-actin (1:5000, CST 8H10D10) or GAPDH (1:1000, CST D4C6R). Blotted membranes were visualized using secondary antibodies conjugated to IRDye 800CW (goat anti-mouse IgG, 1:15,000) or IRDye 680RD (goat anti-rabbit IgG, 1:15,000) and the Odyssey CLx fluorescence scanner (Li-Cor), according to manufacturer’s instructions (Li-Cor, Lincoln, Nebraska, USA). After visualization, signal intensity of the bands of interest was quantified using the Image Studio Lite (Li-Cor). For chemiluminescent western blotting, blotted membranes were blocked using 5% non-fat milk (Rockland) in TBS, incubated in primary antibodies as indicated, and visualized using HRP-conjugated secondary antibodies (donkey anti-rabbit, 1:15,000, and sheep anti-mouse, 1:15,000, GE Healthcare) and ECL substrate (SuperSignal West Femto Maximum Sensitivity Substrate and Atto Ultimate Sensitivity Substrate, ThermoFisher) with Amersham ImageQuant 800 OD, according to manufacturer’s instructions (Cytiva, Marlborough, MA, USA).

### Cell lines

Human AML cell lines, OCIAML2 and OCIAML3 were obtained from DSMZ and cultured in RPMI-1640 medium supplemented with 10% fetal bovine serum 100 U/ml penicillin and 100 μg/ml streptomycin in a humidified atmosphere at 37 °C and 5% CO2. HEK293T cells were obtained from ATCC and cultured in Dulbecco’s Modified Eagle medium (DMEM) supplemented with 10% FBS and 100 U/ml penicillin and 100 μg/ml streptomycin. All cell lines were authenticated by STR genotyping (Integrated Genomics Operation, Center for Molecular Oncology, MSKCC). The absence of Mycoplasma species contamination was verified using MycoAlert Mycoplasma detection kit, according to manufacturer’s instruction (Lonza).

### Statistical methods

Statistical analyses were performed using OriginPro 2018 and 2022 (Microcal). Survival analyses were performed using Kaplan-Meier log-rank test. Statistical significance values were determined using two-tailed Welch’s t-tests for continuous variables, and two-tailed Fisher exact tests for discrete variables. Confidence intervals were calculated using OriginPro 2018. Significance of gene set comparisons was assessed using hypergeometric tests as implemented in GeneOverlap v4.1^76^. Network diagrams were constructed using Cytoscape v3.9.1.

## Data and material availability

All data are available openly via Zenodo (DOI 10.5281/zenodo.6496278), with raw RNA-seq and ATAC-seq data available via NCBI Gene Expression Omnibus (accession number GSE205994). All plasmids are available via Transomic Technologies.

## Code Availability

All computational code is available via Zenodo.

## Acknowledgements.

We thank Michael Kharas, Alejandro Gutierrez, Peter van Galen, Marc Mansour, and Helen Mueller for critical discussions. We thank the MSKCC Integrated Genomics and Bioinformatics Cores, Flow Cytometry Core Facility, Antitumor Assessment Core Facility, Gene Editing and Screening Core, and Hematologic Oncology Tissue Bank, which are funded by MSKCC Support Grant NIH P30 CA008748, and the Integrated Genomics Operation Core, which is funded by the NCI Cancer Center Support Grant (CCSG, P30 CA08748), Cycle for Survival, and the Marie-Josée and Henry R. Kravis Center for Molecular Oncology. AK is a Scholar of the Leukemia & Lymphoma Society and acknowledges support from NIH R01 CA204396, R21 CA235285, Starr Cancer Consortium, St. Baldrick’s Foundation, Burroughs Wellcome Fund, Pershing Square Sohn Cancer Research Alliance, Mathers Foundation, Damon Runyon-Richard Lumsden Foundation, Mr. William H. and Mrs. Alice Goodwin and the Commonwealth Foundation for Cancer Research, and the Susan and Peter Solomon Divisional Genomics Program and the Center for Experimental Therapeutics at MSKCC. LKS is supported by the German Research Foundation (DFG, SCHM 3906/1-1).

## Author Contributions

AK and ST conceptualized the study; AK and ST designed and optimized experimental methods; ST, VM, FB, MU, AS, SR, SC and LKS performed experiments; ST, RK, SBC and LV analyzed data; AK and ST wrote the original manuscript, which was edited by all authors.

## References

1. Lapidot, T. et al. A cell initiating human acute myeloid leukaemia after transplantation into SCID mice. Nature 367, 645–648 (1994).

2. Bonnet, D. & Dick, J. E. Human acute myeloid leukemia is organized as a hierarchy that originates from a primitive hematopoietic cell. Nat. Med. 3, 730–737 (1997).

3. Goardon, N. et al. Coexistence of LMPP-like and GMP-like Leukemia Stem Cells in Acute Myeloid Leukemia. Cancer Cell 19, 138–152 (2011).

4. Sarry, J.-E. et al. Human acute myelogenous leukemia stem cells are rare and heterogeneous when assayed in NOD/SCID/IL2Rγc-deficient mice. J. Clin. Invest. 121, 384–395 (2011).

5. Bewersdorf, J. P. & Abdel-Wahab, O. Translating recent advances in the pathogenesis of acute myeloid leukemia to the clinic. Genes Dev. 36, 259–277 (2022).

6. Shallis, R. M., Wang, R., Davidoff, A., Ma, X. & Zeidan, A. M. Epidemiology of acute myeloid leukemia: Recent progress and enduring challenges. Blood Rev. 36, 70–87 (2019).

7. Papaemmanuil, E. et al. Genomic Classification and Prognosis in Acute Myeloid Leukemia. N. Engl. J. Med. 374, 2209–2221 (2016).

8. Fennell, K. A. et al. Non-genetic determinants of malignant clonal fitness at single-cell resolution. Nature 601, 125–131 (2022).

9. Iacobucci, I. et al. Modeling and targeting of erythroleukemia by hematopoietic genome editing. Blood 137, 1628–1640 (2021).

10. Tothova, Z. et al. Multiplex CRISPR/Cas9-Based Genome Editing in Human Hematopoietic Stem Cells Models Clonal Hematopoiesis and Myeloid Neoplasia. Cell Stem Cell 21, 547–555.e8 (2017).

11. You, X. et al. Asxl1 loss cooperates with oncogenic Nras in mice to reprogram the immune microenvironment and drive leukemic transformation. Blood 139, 1066–1079 (2022).

12. Díaz de la Guardia, R., et al. Engraftment characterization of risk-stratified AML in NSGS mice. Blood Adv. 5, 4842–4854 (2021).

13. Woiterski, J. et al. Engraftment of low numbers of pediatric acute lymphoid and myeloid leukemias into NOD/SCID/IL2Rcγnull mice reflects individual leukemogenecity and highly correlates with clinical outcome. Int. J. Cancer 133, 1547–1556 (2013).

14. Griessinger, E. et al. Frequency and Dynamics of Leukemia-Initiating Cells during Short-term Ex Vivo Culture Informs Outcomes in Acute Myeloid Leukemia Patients. Cancer Res. 76, 2082–2086 (2016).

15. Foudi, A. et al. Analysis of histone 2B-GFP retention reveals slowly cycling hematopoietic stem cells. Nat. Biotechnol. 27, 84–90 (2009).

16. Qiu, J., Papatsenko, D., Niu, X., Schaniel, C. & Moore, K. Divisional history and hematopoietic stem cell function during homeostasis. Stem Cell Rep. 2, 473–490 (2014).

17. Takizawa, H., Regoes, R. R., Boddupalli, C. S., Bonhoeffer, S. & Manz, M. G. Dynamic variation in cycling of hematopoietic stem cells in steady state and inflammation. J. Exp. Med. 208, 273–284 (2011).

18. Wilson, A. et al. Hematopoietic stem cells reversibly switch from dormancy to self-renewal during homeostasis and repair. Cell 135, 1118–1129 (2008).

19. Ebinger, S., et al. Characterization of Rare, Dormant, and Therapy-Resistant Cells in Acute Lymphoblastic Leukemia. Cancer Cell (2016) doi:10.1016/j.ccell.2016.11.002.

20. Ebinger, S. et al. Plasticity in growth behavior of patients’ acute myeloid leukemia stem cells growing in mice. Haematologica 105, 2855–2860 (2020).

21. Eddaoudi, A., Canning, S. L. & Kato, I. Flow Cytometric Detection of G0 in Live Cells by Hoechst 33342 and Pyronin Y Staining. in Cellular Quiescence: Methods and Protocols (ed. Lacorazza, H. D.) 49–57 (Springer, New York, NY, 2018). doi:10.1007/978-1-4939-7371-2_3.

22. Adane, B. et al. The Hematopoietic Oxidase NOX2 Regulates Self-Renewal of Leukemic Stem Cells. Cell Rep. 27, 238–254.e6 (2019).

23. Stevens, B. M., O’Brien, C., Jordan, C. T. & Jones, C. L. Enriching for human acute myeloid leukemia stem cells using reactive oxygen species-based cell sorting. STAR Protoc. 2, 100248 (2021).

24. Wunderlich, M. et al. AML cells are differentially sensitive to chemotherapy treatment in a human xenograft model. Blood 121, e90–e97 (2013).

25. Grommes, C. et al. Ibrutinib Unmasks Critical Role of Bruton Tyrosine Kinase in Primary CNS Lymphoma. Cancer Discov. 7, 1018–1029 (2017).

26. Miles, L. A. et al. Single-cell mutation analysis of clonal evolution in myeloid malignancies. Nature 587, 477–482 (2020).

27. Sun, Q.-Y. et al. Ordering of mutations in acute myeloid leukemia with partial tandem duplication of MLL (MLL-PTD). Leukemia 31, 1–10 (2017).

28. Schmalbrock, L. K. et al. Clonal evolution of acute myeloid leukemia with FLT3-ITD mutation under treatment with midostaurin. Blood 137, 3093–3104 (2021).

29. Wilson, N. K. et al. Combinatorial Transcriptional Control In Blood Stem/Progenitor Cells: Genome-wide Analysis of Ten Major Transcriptional Regulators. Cell Stem Cell 7, 532–544 (2010).

30. Diffner, E. et al. Activity of a heptad of transcription factors is associated with stem cell programs and clinical outcome in acute myeloid leukemia. Blood 121, 2289–2300 (2013).

31. Passegué, E., Wagner, E. F. & Weissman, I. L. JunB Deficiency Leads to a Myeloproliferative Disorder Arising from Hematopoietic Stem Cells. Cell 119, 431–443 (2004).

32. Santaguida, M. et al. JunB Protects against Myeloid Malignancies by Limiting Hematopoietic Stem Cell Proliferation and Differentiation without Affecting Self-Renewal. Cancer Cell 15, 341–352 (2009).

33. Park, C. S. et al. A KLF4-DYRK2–mediated pathway regulating self-renewal in CML stem cells. Blood 134, 1960–1972 (2019).

34. Bungartz, G., Land, H., Scadden, D. T. & Emerson, S. G. NF-Y is necessary for hematopoietic stem cell proliferation and survival. Blood 119, 1380–1389 (2012).

35. Wang, E. et al. Surface antigen-guided CRISPR screens identify regulators of myeloid leukemia differentiation. Cell Stem Cell 28, 718–731.e6 (2021).

36. Takakura, N. et al. A Role for Hematopoietic Stem Cells in Promoting Angiogenesis. Cell 102, 199–209 (2000).

37. Zhou, B. O., Ding, L. & Morrison, S. J. Hematopoietic stem and progenitor cells regulate the regeneration of their niche by secreting Angiopoietin-1. eLife 4, e05521 (2015).

38. Gíslason, M. H. et al. BloodSpot 3.0: a database of gene and protein expression data in normal and malignant haematopoiesis. Nucleic Acids Res. 52, D1138–D1142 (2024).

39. Arceci, R. J., Berman, J. N. & Meshinchi, S. Chapter 17 - Acute Myeloid Leukemia. in Cancer Genomics (eds. Dellaire, G., Berman, Jason. N. & Arceci, R. J.) 283–300 (Academic Press, Boston, 2014). doi:10.1016/B978-0-12-396967-5.00017-7.

40. Cabezas-Wallscheid, N. et al. Vitamin A-Retinoic Acid Signaling Regulates Hematopoietic Stem Cell Dormancy. Cell 169, 807–823.e19 (2017).

41. Wilson, A. et al. c-Myc controls the balance between hematopoietic stem cell self-renewal and differentiation. Genes Dev. 18, 2747–2763 (2004).

42. García-Prat, L. et al. TFEB-mediated endolysosomal activity controls human hematopoietic stem cell fate. Cell Stem Cell 28, 1838–1850.e10 (2021).

43. Signer, R. A. J., Magee, J. A., Salic, A. & Morrison, S. J. Haematopoietic stem cells require a highly regulated protein synthesis rate. Nature 509, 49–54 (2014).

44. Galen, P. van, et al. Single-Cell RNA-Seq Reveals AML Hierarchies Relevant to Disease Progression and Immunity. Cell 176, 1265–1281.e24 (2019).

45. Duy, C. et al. Chemotherapy Induces Senescence-Like Resilient Cells Capable of Initiating AML Recurrence. Cancer Discov. 11, 1542–1561 (2021).

46. Boyd, A. L. et al. Identification of Chemotherapy-Induced Leukemic-Regenerating Cells Reveals a Transient Vulnerability of Human AML Recurrence. Cancer Cell 34, 483–498.e5 (2018).

47. Farge, T. et al. Chemotherapy-Resistant Human Acute Myeloid Leukemia Cells Are Not Enriched for Leukemic Stem Cells but Require Oxidative Metabolism. Cancer Discov. 7, 716–735 (2017).

48. Morris, V. et al. Hypoxic, glycolytic metabolism is a vulnerability of B-acute lymphoblastic leukemia-initiating cells. Cell Rep. 39, (2022).

49. Eppert, K. et al. Stem cell gene expression programs influence clinical outcome in human leukemia. Nat. Med. 17, 1086–1093 (2011).

50. Ng, S. W. K. et al. A 17-gene stemness score for rapid determination of risk in acute leukaemia. Nature 540, 433–437 (2016).

51. Gentles, A. J., Plevritis, S. K., Majeti, R. & Alizadeh, A. A. Association of a Leukemic Stem Cell Gene Expression Signature With Clinical Outcomes in Acute Myeloid Leukemia. JAMA 304, 2706–2715 (2010).

52. Gal, H. et al. Gene expression profiles of AML derived stem cells; similarity to hematopoietic stem cells. Leukemia 20, 2147–2154 (2006).

53. Szklarczyk, D. et al. The STRING database in 2021: customizable protein–protein networks, and functional characterization of user-uploaded gene/measurement sets. Nucleic Acids Res. 49, D605–D612 (2021).

54. Brinkman, E. K., Chen, T., Amendola, M. & van Steensel, B. Easy quantitative assessment of genome editing by sequence trace decomposition. Nucleic Acids Res. 42, e168 (2014).

55. Tirosh, I. et al. Dissecting the multicellular ecosystem of metastatic melanoma by single-cell RNA-seq. Science 352, 189–196 (2016).

56. Beneyto-Calabuig, S. et al. Clonally resolved single-cell multi-omics identifies routes of cellular differentiation in acute myeloid leukemia. Cell Stem Cell 30, 706–721.e8 (2023).

57. Triana, S. et al. Single-cell proteo-genomic reference maps of the hematopoietic system enable the purification and massive profiling of precisely defined cell states. Nat. Immunol. 22, 1577–1589 (2021).

58. Pei, S. et al. A Novel Type of Monocytic Leukemia Stem Cell Revealed by the Clinical Use of Venetoclax-Based Therapy. Cancer Discov. 13, 2032–2049 (2023).

59. Aibar, S. et al. SCENIC: single-cell regulatory network inference and clustering. Nat. Methods 14, 1083–1086 (2017).

60. Van de Sande, B. et al. A scalable SCENIC workflow for single-cell gene regulatory network analysis. Nat. Protoc. 15, 2247–2276 (2020).

61. Xie, X. P. et al. Quiescent human glioblastoma cancer stem cells drive tumor initiation, expansion, and recurrence following chemotherapy. Dev. Cell 57, 32–46.e8 (2022).

62. Signer, R. A. J. et al. The rate of protein synthesis in hematopoietic stem cells is limited partly by 4E-BPs. Genes Dev. 30, 1698–1703 (2016).

63. Goncalves, K. A. et al. Angiogenin Promotes Hematopoietic Regeneration by Dichotomously Regulating Quiescence of Stem and Progenitor Cells. Cell 166, 894–906 (2016).

64. Jose, L. H. S. et al. Modest Declines in Proteome Quality Impair Hematopoietic Stem Cell Self-Renewal. Cell Rep. 30, 69–80.e6 (2020).

65. Galen, P. van, et al. Integrated Stress Response Activity Marks Stem Cells in Normal Hematopoiesis and Leukemia. Cell Rep. 25, 1109–1117.e5 (2018).

66. Shlush, L. I. et al. Tracing the origins of relapse in acute myeloid leukaemia to stem cells. Nature 547, 104–108 (2017).

67. Blatter, S. & Rottenberg, S. Minimal residual disease in cancer therapy – Small things make all the difference. Drug Resist. Updat. 21–22, 1–10 (2015).

68. Ishikawa, F. et al. Chemotherapy-resistant human AML stem cells home to and engraft within the bone-marrow endosteal region. Nat. Biotechnol. 25, 1315–1321 (2007).

69. Jones, C. L., Inguva, A. & Jordan, C. T. Targeting Energy Metabolism in Cancer Stem Cells: Progress and Challenges in Leukemia and Solid Tumors. Cell Stem Cell 28, 378–393 (2021).

70. Ye, H. et al. Leukemic Stem Cells Evade Chemotherapy by Metabolic Adaptation to an Adipose Tissue Niche. Cell Stem Cell 19, 23–37 (2016).

71. Takao, S. et al. Convergent organization of aberrant MYB complex controls oncogenic gene expression in acute myeloid leukemia. eLife 10, e65905 (2021).

72. Smeenk, L. et al. Selective Requirement of MYB for Oncogenic Hyperactivation of a Translocated Enhancer in Leukemia. Cancer Discov. 11, 2868–2883 (2021).

73. Kiselev, V. Y., Yiu, A. & Hemberg, M. scmap: projection of single-cell RNA-seq data across data sets. Nat. Methods 15, 359–362 (2018).

74. Somervaille, T. C. P. et al. Hierarchical Maintenance of MLL Myeloid Leukemia Stem Cells Employs a Transcriptional Program Shared with Embryonic Rather Than Adult Stem Cells. Cell Stem Cell 4, 129–140 (2009).

75. Finak, G. et al. MAST: a flexible statistical framework for assessing transcriptional changes and characterizing heterogeneity in single-cell RNA sequencing data. Genome Biol. 16, 278 (2015).

76. Shen L, Sinai ISoMaM (2022). GeneOverlap: Test and visualize gene overlaps. R package version 1.32.0, http://shenlab-sinai.github.io/shenlab-sinai/.

