## Extended Data Figures and Tables for "Epigenetic mechanisms controlling human leukemia stem cells and therapy resistance"

Extended Data Figure 1.

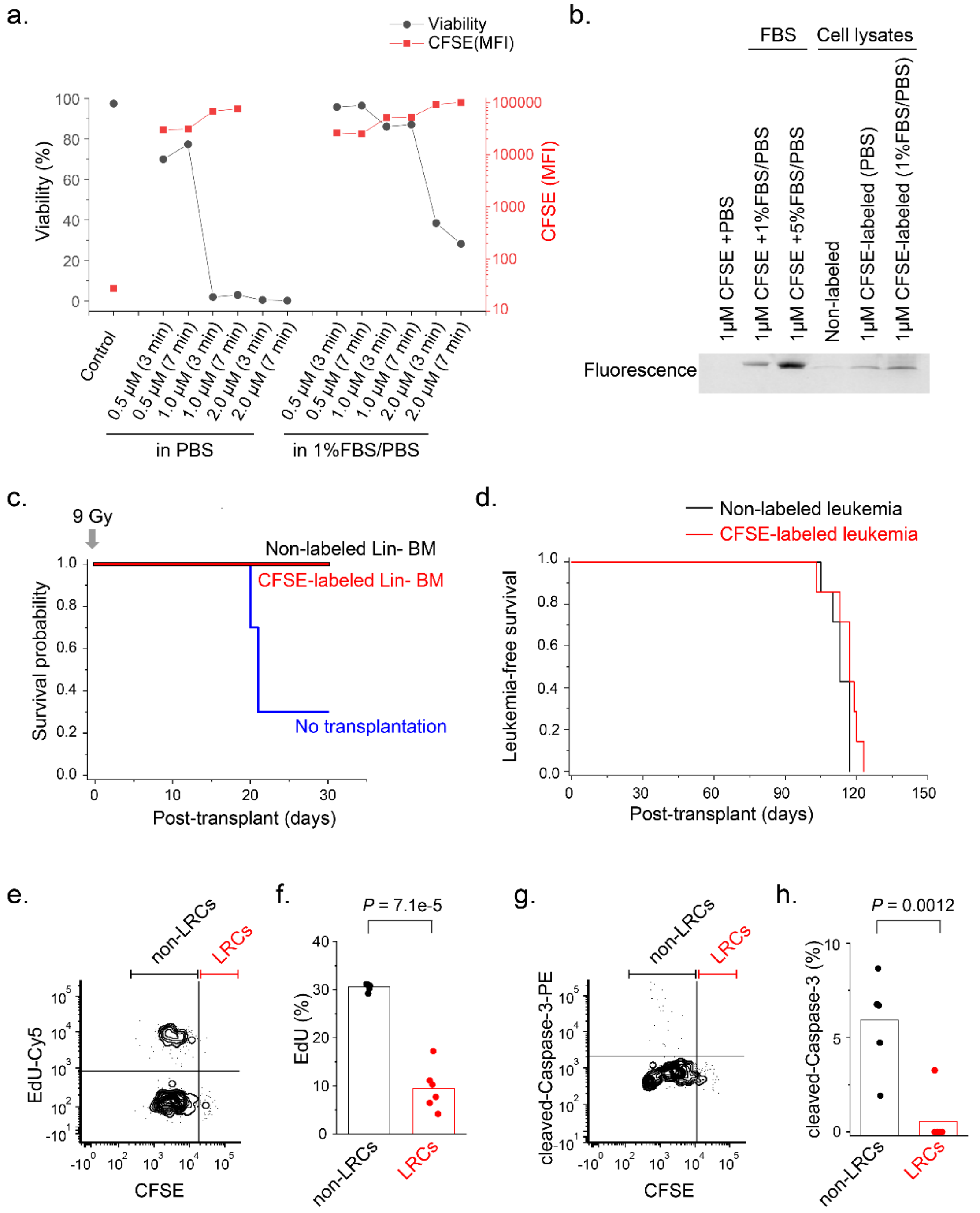


**Extended Data Figure 1. Optimized CFSE label tracing method to preserve normal hematopoietic and leukemia stem cell functions in vivo.**

**a.** To optimize CFSE labeling condition, OCI-AML2 cells were labeled with CFSE at the indicated concentrations (0 to 2 μM), at the indicated incubation time (3 or 7 minutes) and in two different buffers (PBS or PBS supplemented with 1% FBS). Red lines represent mean fluorescent intensity (MFI) of CFSE determined by fluorescence-activated cell scanning, and black lines represent cell viability determined by trypan blue exclusion one day after CFSE labeling.

**b.** Fluorescent gel imaging demonstrates fluorescent labeling of CFSE-labeled FBS or CFSE-labeled cell lysates in the indicated condition.

**c.** Overall survival of C57BL/6J mice transplanted with equal numbers of CFSE-labeled (red) or non-labeled (black) lineage cell-depleted (Lin-) bone marrow (BM) cells, or without BM transplantation (blue) after lethally irradiated with 900 cGy. CFSE-labeled Lin- BM cells completely rescued lethally irradiated mice, which is the same as non-labeled Lin- BM cells, while almost all non-transplanted mice died (log-rank $p = 4.6 \times 10^{-6}$).
**d.** Leukemia-free survival of NOD.Cg-Prkdc$^{scid}$ Il2r$^{gtm1Wjl}$/SzJ (NSG) mice transplanted with equal numbers of human MSK011 patient AML cells labeled CFSE (red) or non-labeled (black), where CFSE-labeled leukemia cells propagate leukemia in mice with similar kinetics as non-labeled leukemia cells. (log-rank $p = 0.10$).
**e-f.** Representative flow cytometry plots of bone marrow human leukemia cells harvested from mice transplanted with CFSE-labeled MSK011 patient AML cells 24 hours after EdU intraperitoneal administration (e). Label-retaining leukemia cells (LRCs) are less proliferative (red), as compared to non-label retaining leukemia cells (non-LRCs, black) which are actively proliferating (t-test $p = 7.1 \times 10^{-5}$). Bars represent mean values of measurement of 6 biological replicates (f).
**g-h.** Representative flow cytometry plots of cleaved caspase-3 stained bone marrow human leukemia cells harvested from mice transplanted with CFSE-labeled MSK011 patient AML cells (g). LRCs exhibit little to no detectable apoptosis (red), in contrast to non-LRCs (black), a fraction of which undergo apoptosis. (t-test $p = 0.0012$). Bars represent mean values of measurement of 6 biological replicates (h).

Extended Data Figure 2.

a.

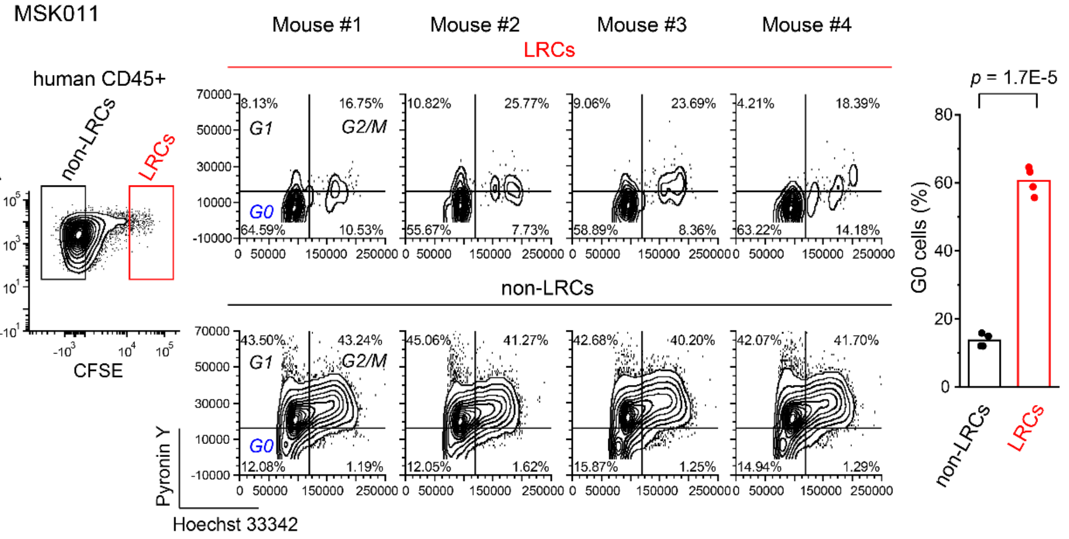


b.

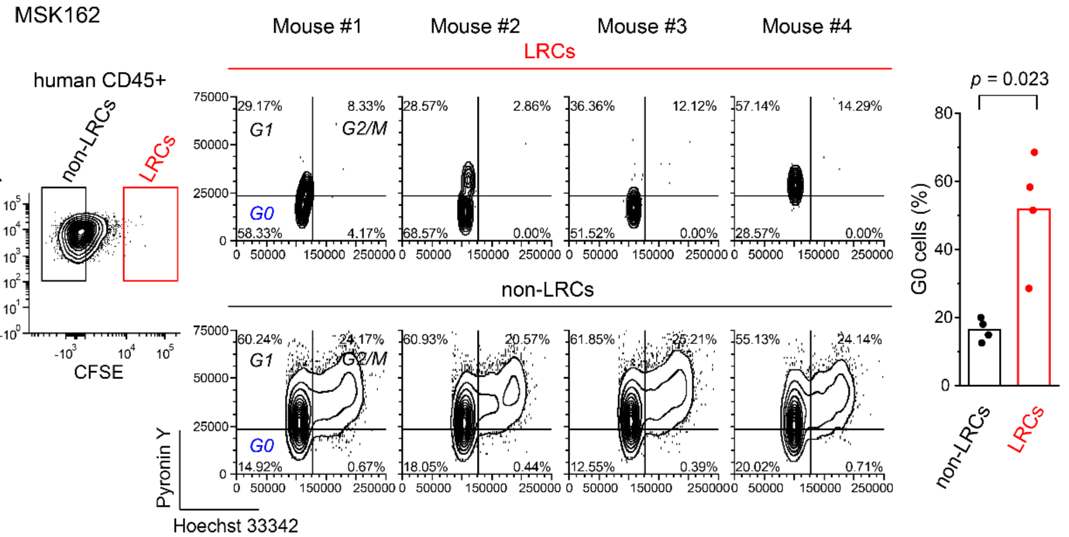


c.

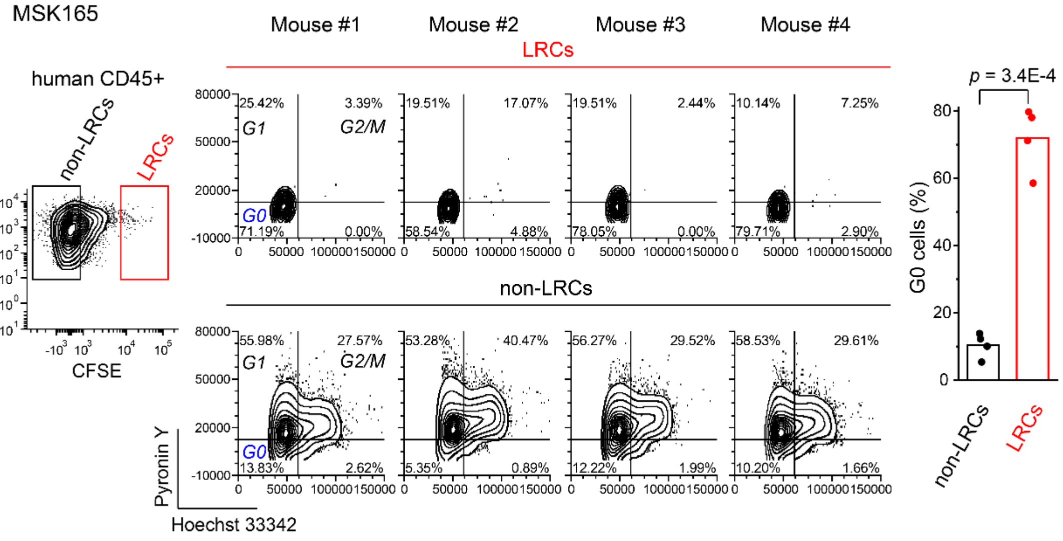

**Extended Data Figure 2. Most of patient AML LRCs exhibit G0 cell cycle phase.**
Hoechst 33342 and Pyronin Y (H-Y) staining was performed to analyze cell cycle status of bone marrow human leukemia cells isolated from mice transplanted with CFSE-labeled MSK011 (a), MSK162 (b) and MSK165 (c) patient AML cells.
**a-c.** Representative H-Y staining flow cytometry plots of LRCs (upper panels) and non-LRCs (lower panels) are shown. Most of LRCs (red) exhibit G0 cell cycle status, whereas most of non-LRCs (black) do not (t-test $p = 1.7 \times 10^{-5}$, $2.3 \times 10^{-2}$, and $3.4 \times 10^{-4}$ for MSK011, MSK162 and MSK165, respectively; right panels). Bars represent mean values of measurement of 4 biological replicates for each patient AML. (G0 phase: low Pyronin Y and low Hoechst 33342, G1 phase: high Pyronin Y and low Hoechst 33342, G2/M phase: high Pyronin Y and high Hoechst 33342).

### Extended Data Figure 3.

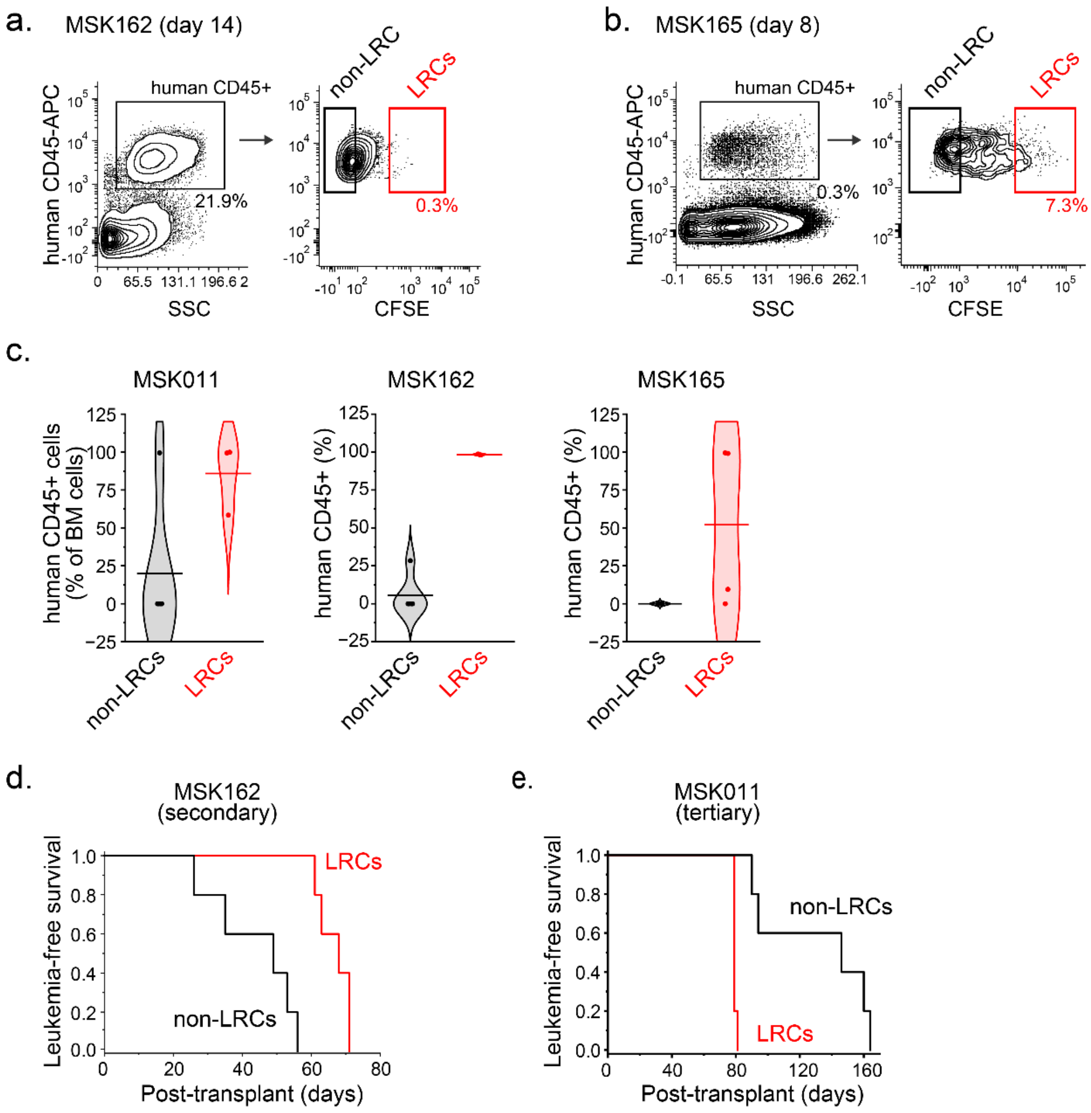


**Extended Data Figure 3. Patient AML LRCs maintain leukemia initiating property.**

**a-b.** Representative flow cytometry analysis of bone marrow human leukemia cells harvested from mice transplanted with CFSE-labeled MSK162 (a) and MSK165 (b) patient AML cells show gating strategies to isolate patient AML LRCs and non-LRCs by fluorescence-activated cell sorting.

**c.** Human CD45-positive cells are measured in bone marrow cells harvested from moribund mice after MSK011 (left panel), MSK162 (center panel), and MSK165 (right panel) patient AML LRCs (red) or non-LRCs (black) transplantation. Human leukemia cell engraftment is confirmed in most mice transplanted with LRCs.

**d.** Leukemia-free survival of mice secondary transplanted with equal numbers (800 cell/mouse) of MSK162 patient LRCs (red) or non-LRCs (black), where both LRCs and non-LRCs propagate leukemia (log-rank $p$ = 0.0018), but only LRCs initiate leukemias in tertiary recipients, whereas non-LRCs largely do not (refer to Figure 1c).

**e.** Leukemia-free survival of mice transplanted with equal numbers (1,000 cell/mouse) of MSK011 patient AML cells harvested from mice which are transplanted with MSK11 LRCs or non-LRCs and fully progressed leukemia (refer to Figure 1b). Both LRC progenies (red) or non-LRC progenies (black) propagate leukemia in tertiary recipient mice, although LRC progenies progressed faster than non-LRC progenies (log-rank $p$ = 0.00159).

Extended Data Figure 4.

a. MSK011

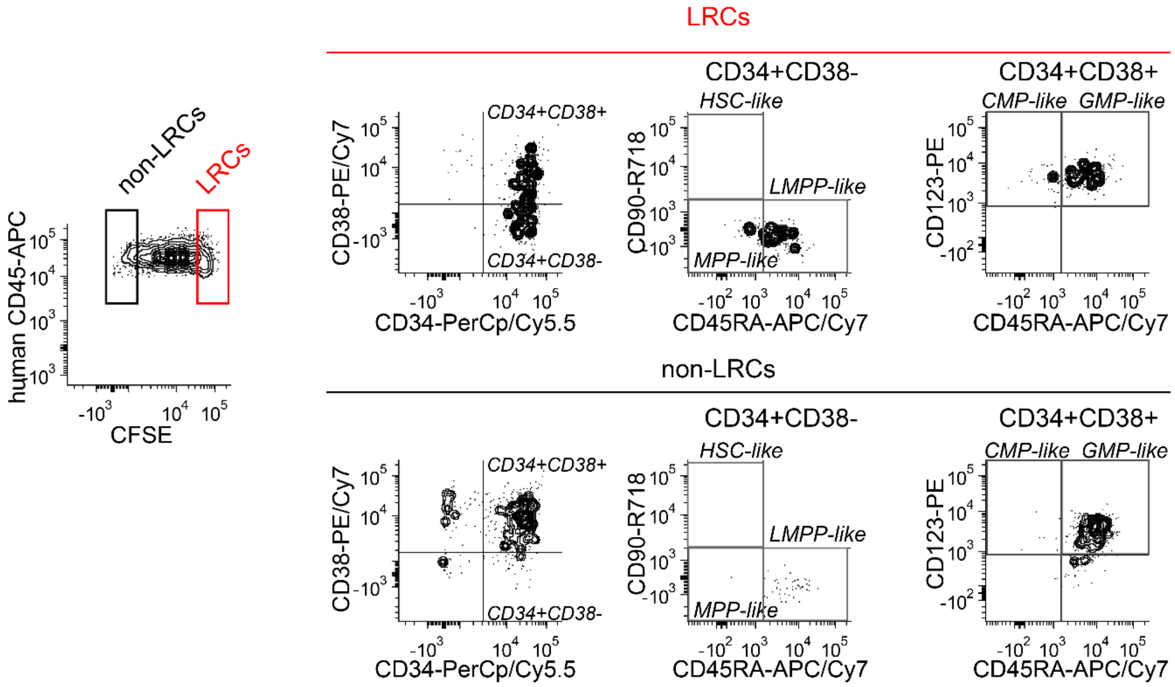


b. MSK162

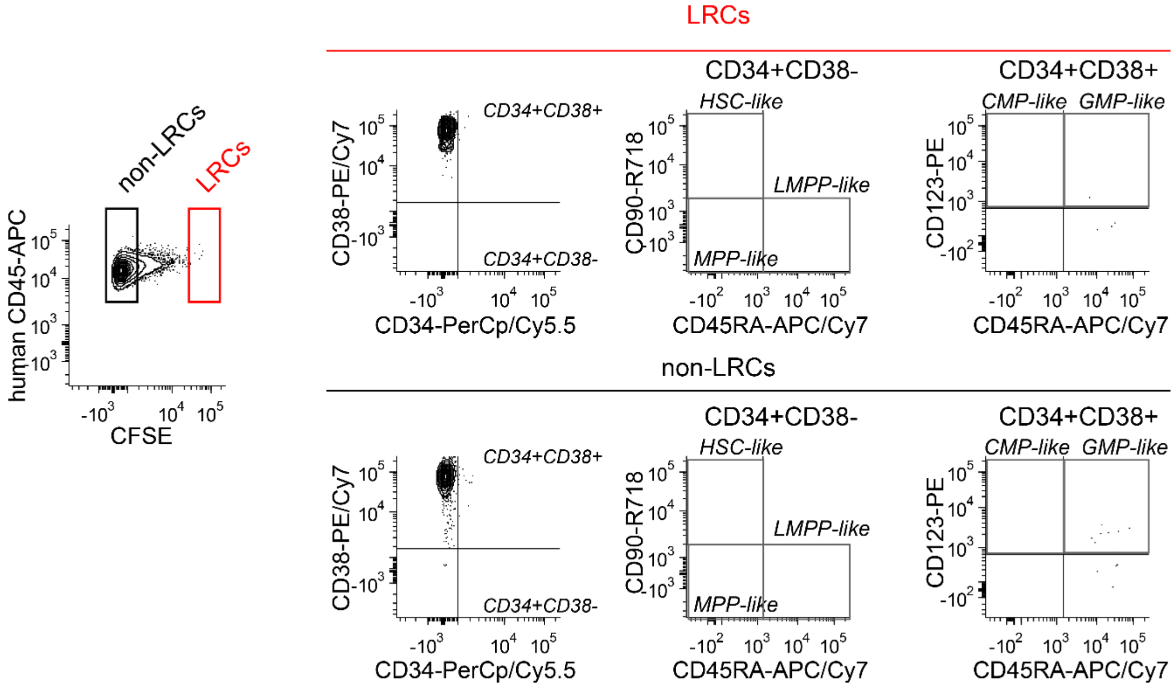

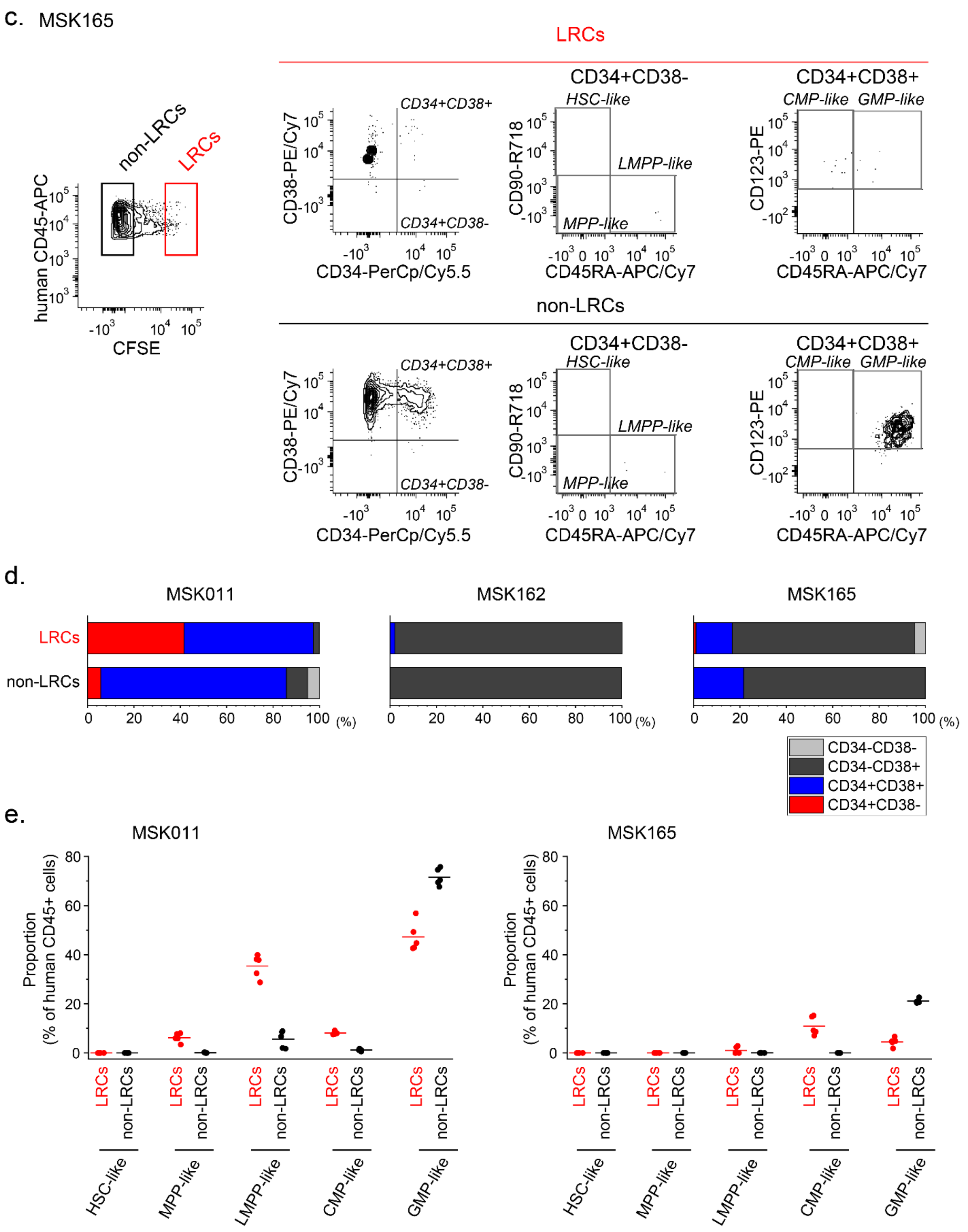


**Extended Data Figure 4. Quiescent patient AML LRCs exhibit variable association with currently known LSC surface markers.**

Multi-color flow cytometry analysis of bone marrow human leukemia cells isolated from mice transplanted with CFSE-labeled MSK011, MSK162, and MSK165 patient AML cells is performed to assess cell surface marker expression in LRCs and non-LRCs.

**a-c.** Representative multi-color flow cytometry plots represent gating strategies to classify cell types for MSK011 (a), MSK162 (b), and MSK165 (c). First, cells are gated by selecting SYTOX Blue-negative and human CD45-positive cells, and then LRCs and non-LRCs are separately analyzed (left panel). Cells are further gated into three cell populations based on CD34 and CD38 expression patterns: CD34+CD38-, CD34+CD38+, and CD34-CD38+ (second left panels). CD34+CD38- cells (second right panels) and CD34+CD38+ cells (right panels) are further classified based on CD45RA and CD90 expression or CD45RA and CD123 expression, respectively.

**d.** Stacked bar charts represent cell classification based on CD34 and CD38 surface expression in LRCs (upper bar) and non-LRCs (bottom bar). CD34+CD38- cells are partially enriched in MSK011 LRCs, as compared to non-LRCs. (41.6 and 5.7%, respectively; Fisher's exact test $p = 1.5 \times 10^{-9}$; left panel), but not in MSK162 (center panel) and MSK165 (right panel).

**c.** Dot plots represent further detailed cell classification in LRCs (red) and non-LRCs (black) of MSK011 (left panel) and MSK165 (right panel) patient AML cells based on the following criteria: Hematopoietic stem cell (HSC)-like; CD34+CD38-CD90+CD45RA-, Multipotential progenitor (MPP)-like; CD34+CD38-CD90-CD45RA-, Lymphoid-primed multipotential progenitor (LMPP)-like; CD34+CD38-CD90-CD45RA+, Common myeloid progenitor (CMP)-like; CD34+CD38+CD123+CD45RA-, and Granulocyte-monocyte progenitor (GMP)-like; CD34+CD38+CD123+CD45RA+.

In addition to LMPP-like cells, other progenitor-like cells, such as MPP-like and CMP-like are also partially and variably enriched in LRCs, as compared to non-LRCs.

#### Extended Data Figure 5.

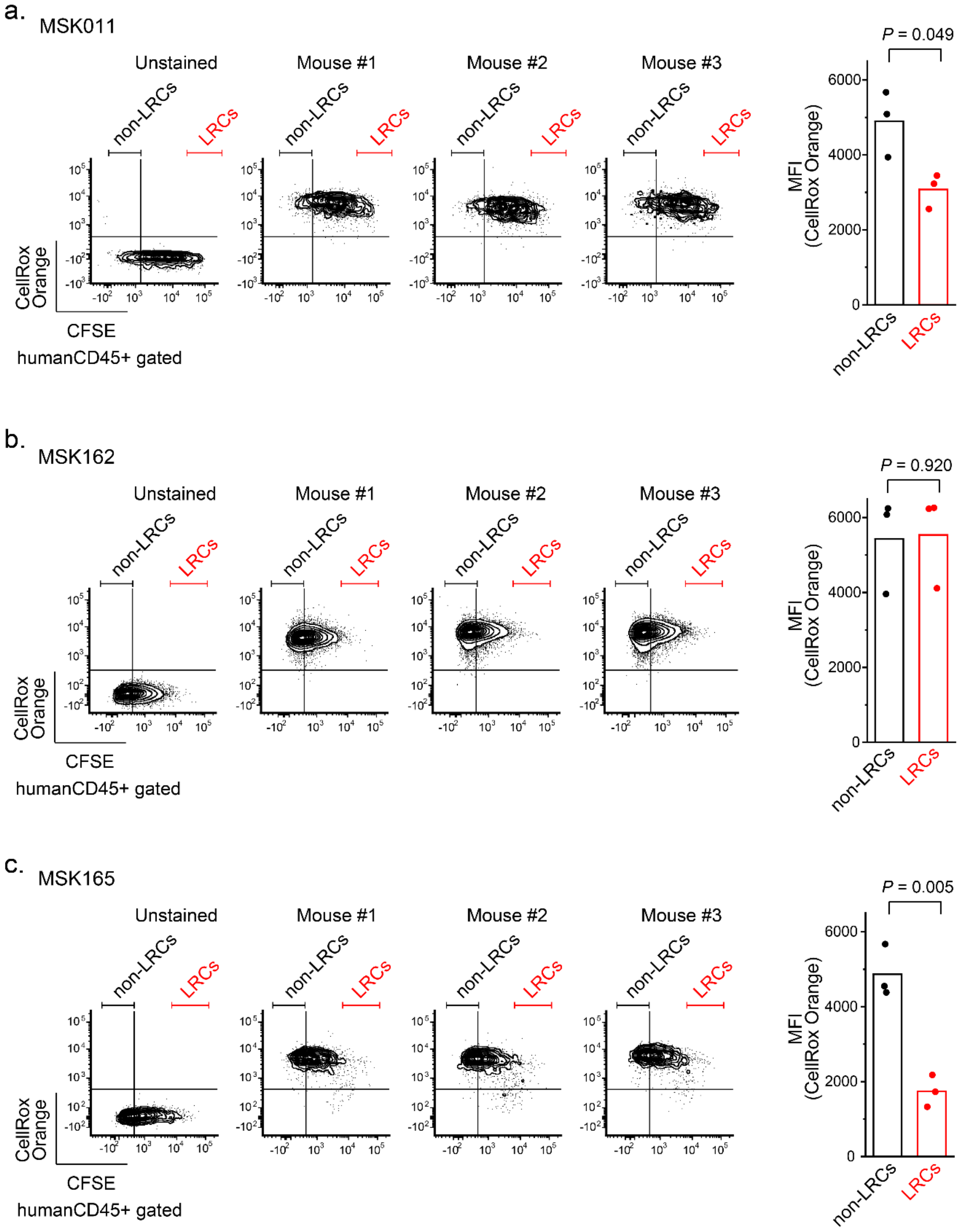


**Extended Data Figure 5. Variable differences in reactive oxygen species (ROS) levels between LRCs and non-LRCs.**

Reactive oxygen species (ROS) levels of bone marrow human leukemia cells isolated from mice transplanted with CFSE-labeled MSK011 (a), MSK162 (b) and MSK165 (c) patient AML cells are measured using the oxidation-sensitive fluorogenic probe CellROX by flow cytometry.

**a-c.** Representative flow cytometry plots of unstained (left panel) and CellROX-stained (3 center panels) human CD45-positive leukemia cells are shown. Mean fluorescence intensities (MFI) of CellROX in LRCs (red) and non-LRCs (black) are demonstrated in right panels. Differences in ROX levels between LRCs and non-LRCs are variable. MSK165 LRCs exhibit significantly lower CellROX fluorescence intensity as compared to its non-LRCs (MFI of 1743.5 vs 4864.0, respectively, t-test $p = 5 \times 10^{-3}$; c), whereas MSK162 LRCs and non-LRCs show similar CellROX fluorescence intensity (MFI of 5535.3 vs 5426.9, respectively, t-test $p = 0.92$; b). Bars represent mean values of measurement of biological triplicates.

Extended Data Figure 6.

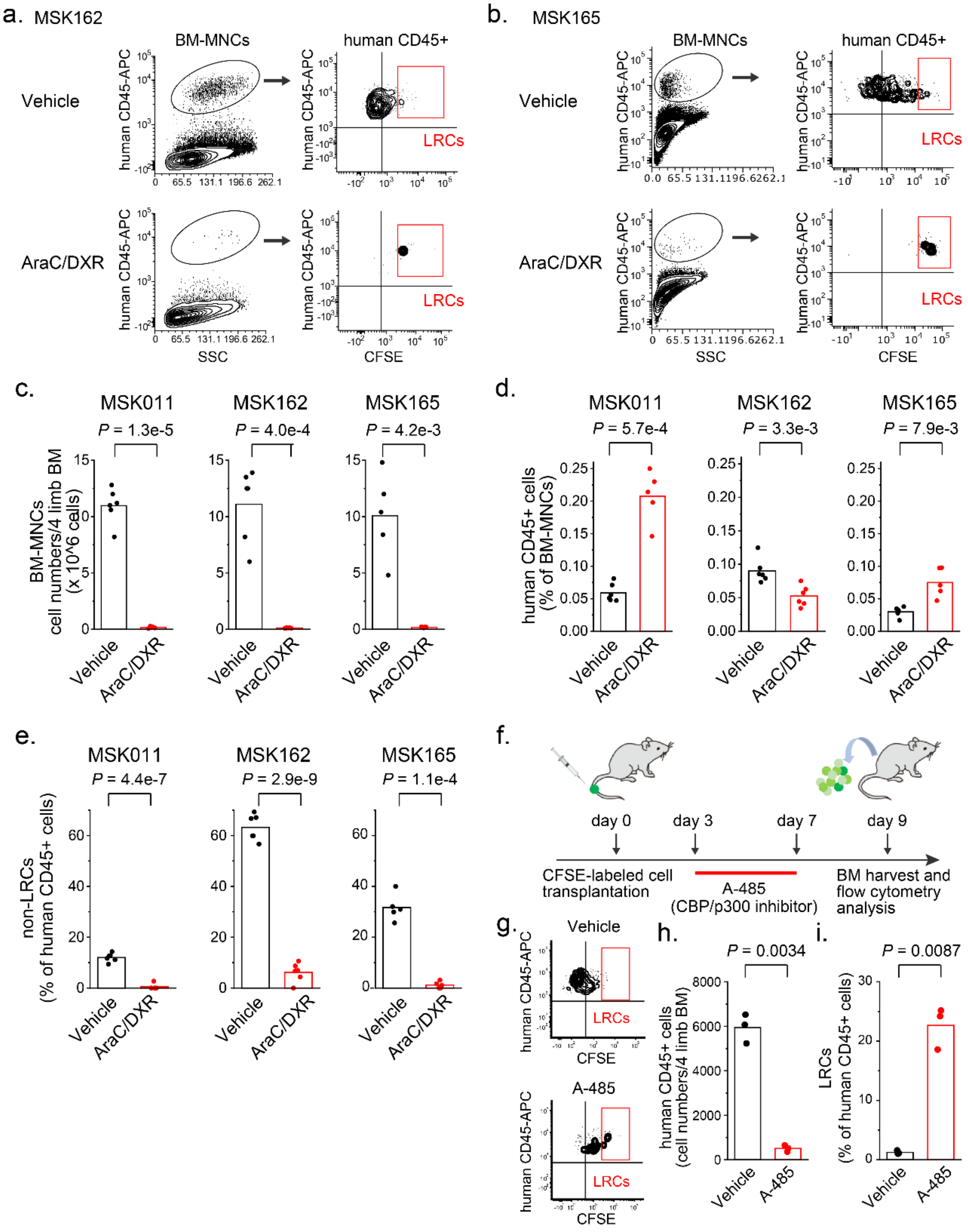


**Extended Data Figure 6. Quiescent patient AML cells exhibit chemotherapy resistance.**
**a-b.** Representative flow cytometry plots to analyze LRC frequencies in bone marrow human leukemia cells isolated from mice transplanted with CFSE-labeled MSK162 (a) and MSK165 (b) patient AML cells treated with AraC and DXR chemotherapy or vehicle.
**c.** Combined AraC and DXR chemotherapy treatment reduces bone marrow mononuclear cells (BM-MNCs) in MSK011 (left panel), MSK162 (center panel) and MSK165 (right panel) AML cells-transplanted mice (t-

test $p$ = 1.3 x $10^{-5}$, 4.0 x $10^{-4}$, and 4.2 x $10^{-3}$, respectively). Bars represent mean values of 6 biological replicates.
**d.** Differences in human CD45-positive patient leukemia cell percentage in BM-MNCs of AraC and DXR-versus vehicle-treated mice are various, while the absolute cell numbers of human CD45-positive patient leukemia cells are significantly reduced in AraC and DXR-treated mice compared to vehicle-treated mice (refer to Figure 1g). Bars represent mean values of 6 biological replicates.
**e.** Non-LRC frequencies of MSK011 (right panel), MSK162 (center panel) and MSK165 (right panel) patient AML cells are markedly reduced upon AraC and DXR chemotherapy (red), as compared to vehicle-treated controls (black), exhibiting that these patient AML non-LRCs are sensitive to AraC and DXR chemotherapy, as opposed to LRCs (t-test $p$ = 4.4 x $10^{-7}$, 2.9 x $10^{-9}$ and 1.1 x $10^{-4}$, respectively;refer to Figure 1h). Bars represent mean values of measurement of 6 biological replicates.
**f.** Experimental design for the analysis of mice transplanted with CFSE-labeled patient AML cells and treated with the CBP/p300 acetyltransferase inhibitor A-845.
**g.** Representative flow cytometry plots to analyze LRC frequencies in bone marrow human leukemia cells isolated from mice transplanted with CFSE-labeled MSK162 patient AML cells and treated with A-485 (lower panel) or vehicle (upper panel).
**h.** A-485 treatment reduces bone marrow disease burden of human CD45-positive MSK162 leukemia cell numbers in mice (t-test $p$ = 3.4 x $10^{-3}$). Bars represent mean values of measurement of biological triplicates.
**i.** LRC frequencies of MSK162 patient AML cells are significantly increased upon A-485 treatment (red) as compared to vehicle-treated controls (black), exhibiting that MSK162 patient AML LRCs are resistant to A-485 treatment (t-test p = 8.7 x $10^{-3}$), as well as AraC and DXR chemotherapy. Bars represent mean values of measurement of biological triplicates.

Extended Data Figure 7.

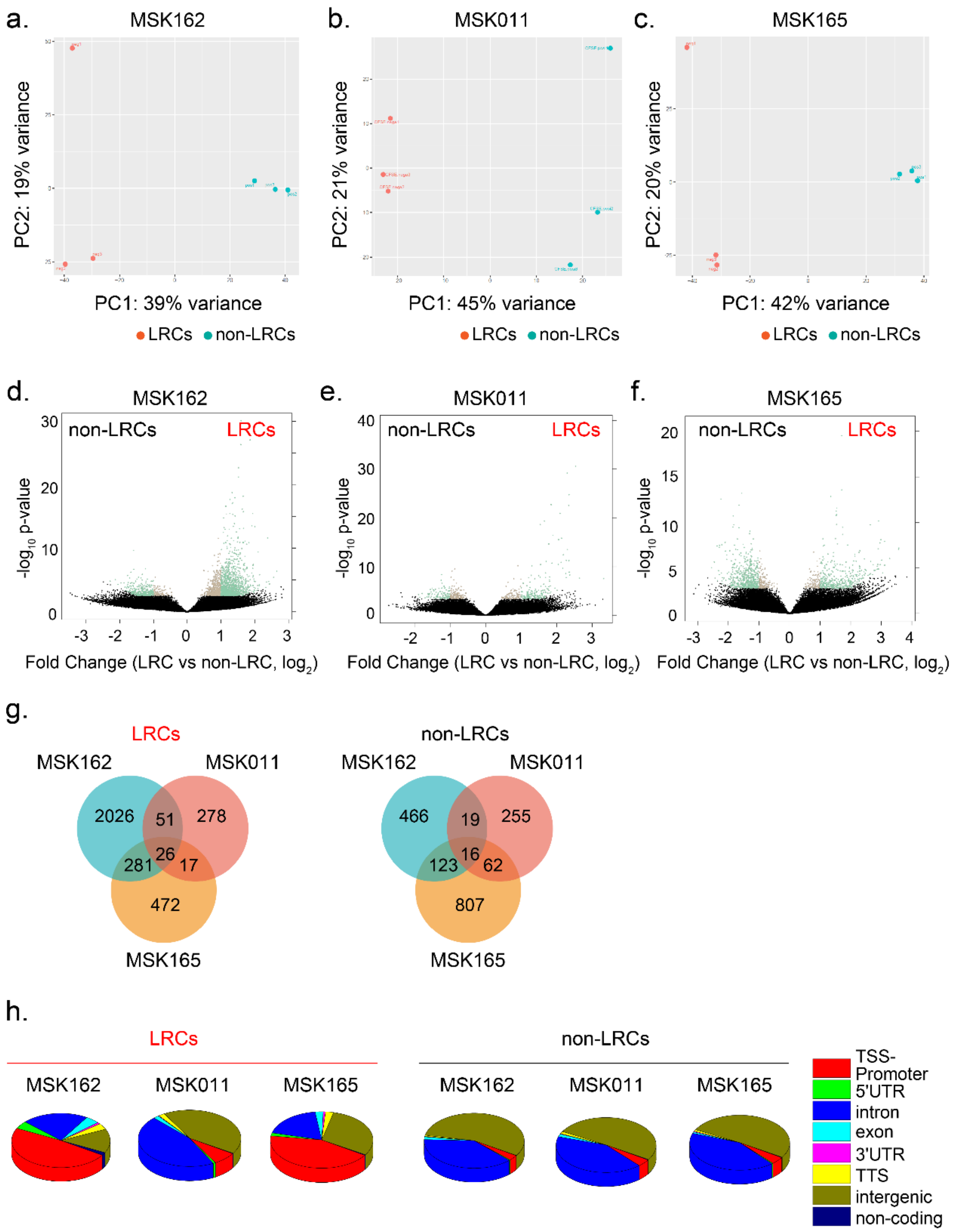


**Extended Data Figure 7. Quiescent patient AML LRCs exhibit promoter-centered distinct chromatin signatures (related to Figure 3).**

**a-c.** Principal component analysis (PCA) plots demonstrate segregation in differentially accessible chromatin loci between LRCs (red) versus non-LRCs (blue) determined by ATAC-seq in MSK162 (a), MSK011 (b) and MSK165 (c) human patient leukemias.

**d-f.** Volcano plots represent differentially accessible chromatin loci between LRCs versus non-LRCs with statistical significance of measurement triplicates in MSK162 (d), MSK011 (e) and MSK165 (f) human patient leukemias. (Green dots mark adjusted $p < 0.1$, fold change > 1.5)
**g.** Venn diagrams represent the numbers of significantly more accessible (right) or less accessible (left) chromatin loci in LRCs versus non-LRCs in MSK011, MSK162, and MSK165 human patient leukemias (adjusted $p < 0.1$, fold change > 1.5).
**h.** Pie charts represent the occupancy in genome components of increased accessible chromatin loci in LRCs (left) and non-LRCs (right) in MSK162 (left), MSK011 (center) and MSK165 (right), where quiescent LRCs exhibit increased chromatin accessibility of transcriptional start promoter regions as compared to non-LRCs.

Extended Data Figure 8.

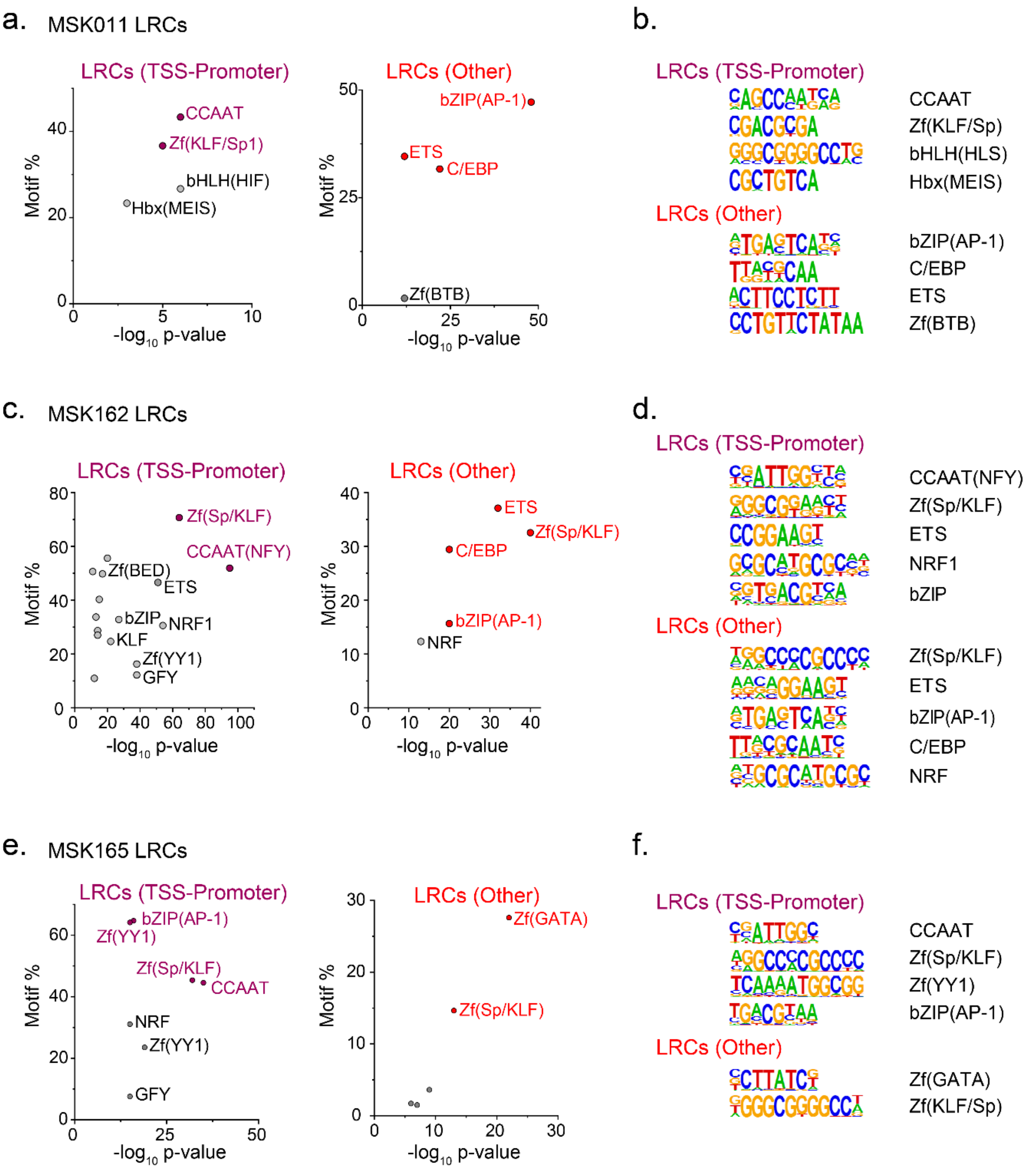


**Extended Data Figure 8. Specific transcription factor DNA binding sequence motifs are identified in quiescent patient AML LRCs (related to Figure 3).**
**a-f.** Transcription factor binding sequence motifs enriched in LRCs at transcription start site promoter (purple) and other non-promoter (red) regions in MSK011 (a), MSK162 (c) and MSK165 (e) as a function of their statistical significance of enrichment, with specific motif sequences shown (b, d and f, respectively).

Extended Data Figure 9.

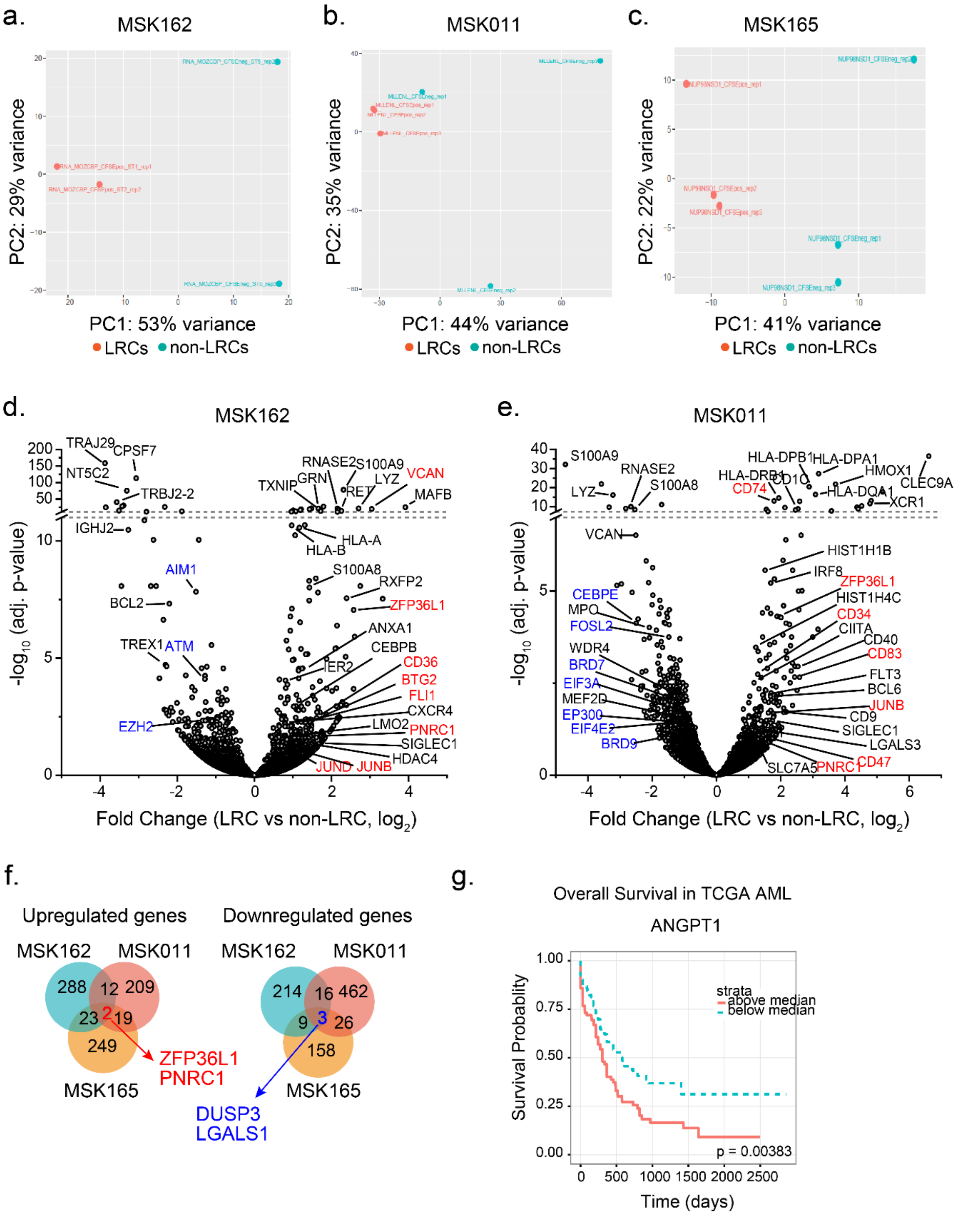


**Extended Data Figure 9. Commonly dysregulated gene signatures in quiescent patient AML LRCs. (related to Figure 4)**

**a-c.** Principal component analysis (PCA) plots demonstrate segregation in differentially regulated genes between LRCs and non-LRCs determined by RNA-seq in MSK162 (a), MSK011 (b) and MSK165 (c) human patient leukemias.

**d-e.** Volcano plots represent gene expression of MSK162 (d) and MSK011 (e) human patient AML cells with statistical significance of measurement triplicates as a function of differential gene expression of LRCs versus non-LRCs. Notable upregulated genes and downregulated genes are labeled in red and blue, respectively.

**f.** Venn diagrams showing the numbers of significantly upregulated (left) and downregulated (right) genes in LRCs versus non-LRCs in MSK011, MSK162, and MSK165 human patient leukemias (adjusted $p < 0.1$, fold change > 1.5).

**g**. Overall survival of the cohort of 172 AML patients in TCGA data exhibit that ANGPT1 expression is associated with inferior patient survival (log-rank $p = 0.0038$).

Extended Data Figure 10.

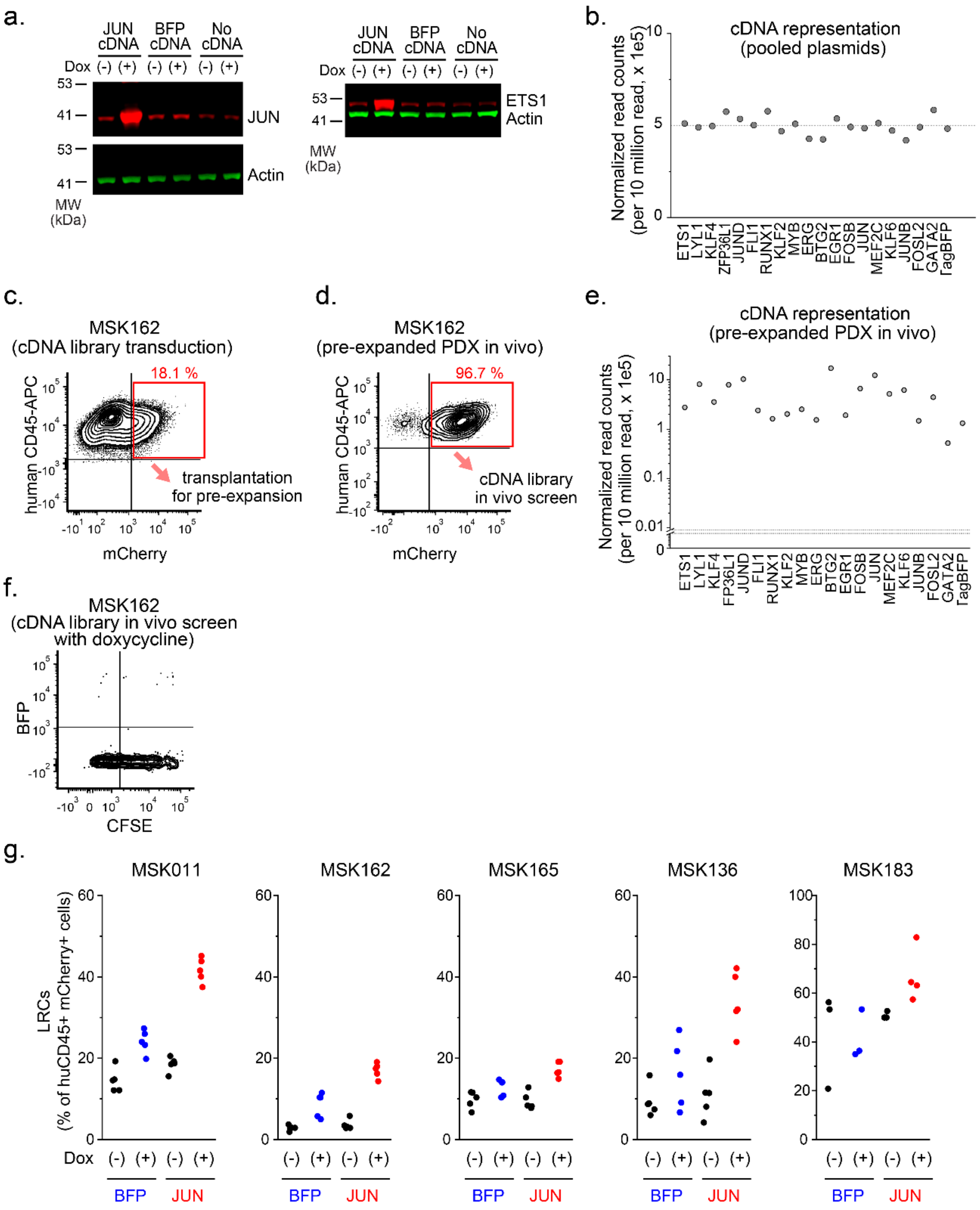


**Extended Figure 10. Functional cDNA library screen for LRC regulators in human patient AML in vivo identifies a distinct transcription network controlling AML LRC quiescence (related to Figure 6).**

**a.** Western blots demonstrating protein expression of target cDNA with or without doxycycline (Dox) treatment at 1μg/ml for 3 days in Dox-inducible cDNA-transduced OCI-AML3 cells, for JUN (left panel) and ETS1 (right panel). TagBFP cDNA-transduced and non-transduced OCI-AML3 cells serve as negative controls, and actin serves as a loading control.

**b.** Dot plots represent normalized read counts (10 million reads per sample) of each cDNA construct in pooled plasmid cDNA library measured by barcode sequencing, where equimolar representation of each cDNA is confirmed.
**c.** Representative flow cytometry plot of MSK162 patient leukemia cells 3 days after lentiviral transduction with LRC regulator cDNA library at the indicated multiplicity of infection (MOI), where appropriate transduction efficiency (<20 %) is confirmed by mCherry expression using flow cytometry.
**d.** Representative flow cytometry plots of bone marrow human leukemia cells harvested from mice transplanted with MSK162 patient leukemia cells genetically-modified with LRC regulator cDNA library, confirming stable mCherry expression after expanding in mice.
**e.** Dot plots represent normalized read counts (10 million reads per sample) of each cDNA construct in MSK162 patient leukemia cells genetically-modified with LRC regulator cDNA library after expanding in mice, where each cDNA representation is confirmed to be distributed within an appropriate range.
**f.** Representative flow cytometry plot of bone marrow human leukemia cells harvested from mice transplanted with CFSE-labeled MSK162 patient leukemia cells genetically-modified with LRC regulator cDNA library, where control TagBFP expression is confirmed by flow cytometry as a proof of successful Dox-inducible gene expression of cDNA library.
**g.** Patient AML cells are transduced with mCherry-expressing doxycycline-inducible *JUN* or TagBFP lentivirus vectors, and after labeled with CFSE, transplanted into NSG mice with or without doxycycline diet *in vivo* (refer to Figure 6e). LRC frequencies of human CD45-positive, mCherry-expressing cells are measured in *JUN* (red) or TagBFP (blue) transduced cells with or without doxycycline induction in five different patient AMLs: MSK011, MSK162, MSK165, MSK136 and MSK183.

Extended Data Figure 11.

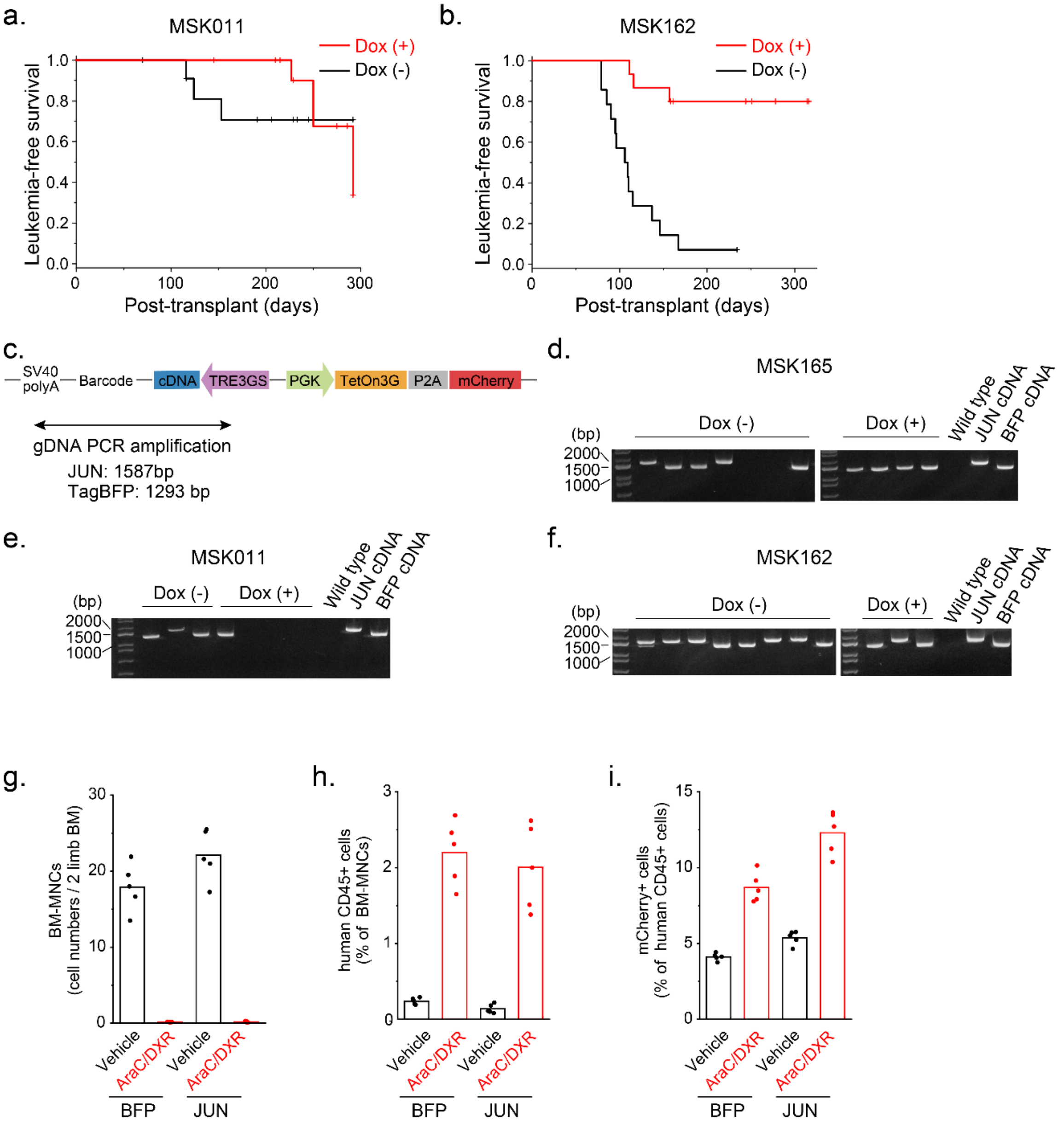


**Extended Data Figure 11. Enforced JUN expression impairs leukemia progression but confers chemotherapy resistance.**

**a-b.** Leukemia-free survival of mice transplanted with a mixture of *JUN* or TagBFP-transduced MSK011 (a) and MSK162 (b) patient leukemia cells with or without doxycycline diet is shown (12 and 15 mice per group for MSK011 and MSK162, respectively).

**c.** Exogenous *JUN* or TagBFP cDNA-encoding regions integrated into gDNA of engrafted human leukemia cells is amplified by PCR. Clonality is determined by the size of PCR products: 1587 bp for JUN and 1293 bp for TagBFP.

**d-f.** PCR products described above (c) are visualized on gels to determine the dominant clone in each mouse transplanted with a mixture of JUN or TagBFP-transduced MSK165 (d), MSK011 (e) and MSK162 (f) patient leukemia cells.

**g.** Mouse bone marrow mononuclear cells (BM-MNCs) are markedly reduced by combined AraC and DXR chemotherapy treatment regardless of JUN or TagBFP transduction (t-test $p = 2.3 \times 10^{-4}$ and $1.3 \times 10^{-4}$ for TagBFP and JUN transduction, respectively). Bars represent mean values of 5 biological replicates.
**h-i.** The proportions of human CD45-positive patient leukemia cells in mouse BM-MNCs (h) and mCherry-expressing cells in human CD45-positive cells (i) are analyzed in mice transplanted with MSK011 patient leukemia cells containing mCherry-expressing, JUN or TagBFP-transduced cells and treated with AraC/DXR or vehicle. Bars represent mean values of 5 biological replicates.

Extended Data Figure 12.

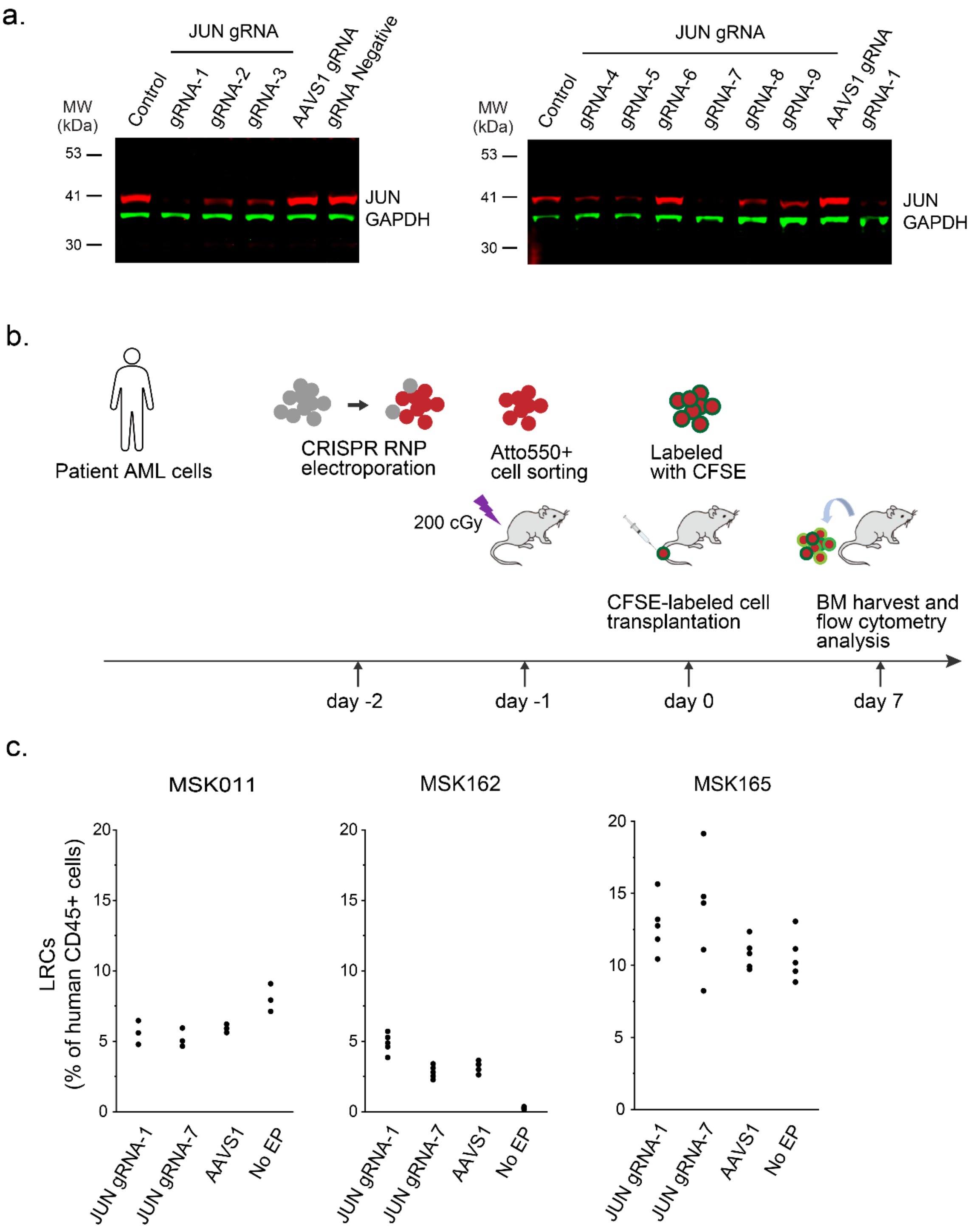


**Extended Data Figure 12. Loss of JUN does not change LRC frequencies in patient AML cells.**
**a.** Western blotting demonstrating protein expression of JUN in 9 different *JUN*-knockout OCIAML3 cells which are engineered using electroporation of Cas9 crRNA;tracrRNA ribonucleoprotein (RNP) complex. *AAVS1*-tageted and non-electroporated OCIAML3 cells serve as negative controls, and GAPDH serves as loading control. Two gRNAs, gRNA-1 and -7, are identified to exhibit loss of measurable JUN protein expression.

**b.** Experimental design to measure LRC frequencies of *JUN*-knockout patient leukemia cells, which are generated using Cas9 crRNA;ATT0550-labeled-tracrRNA RNP electroporation, isolated by fluorescence-activated cell sorting, labeled with CFSE, and transplanted into NSG mice.
**c.** LRC distribution of *JUN*-knockout MSK011 (left panel), MSK162 (center panel) and MSK165 (right panel) patient leukemia cells are measured by flow cytometry, which do not show substantial differences in LRC frequencies of *JUN*-knockout cells compared to control *AAVS1*-targted cells.

Extended Data Figure 13.

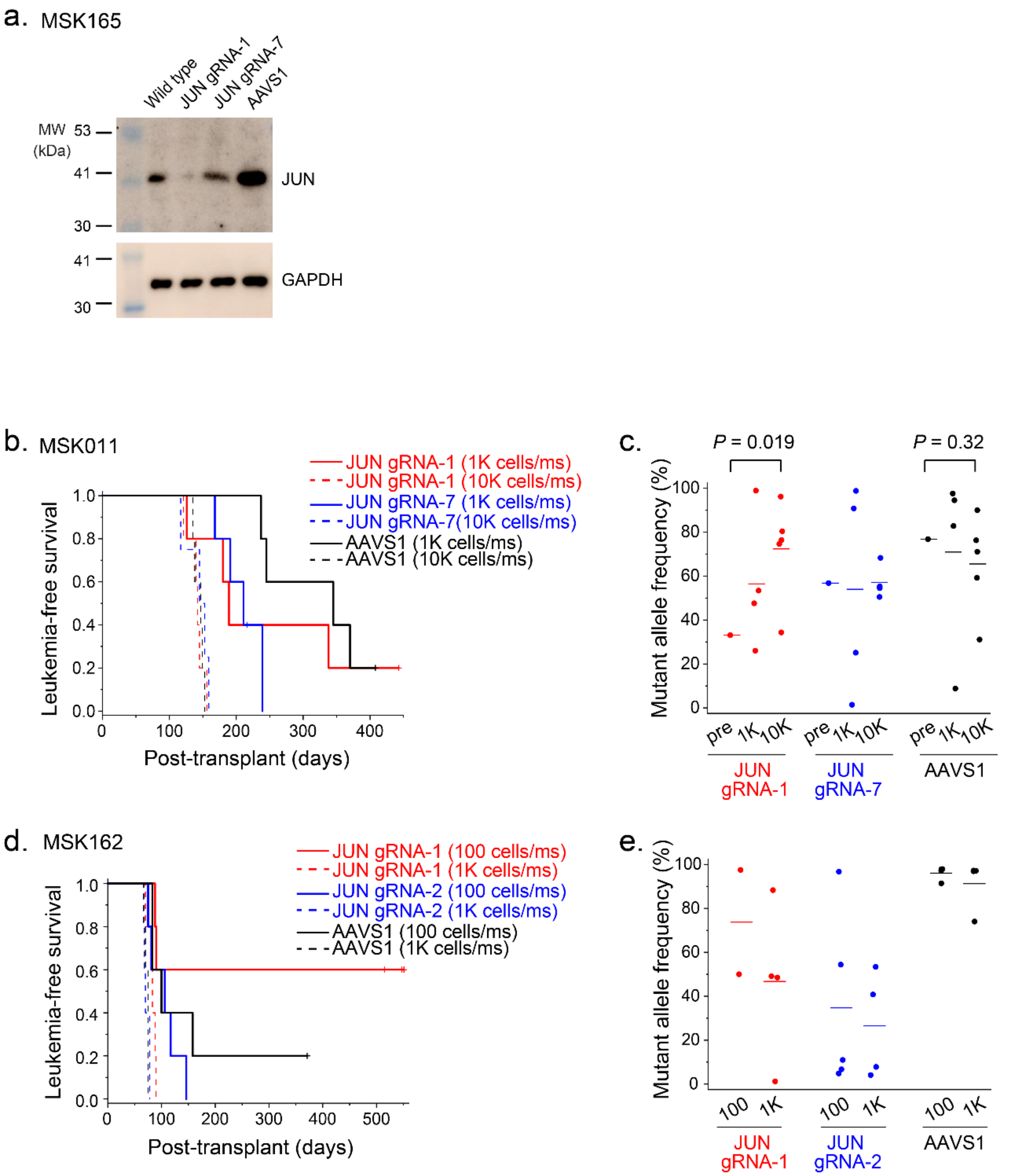


**Extended Data Figure 13. Loss of JUN promotes leukemia progression.**

**a.** Western blotting demonstrating reduced protein expression of JUN in *JUN*-knockout MSK165 patient leukemia cells, which are engineered using electroporation of Cas9 crRNA;tracrRNA ribonucleoprotein (RNP) complex. *AAVS1*-tageted and non-electroporated MSK165 cells serve as negative control, and GAPDH serves as loading control.

**b.** Leukemia-free survival of mice transplanted with *JUN*-knockout or control *AAVS1*-targeted MSK011 patient leukemia cells at different cell doses (100 cell/mouse and 1,000 cells/mouse, 5 mice for each group), where *JUN*-knockout cells propagate with similar kinetics as control *AAVS1*-targeted cells or ever faster in gRNA-7 *JUN*-knockout cells at 100 cells/mouse transplantation (log-rank $p = 0.023$).

**c.** Mutant allele frequencies are analyzed using the Tracing of Indels by Decomposition (TIDE) method before and after transplantation of *JUN*-knockout or control *AAVS1*-targeted MSK011 patient leukemia cells.

gRNA-1 *JUN*-knockout clones are enriched upon leukemia progression in vivo, whereas *AAVS1*-targeted cells are not (t-test p = 0.019, 0.91 and 0.32 for *JUN* gRNA-1, gRNA-7 and *AASV1*, respectively).

**d.** Leukemia-free survival of mice transplanted with *JUN*-knockout or control *AAVS1*-targeted MSK162 patient leukemia cells at different cell doses (100 cell/mouse and 1,000 cells/mouse, 5 mice for each group), where there is no significant difference in kinetics of leukemia progression between *JUN*-knockout or control *AAVS1*-targeted cells.

**e.** Mutant allele frequencies are analyzed using TIDE method after transplantation of *JUN*-knockout or control *AAVS1*-targeted MSK162 patient leukemia cells.

Extentended Data Figure 14.

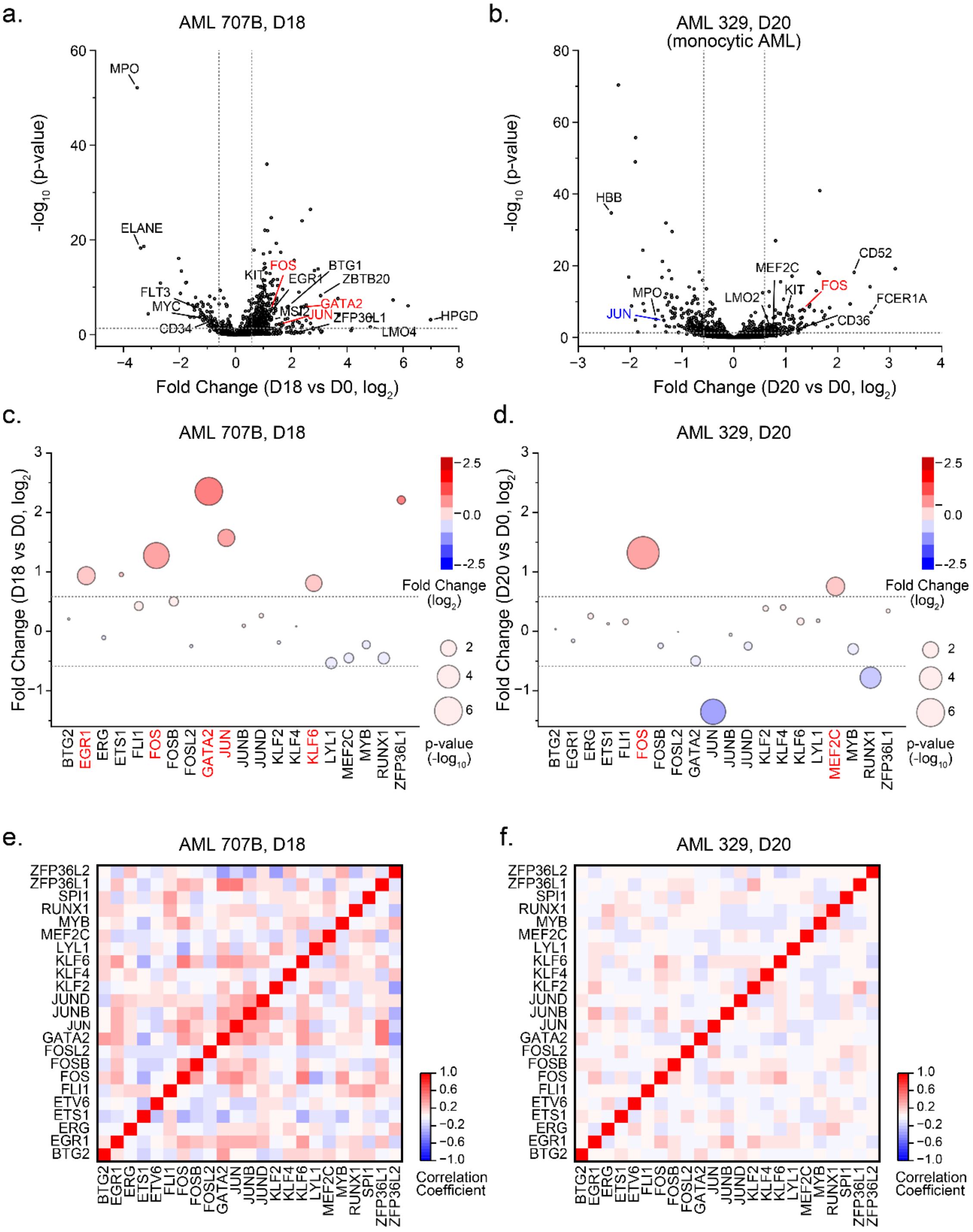


**Extended Data Figure 14. LRC regulatory factors are upregulated in residual leukemia cells after chemotherapy in AML patients.**
**a-b.** Volcano plots represent gene expression of human patient AML 707B (a) and AML 329 (b) (#42? Galen et al. Cell 2019) as a function of differential gene expression of patient bone marrow leukemia cells before and after induction chemotherapy at day 18 and day 20, respectively, with statistical significance of

measurement of individual single cells. Notable regulated genes including some LRC regulatory factors are labeled. (Dashed lines represent Fold change = ±1.5 and p-value = 0.05 for x- and y-axis, respectively).

**c-d.** Dot plots represent mRNA expression of LRC regulators in patient bone marrow residual leukemia cells of AML 707B (c) and AML 329 (d) as a function of differential gene expression before and after induction chemotherapy with statistical significance of measurement of individual single cells shown as circle size gradient of $-\log_{10}$ p-value. Blue to red color gradient represents relative decrease and increase of fold change of gene expression, respectively. (Dashed lines represent Fold change = ±1.5.) Some of the LRC regulatory factors, such as *JUN* and *GATA2* are upregulated in residual leukemia cells after chemotherapy in AML 707B, and another AP-1 factor *FOS* is upregulated in AML329.

**e-f.** Heatmaps show Pearson correlation coefficients of gene expression between each pair of LRC regulators in patient bone marrow residual leukemia cells after induction chemotherapy in AML707B (e) and AML 320 (f) with blue to red color gradient representing negative and positive values of correlation coefficients, respectively. In AML707B, LRC regulators, such as *JUN*, *GATA2*, and *ZFP36L1* are preferentially co-expressed together in individual leukemia cells, suggesting their cooperative regulation of chemotherapy resistance.

Extended Data Figure 15.

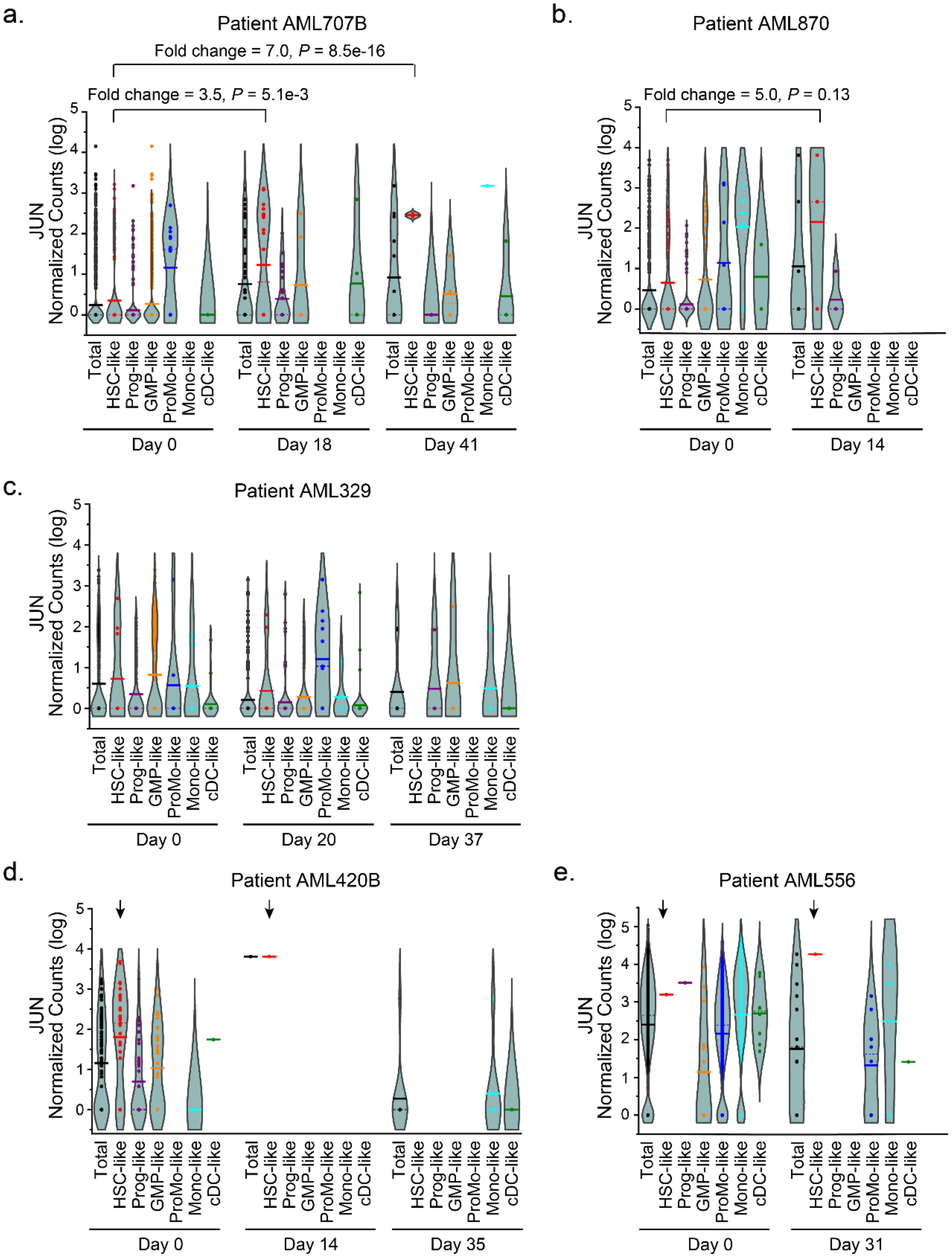


**Extended Data Figure 15. High *JUN* expression in HSC-like residual leukemia cells after chemotherapy in AML patients.**

**a-e.** *JUN* mRNA expression level in each cell type before and after chemotherapy treatment in diverse AML and monocytic AML patients, as measured using scRNA-seq[37]. Violin plots show normalized *JUN* read counts per cell including cells with zero reads in AML707B (a), AML870 (b), AML329 (c), AML420B (d) and

AML556 (e). JUN is upregulated in persistent leukemia cells, especially in HSC-like cells in AML707B and 870 (Fold Change = 3.5, $p$ = 0.0051 for AML 707B day 18; Fold Change = 7.0, $p = 8.5 \times 10^{-16}$ for AML 707B day 41; Fold change = 5.0, $p$ = 0.13 for AML 870). High JUN expression in HSC-like persistent leukemia cells is also maintained in patients with monocytic AML, AML 420B and AML 556) Bars and dashed lines represent mean and median values, respectively. (Black; total leukemia cells, Red: Hematopoietic stem cell (HSC)-like cells, Purple; Multipotent progenitor (progenitor)-like cells, Orange; Granulocyte-monocyte progenitor (GMP)-like cells, Blue; Promonocyte (ProMo)-like cells, Light blue; Monocyte (Mono)-like, Green; Conventional dendritic (cDC)-like cells).

Extended Data Figure 16.

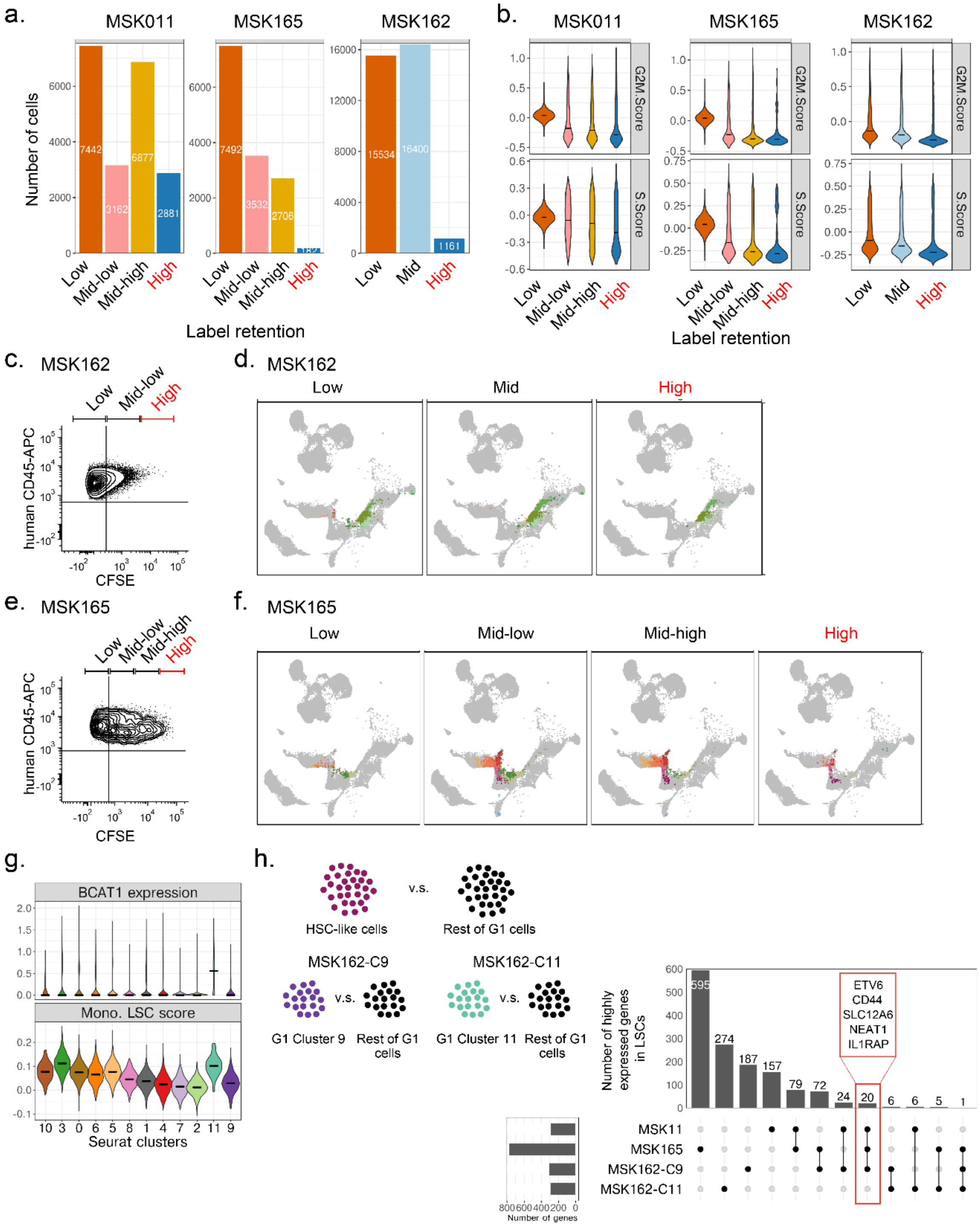


**Extended Data Figure 16. Single-cell analysis of LRC quiescence reveals shared and distinct gene expression programs. (Supplement data for Figure 8)**

**a.** Bars represent cell numbers which have passed stringent quality control filters in the indicated cell fractions based on the levels of label retention for MSK011 (left panel), MSK162 (center panel) and MSK165 (right panel).

**b.** G2M (upper panels) and S (lower panels) cell cycle scores computed using the default Seurat workflow, in which low label retaining cell fractions exhibit higher G2M and S cell cycle scores.
**c.** Representative flow cytometry plots to isolate human patient leukemia cells based on the levels of CFSE-label retention in MSK162.
**d.** uMAPs for high, middle and low-label retaining cells of MSK162 exhibit higher label retaining cell fractions are comprised of diverse cell types, including myelocyte-like cells.
**e.** Representative flow cytometry plots to isolate human patient leukemia cells based on the levels of CFSE-label retention in MSK165.
**f.** uMAPs for high, middle-high, middle-low and low-label retaining cells of MSK165 exhibit higher label retaining cell fractions are comprised of more diverse cell types, including HSC- and MPP-like cells.
**g.** *BCAT1* gene expression and monocytic LSC (mLSC) scores are measured in each of 12 clusters of MSK162 G1 cells, where cluster 11 shows high levels of *BCAT1* expression and high mLSC scores.
**h.** Gene expression features of cluster 9 of MSK162 G1 cells are partially overlapped with those of HSC-like cells in MSK011 and MSK165 high-LRCs.

Extended Data Figure 17.

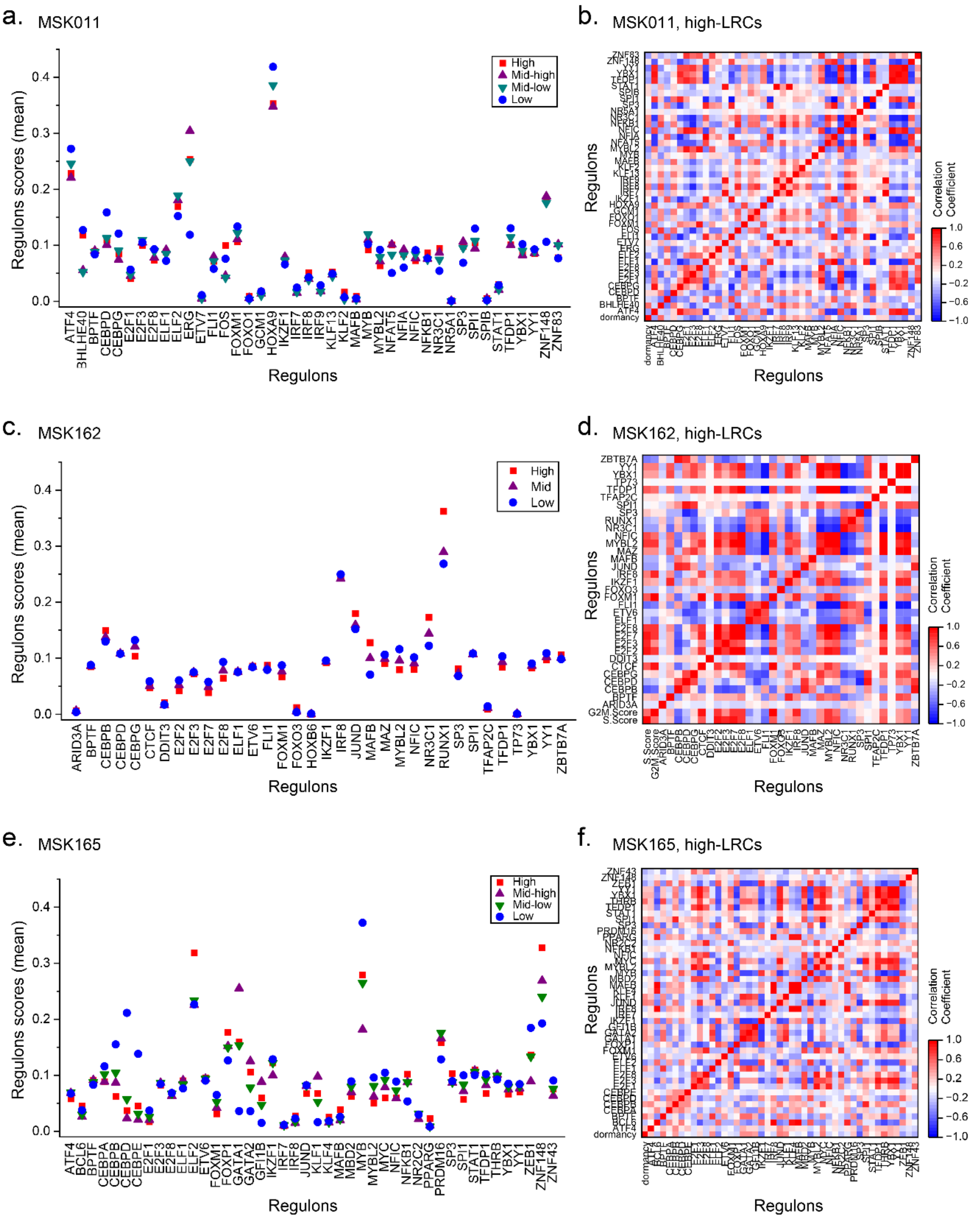


**Extended Data Figure 17. Shared and distinct transcription factor networks are associated with LRC quiescence.**
Each cell status is analyzed using single-cell regulatory network inference and clustering (SCENIC). Dot plots represent mean values of regulon scores of individual cells in the indicated cell fractions for MSK011 (a), MSK162 (c) and MSK165 (e). Specific regulons, including AP-1 and ETS TFs, such as FOS in MSK011, JUND in MSK162, and ELF2 in MSK165, are identified in LRCs..

Heatmaps show Pearson correlation coefficients of regulon scores between each pair of regulons in MSK011 (b), MSK162 (d) and MSK165 (f), with blue to red color gradient representing negative and positive values of correlation coefficients, respectively.

**Extended Data Table 1. Orthotopic transplantation of AML cells in immunodeficient mice isolated at diagnosis from patients with primary chemotherapy induction failure or remission**

**Primary transplantation**

| | Total patient specimens | Engrafted specimens | Engrafted (%) | Median Latency (days) | Mean Latency (days) |
|---|---|---|---|---|---|
| Induction Failure | 23 | 13 | 57 | 134 | 149 |
| Induction Remission | 25 | 9 | 36 | 96 | 139 |

**Secondary transplantation**

| | Total patient specimens | Engrafted specimens | Engrafted (%) | Median Latency (days) | Mean Latency (days) |
|---|---|---|---|---|---|
| Induction Failure | 5 | 4 | 80 | 107 | 126 |
| Induction Remission | 5 | 3 | 60 | 155 | 139 |

**Extended Data Table 2. Characteristics of genetically diverse patient AML specimens used in this study.**

| Patient ID | Age | Gender | Disease status | Genomic rearrangement | Somatic mutations |
|---|---|---|---|---|---|
| **MSK011** | 44 | M | Relapsed | MLL-ENL | RAD54L, SETD5, ARID1B, RECQL4, PTCH1, ETV6, KRAS, ATXN2, CREBBP, SETD6, TCF3, MLL4, POLD1, BCR, AR, NCOA3, WT1, FBXO11 |
| **MSK162** | 17 | M | Relapsed | MOZ-CBP | SPEN, ALK, IDH1, FAT1, TERT, IL7R, MLL, PIK3C2G, NCOR2, FLT3, IGF1R, GRIN2A, NCOR1, RPTOR, RPTOR, PTPRS, HLA-A, MLL2, MAPK1, MSH3, SETD1B |
| **MSK165** | 12 | F | Relapsed | NUP98-NSD1 | FLT3, IDH1, WT1 (S381*), WT1 (X216 splice), NOTCH2, BCORL1, PALB2, INPP4B, MED12, NOTCH3, SLX4 (A1221V), ASXL1, SLX4 (N457K), HLA-C, MCL1, TET2, STAG1, SETD2, CREBBP, LATS2, MLH1, HLA-C, PRKN, EPHA3, RPS6KB2, TCF7L2, PLCG2, SLX4 (M386V), SETD2, INHA, EP400, TAP1, SLX4 (L671S), SLX4 (R204C), ARID4B, RECQL4, SMG1, MGAM, SLX4, POLD1, MST1, SPRED1, ZFHX3, ANKRD11, EED, FLT1, BLM, FANCA, STAT5B, ERBB4 (L1008M), ERBB4 (M501V), NF1, MAP3K13 |
| **MSK136** | 19 | M | Refractory | inv(3) GATA2::MEOCM | NRAS, EP300, NCOR2, MLL, CEBPA, FANCF, MSH2, BARD1, WDR90 |
| **MSK183** | 50 | M | Relapsed | Normal karyotype | DNMT3A, NPM1, IDH1, FLT3, BCORL1 |

* Complete list of detected mutations is provided as Extended Data Table 3.

**Extended Data Table 3. Genetic variant allele frequencies (VAF) in LRCs and non-LRCs**

**a. MSK011**

| Gene | DNA change | Amino acid change | VAF in LRC | VAF in non-LRC |
|---|---|---|---|---|
| RAD54L | 604C>T | R202C | 0.523 | 0.502 |
| SETD5 | 3929C>T | S1310L | 0.485 | 0.491 |
| ARID1B | 4067C>A | P1356Q | 0.496 | 0.496 |
| RECQL4 | 3443C>T | S1148F | 0.501 | 0.506 |
| PTCH1 | 20C>G | A7G | 0.454 | 0.559 |
| ETV6 | 1252A>G | R418G | 0.463 | 0.497 |
| KRAS | 34G>T | G12C | 0.501 | 0.509 |
| ATXN2 | 212C>T | P71L | 0.421 | 0.517 |
| CREBBP | 1399G>A | A467T | 0.449 | 0.478 |
| SETD6 | 1373G>T | R458L | 0.468 | 0.513 |
| TCF3 | 1571C>T | P524L | 0.510 | 0.505 |
| MLL4 | 1043A>C | Q348P | 0.489 | 0.456 |
| POLD1 | 88C>T | R30W | 0.483 | 0.487 |
| BCR | 334G>A | E112K | 0.507 | 0.527 |
| AR | 475G>A | A159T | 0.997 | 0.997 |
| NCOA3 | 3741_3749delGCAGCAGCA | 1247_1250del | 0.291 | 0.287 |
| WT1 | 1107_1108insG | R370fs | 0.413 | 0.443 |
| FBXO11 | 169_170insAGC | P57_P58insQ | 0.262 | 0.264 |

**b. MSK162**

| Gene | DNA change | Amino acid change | VAF in LRC | VAF in non-LRC |
|---|---|---|---|---|
| SPEN | 2429G>A | R810Q | 0.490 | 0.513 |
| ALK | 3401A>T | Q1134L | 0.475 | 0.482 |
| IDH1 | 532G>A | V178I | 0.486 | 0.484 |
| FAT1 | 10478C>T | P3493L | 0.496 | 0.514 |
| TERT | chr5:1295228G>A | upstream | 0.539 | 0.433 |
| IL7R | 258G>T | R86S | 0.460 | 0.446 |
| MLL | 3180G>T | E1060D | 0.508 | 0.542 |
| PIK3C2G | 2307C>A | N769K | 0.455 | 0.464 |
| NCOR2 | 4690G>A | E1564K | 0.572 | 0.524 |
| IGF1R | 1976G>A | R659Q | 0.479 | 0.496 |
| GRIN2A | 2899G>C | V967L | 0.527 | 0.536 |
| NCOR1 | 6544G>A | A2182T | 0.468 | 0.425 |
| RPTOR | 2527G>A | V843M | 0.474 | 0.505 |
| RPTOR | 3842A>G | N1281S | 0.541 | 0.462 |
| PTPRS | 4151G>A | R1384Q | 0.512 | 0.526 |
| MAPK1 | 22_23insCGG | G8_A9insA | 0.203 | 0.161 |
| HLA-A | 618delG | T206fs | 0.156 | 0.156 |
| MLL2 | 5852dupC | P1951fs | 0.461 | 0.451 |
| FLT3 | 2503G>T | D835Y | not detected | 0.759 |
| MSH3 | 178_179ins CCGCAGCGGCCGCAGCGC | A60_A61insAAAAAP | not detected | 0.170 |
| SETD1B | 15dupC | H5fs | not detected | 0.681 |

**c. MSK165**

| Gene | DNA change | Amino acid change | VAF in LRC | VAF in non-LRC |
|---|---|---|---|---|
| FLT3 | chr13:28608244_28608245 insTCCCATTTGAGTCATA TTCACCCCCT | E604_F605ins GGEYDLKWE | 0.305 | 0.656 |
| IDH1 | chr2:209113113G>C | R132G | 0.466 | 0.529 |
| WT1 | chr11:32417910G>T | S381* | 0.506 | 0.525 |
| WT1 | chr11:32456243_32456244 insAA | X216_splice | 0.440 | 0.519 |
| NOTCH2 | chr1:120539668T>A | T235S | 0.000 | 0.023 |
| BCORL1 | chr23:129190051C>G | Y1692* | 0.542 | 0.495 |
| PALB2 | chr16:23652476C>A | M1I* | 0.479 | 0.432 |
| INPP4B | chr4:143043397G>A | X673_splice | 0.482 | 0.524 |
| MED12 | chr23:70361151_70361152 insCAGCAACACCAG | H2116_Q2119dup | 0.405 | 0.523 |
| NOTCH3 | chr19:15299048G>A | S497L | 0.478 | 0.500 |
| SLX4 | chr16:3639977G>A | A1221V | 0.478 | 0.513 |
| ASXL1 | chr20:31024704G>A | G1397S | 0.492 | 0.496 |
| SLX4 | chr16:3647692A>C | N457K | 0.492 | 0.503 |
| HLA-C | chr6:31238889T>C | R194G | 0.357 | 0.369 |
| MCL1 | chr1:150551649T>C | M120V | 0.500 | 0.481 |
| TET2 | chr4:106155185C>G | P29R | 1.000 | 1.000 |
| STAG1 | chr3:136062724C>G | Q1132H | 0.508 | 0.532 |
| SETD2 | chr3:47163422C>G | E902Q | 0.484 | 0.468 |
| CREBBP | chr16:3831230G>T | L551I | 0.545 | 0.574 |
| LATS2 | chr13:21563350G>C | P190R | 0.492 | 0.476 |
| MLH1 | chr3:37067306G>A | S406N | 0.458 | 0.442 |
| HLA-C | chr6:31238970T>A | T167S | 0.505 | 0.397 |
| PRKN | chr6:162864388C>T | R42H | 0.518 | 0.472 |
| EPHA3 | chr3:89456514A>G | I564V | 0.485 | 0.463 |
| RPS6KB2 | chr11:67196068G>T | E18* | 0.560 | 0.558 |
| TCF7L2 | chr10:114925687A>G | T589A | 0.495 | 0.467 |
| PLCG2 | chr16:81942175A>G | N571S | 0.493 | 0.506 |
| SLX4 | chr16:3650987T>C | M386V | 0.441 | 0.459 |
| SETD2 | chr3:47165569G>A | P186L | 0.492 | 0.508 |
| INHA | chr2:220439916G>A | A257T | 0.503 | 0.483 |
| EP400 | chr12:132445675G>A | V171M | 0.507 | 0.556 |
| TAP1 | chr6:32814942C>T | R708Q | 0.498 | 0.518 |
| SLX4 | chr16:3645607A>G | L671S | 0.496 | 0.428 |
| SLX4 | chr16:3656625G>A | R204C | 0.499 | 0.511 |
| ARID4B | chr1:235345346C>T | G963E | 0.509 | 0.540 |
| RECQL4 | chr8:145742799T>C | E71G | 0.459 | 0.509 |
| SMG1 | chr16:18875042C>A | A1209S | 0.498 | 0.433 |
| MGAM | chr7:141750579C>T | T907M | 0.511 | 0.486 |
| SLX4 | chr16:3640784_3640785 delinsAT | A952M | 0.485 | 0.470 |
| POLD1 | chr19:50918229G>A | R849H | 0.557 | 0.532 |
| MST1 | chr3:49724465C>G | G215R | 0.484 | 0.516 |
| SPRED1 | chr15:38545412A>T | D9V | 0.488 | 0.431 |

| | | | | |
|---|---|---|---|---|
| ZFHX3 | chr16:72828279T>A | T2768S | 0.496 | 0.499 |
| ANKRD11 | chr16:89350393T>C | K853E | 0.468 | 0.527 |
| EED | chr11:85988146G>T | W364L | 0.480 | 0.467 |
| FLT1 | chr13:28895635T>C | I1047V | 0.492 | 1.000 |
| BLM | chr15:91303387A>G | I366M | 0.479 | 0.516 |
| FANCA | chr16:89813083G>T | A1141D | 0.483 | 0.470 |
| STAT5B | chr17:40353806G>A | L772F | 0.520 | 0.514 |
| ERBB4 | chr2:212285279A>T | L1008M | 0.429 | 0.482 |
| ERBB4 | chr2:212543898T>C | M501V | 0.497 | 0.516 |
| NF1 | chr17:29559092G>T | D1067Y | 0.118 | 0.000 |
| MAP3K13 | chr3:185003303_185003304 del | X30_splice | 0.477 | 0.454 |

**Extended Data Table 4. Upregulated and downregulated genes in LRCs (Top 90)**

**a. MSK011**

| Upregulated genes | | Downregulated genes | |
|---|---|---|---|
| CLEC9A | RNF149 | S100A9 | ARID3A |
| HLA-DPA1 | NDC80 | FCN1 | CCT5 |
| HMOX1 | HIST1H4C | LYZ | B9D2 |
| HLA-DPB1 | H2AFX | TNNT1 | EPHB3 |
| CTD-2319I12.1 | PSAP | RNASE2 | PSMC1 |
| HLA-DQA1 | NAAA | LPPR3 | MTRNR2L1 |
| HLA-DRB1 | HLA-DRB6 | SCARNA22 | RNA5SP267 |
| GGTA1P | ANPEP | S100A8 | ZC3H15 |
| CD74 | NUSAP1 | VCAN | CITED4 |
| HLA-DRA | HIST2H2BF | BCCIP | FARSA |
| XCR1 | ADAM19 | CD300E | MNDA |
| HLA-DOB | CD40 | PTAFR | GSR |
| CLIC2 | S100B | C19orf59 | MFAP1 |
| HLA-DQB1 | HSPA1A | KRT81 | IRAK1 |
| CD1C | ST3GAL5 | CLEC5A | TRIM56 |
| FPR3 | CD34 | PADI4 | SRRT |
| TOP2A | CD83 | CAPG | GPATCH4 |
| LGALS2 | CLCN4 | CKAP4 | LENG8-AS1 |
| C1orf54 | CLNK | SNORA76 | SGTA |
| WDFY4 | ZSCAN18 | RP11-304L19.5 | VRK2 |
| BTLA | CIITA | APLP2 | FLNB |
| NAPSB | PTK2 | ZEB2 | TOM1 |
| CPVL | HLA-F | IL17RA | DDX54 |
| HIST1H1B | RP1-140K8.2 | RETN | HNRNPM |
| PIF1 | KIF15 | DSP | TKT |
| AURKB | ARL4C | CEBPE | GSTZ1 |
| IRF8 | VAC14-AS1 | CXorf21 | HSPBP1 |
| TCEA3 | ZNF462 | SLC16A3 | CHMP4A |
| ST14 | GNAI1 | MPO | ERP29 |
| KIFC1 | TNFRSF12A | CYCS | TOLLIP |
| S100A10 | FLT3 | FOSL2 | GMIP |
| CDCA2 | APOL3 | SERPINB8 | MRPL28 |
| NET1 | PRC1 | FAM101B | RASGRP4 |
| ASPM | CKAP2L | LRG1 | RBKS |
| HLA-DMA | BCL6 | ATXN2L | ZNF318 |
| ZFP36L1 | RUFY3 | RUNDC1 | CTC-338M12.5 |
| SCN3A | FNIP2 | TMEM223 | SYNE3 |
| RAB31 | ITM2C | DHX38 | NSMF |
| KIAA1598 | HIST1H2AJ | AGTRAP | NAA15 |
| P2RY14 | PARM1 | SERF2 | MRPS26 |
| TBC1D9 | SLC46A3 | METAP2 | SNORA11 |
| CDCA3 | KIF20A | MRPL33 | VPS28 |
| PPT1 | RP1-17K7.2 | TNIP2 | CALR |
| RP11-543E8.2 | ZNF331 | TRAPPC2L | ISOC2 |
| ZNF366 | FGL2 | DKC1 | MBD3 |

**b. MSK162**

| Upregulated genes | | Downregulated genes | |
|---|---|---|---|
| S100A9 | MARCKS | TRAJ29 | TWIST1 |
| PSAP | ADRBK1 | CPSF7 | SEPTIN6 |
| MAFB | DUSP6 | NT5C2 | RNF165 |
| SRGN | SERPINB1 | TRBJ2-2 | LDHA |
| LYZ | ANXA1 | TRDJ1 | ALG14 |
| GRN | MS4A6A | TRDJ3 | LINC00152 |
| RNASE2 | FCGRT | TRDJ2 | POMP |
| VCAN | NCOA4 | FAM102A | HSPH1 |
| BTG1 | PABPC1 | TRBJ2-5 | NFATC4 |
| RNU4ATAC | IER2 | RANBP1 | HAUS8 |
| FGL2 | MBNL1-AS1 | TRDJ4 | CTD-2026K11.2 |
| CTSG | MS4A3 | TMTC2 | ALB |
| MPO | MEGF9 | IGHJ2 | PES1 |
| RET | QSOX1 | TRBJ2-7 | SEL1L3 |
| TXNIP | DEFB1 | CTD-2540F13.2 | SKA2 |
| MT-TL1 | BAG6 | NRAS | RP4-666F24.3 |
| MT-RNR2 | FRMD3 | TRBJ2-3 | PROK2 |
| ATP8B4 | P4HA1 | TRBJ2-6 | UQCRH |
| LYST | GOLGB1 | AIM1 | RPL18AP3 |
| HLA-B | CDC42EP3 | BCL2 | CDKN3 |
| HLA-A | LAPTM5 | DGKI | CLSPN |
| AC010970.2 | NDRG1 | TTL | GNRHR2 |
| RETN | CSF1R | SLCO4A1 | EZH2 |
| HCST | MTRNR2L12 | TREX1 | FAM110B |
| CECR1 | AP1S2 | SLC1A5 | PA2G4 |
| S100A8 | ADAP2 | HRK | AP003471.1 |
| FRAT2 | H1FX | STMN1 | GLUL |
| RXFP2 | PBXIP1 | MRAS | ZCCHC2 |
| BANK1 | CPEB2 | BRSK1 | EBNA1BP2 |
| CFD | FABP4 | TTTY15 | CDR2L |
| FLNA | CEBPB | ATM | GPT2 |
| ZFP36L1 | FAUP1 | MGLL | NAV3 |
| PLEK | SH3BGRL | ERH | RNU6-758P |
| HOMER3 | HLA-E | CNDP2 | DBI |
| MS4A7 | ZNF503 | IRAK1 | RBM5 |
| TKT | CPM | SLC16A1 | C18orf54 |
| SLC26A7 | NPC2 | MKI67 | RP11-304L19.5 |
| TYROBP | NFAM1 | ATP5A1 | BCAT1 |
| CTSS | B2M | SSRP1 | RP1-20N2.6 |
| PDK4 | CAST | TIMM13 | MCM4 |
| EMB | ANO10 | YARS | GINS2 |
| RP11-301G19.1 | CD70 | CDCA7 | PHF19 |
| CTSH | FTL | AC007969.5 | RP5-1119A7.17 |
| DMXL2 | GABBR1 | FBXL19-AS1 | CCDC38 |
| SCARNA3 | HSP90B1 | POU2AF1 | SF3B3 |

**c. MSK165**

| Upregulated genes | | Downregulated genes | |
|---|---|---|---|
| SPTA1 | TIMP3 | SAMHD1 | MRPL33 |
| ITGA2B | GAS5 | BZRAP1 | IL17RA |
| SNORD17 | HIST1H1E | ITGB2 | NEGR1 |
| GATA2 | IL1B | IL2RA | IQGAP1 |
| EGR1 | SUCO | FGL2 | CST3 |
| ELANE | HIST1H2AB | CXorf21 | GLIPR1 |
| CFH | TFPI | CLSTN2 | RAB31 |
| TCN1 | IL7R | MYBPH | EMILIN2 |
| HDC | SNORD89 | PI15 | PHACTR3 |
| CST7 | BPI | IL6R | DTX4 |
| FCER1A | RRN3 | LGALS1 | WDR49 |
| SLC40A1 | HIST1H2BF | AIM1 | CLU |
| HIST1H4C | HIST1H2BC | KIAA0930 | ATP5O |
| RNASE3 | IER2 | FLT3 | SNRPB2 |
| MINPP1 | EIF4G3 | IRF8 | DOCK1 |
| HIST2H2BF | RP11-84C10.2 | CCR2 | TRIM25 |
| NPR3 | TKT | AHNAK | MEF2D |
| CEACAM6 | SNORA12 | QSOX1 | PPP2R2B |
| HIST1H1C | LNPEP | TNFSF13B | LAT2 |
| FOSB | PROK2 | CYBB | BRI3BP |
| GFI1B | ARG1 | DPYSL2 | LPPR3 |
| PRKAR2B | FRY | PRTN3 | SCARA3 |
| PLXNC1 | SAT1 | SH3KBP1 | LAMP5 |
| ABLIM1 | JUN | MYOF | SLC8A1 |
| TTC27 | LIMS1 | IRX3 | CBX6 |
| S100A9 | EPX | HNMT | NRIP3 |
| ZMYND8 | CDK1 | NCF2 | P2RY6 |
| SCARNA9 | CD226 | RNASE6 | DHRS9 |
| RP11-85G21.3 | TPM1 | LYZ | FH |
| ANXA1 | FBN1 | MS4A6A | TUBA4A |
| CD36 | ICAM3 | ITGB2-AS1 | IL17RE |
| GCSAML | MYCN | CAPG | APAF1 |
| VCL | SELP | MT2A | BCAT1 |
| PDZD8 | RNASE2 | ABCA6 | TFAM |
| MT-TL1 | CLEC5A | PIM1 | PRDM8 |
| XK | DACH1 | OAS2 | SELPLG |
| LTBP1 | FOXL2 | CD93 | AFF3 |
| CALD1 | RP11-354E11.2 | LY86 | GRWD1 |
| ATP6V0A2 | TPM4 | COX6C | RP11-359N11.2 |
| BMP2K | HIST1H2AJ | ADAM28 | RGS10 |
| HIST2H2AC | STXBP5 | RAD23B | TDRD7 |
| ANGPT1 | THRB | ITSN1 | CTSZ |
| TAL1 | PDLIM5 | GAS7 | TNFRSF1B |
| GATA1 | CEBPE | CLECL1 | RRP12 |
| UPP2 | CAV2 | SLC38A1 | DANCR |

**Extended Data Table 5. Commonly upregulated and downregulated genes in LRCs**

**a. Commonly upregulated genes in LRCs**

| | | | | | |
|---|---|---|---|---|---|
| **MSK011&MSK162&MSK165** | PNRC1 | ZFP36L1 | | | |
| **MSK011&MSK162** | RAB31 | PSAP | FGL2 | PPM1M | JUNB |
| | CECR1 | SIGLEC1 | FTL | LGALS3 | B2M |
| | MS4A6A | SMIM14 | | | |
| **MSK162&MSK165** | EGR1 | HIST1H1C | S100A9 | SCARNA9 | ANXA1 |
| | CD36 | MT-TL1 | IER2 | TKT | SNORA12 |
| | SAT1 | RNASE2 | AOAH | NRIP1 | BTG2 |
| | TP53INP1 | S100A8 | S100P | CA2 | RXFP2 |
| | CKAP2 | ZMAT3 | FTH1P16 | | |
| **MSK011&MSK165** | HIST1H4C | HIST2H2BF | ANGPT1 | GAS5 | CDK1 |
| | HIST1H2AJ | CDCA3 | CASC5 | NUSAP1 | ANPEP |
| | NUF2 | ALDH2 | HJURP | DUSP1 | MYLK |
| | ANLN | HLA-DOA | ECT2 | CCNA2 | |

**b. Commonly downregulated genes in LRCs**

| | | | | | |
|---|---|---|---|---|---|
| **MSK011&MSK162&MSK165** | LGALS1 | DUSP3 | RP11-304L19.5 | | |
| **MSK011&MSK162** | TRAPPC2L | IRAK1 | C7orf50 | WDR4 | TIMM13 |
| | NDUFS6 | ALOX5AP | PSMA4 | VPS18 | SMARCA4 |
| | TUFM | MRPS18B | STIP1 | PES1 | CCDC43 |
| | AP3D1 | | | | |
| **MSK162&MSK165** | AIM1 | BCAT1 | AFF3 | DANCR | GINS2 |
| | MDFI | SFXN4 | GPT2 | NAV3 | |
| **MSK011&MSK165** | CXorf21 | IL6R | LYZ | CAPG | MRPL33 |
| | IL17RA | DTX4 | MEF2D | LPPR3 | NRIP3 |
| | APAF1 | GRWD1 | CTSZ | RRP12 | TESC |
| | MX1 | POU2F2 | TNS3 | MRPL12 | FOSL2 |
| | RASSF4 | YDJC | TNNT1 | MRPS26 | TIMM21 |
| | METTL9 | | | | |

#### Extended Data Table 6. Commonly dysregulated GSEA gene sets in LRCs

##### a. Positively correlated gene sets

| | |
|---|---|
| STK33_UP | GO_LEUKOCYTE_CELL_CELL_ADHESION |
| CHANG_CORE_SERUM_RESPONSE_DN | SMIRNOV_RESPONSE_TO_IR_6HR_DN |
| WANG_RESPONSE_TO_GSK3_INHIBITOR_SB216763_UP | MONNIER_POSTRADIATION_TUMOR_ESCAPE_DN |
| MTOR_UP.N4.V1_DN | NAGASHIMA_NRG1_SIGNALING_UP |
| GSE19923_WT_VS_HEB_AND_E2A_KO_DP_THYMOCYTE_DN | HIRSCH_CELLULAR_TRANSFORMATION_SIGNATURE_UP |
| GSE39556_CD8A_DC_VS_NK_CELL_MOUSE_3H_POST_POLYIC_INJ_DN | TARTE_PLASMA_CELL_VS_PLASMABLAST_UP |
| GSE22886_NAIVE_CD4_TCELL_VS_48H_ACT_TH1_UP | GO_POSITIVE_REGULATION_OF_CYTOKINE_PRODUCTION |
| PODAR_RESPONSE_TO_ADAPHOSTIN_UP | MODULE_38 |
| GSE29618_BCELL_VS_PDC_UP | MEK_UP.V1_DN |
| TAKEDA_TARGETS_OF_NUP98_HOXA9_FUSION_8D_DN | GO_SINGLE_ORGANISM_CELL_ADHESION |
| STK33_SKM_UP | BOQUEST_STEM_CELL_CULTURED_VS_FRESH_UP |
| HALLMARK_TNFA_SIGNALING_VIA_NFKB | HOEBEKE_LYMPHOID_STEM_CELL_DN |
| PICCALUGA_ANGIOIMMUNOBLASTIC_LYMPHOMA_DN | GSE8515_IL1_VS_IL6_4H_STIM_MAC_UP |
| GSE37605_TREG_VS_TCONV_NOD_FOXP3_FUSION_GFP_UP | GSE41087_WT_VS_FOXP3_MUT_ANTI_CD3_CD28_STIM_CD4_TCELL_UP |
| KINSEY_TARGETS_OF_EWSR1_FLII_FUSION_DN | DACOSTA_UV_RESPONSE_VIA_ERCC3_COMMON_DN |
| JECHLINGER_EPITHELIAL_TO_MESENCHYMAL_TRANSITION_DN | CREIGHTON_ENDOCRINE_THERAPY_RESISTANCE_5 |
| GSE39556_UNTREATED_VS_3H_POLYIC_INJ_MOUSE_NK_CELL_DN | GSE17301_CTRL_VS_48H_IFNA2_STIM_CD8_TCELL_UP |
| GSE22601_IMMATURE_CD4_SINGLE_POSITIVE_VS_CD8_SINGLE_POSITIVE_THYMOCYTE_UP | GSE17301_CTRL_VS_48H_ACD3_ACD28_IFNA5_STIM_CD8_TCELL_DN |
| TONKS_TARGETS_OF_RUNX1_RUNX1T1_FUSION_HSC_UP | MODULE_60 |
| TAKEDA_TARGETS_OF_NUP98_HOXA9_FUSION_3D_UP | GSE9509_10MIN_VS_30MIN_LPS_STIM_IL10_KO_MACROPHAGE_DN |
| GSE33292_WT_VS_TCF1_KO_DN3_THYMOCYTE_UP | TORCHIA_TARGETS_OF_EWSR1_FLI1_FUSION_DN |
| GSE3982_BASOPHIL_VS_TH1_UP | TGCACTT_MIR519C_MIR519B_MIR519A |
| SARRIO_EPITHELIAL_MESENCHYMAL_TRANSITION_DN | GATA1_04 |
| DUTERTRE_ESTRADIOL_RESPONSE_24HR_DN | SWEET_LUNG_CANCER_KRAS_DN |
| RUTELLA_RESPONSE_TO_HGF_DN | GSE24142_ADULT_VS_FETAL_DN2_THYMOCYTE_UP |
| GSE39556_UNTREATED_VS_3H_POLYIC_INJ_MOUSE_CD8A_DC_DN | MODULE_12 |
| HALLMARK_APOPTOSIS | GSE45365_NK_CELL_VS_BCELL_UP |
| GSE14769_UNSTIM_VS_40MIN_LPS_BMDM_DN | SENESE_HDAC3_TARGETS_UP |
| GSE21927_C26GM_VS_4T1_TUMOR_MONOCYTE_BALBC_DN | EGFR_UP.V1_DN |
| NAGASHIMA_EGF_SIGNALING_UP | DAVICIONI_MOLECULAR_ARMS_VS_ERMS_UP |
| GSE36476_CTRL_VS_TSST_ACT_72H_MEMORY_CD4_TCELL_YOUNG_UP | GO_RESPONSE_TO_ACID_CHEMICAL |

| | |
|---|---|
| GSE37532_TREG_VS_TCONV_PPARG_KO_CD4_TCELL_FROM_VISCERAL_ADIPOSE_TISSUE_DN | GSE36888_UNTREATED_VS_IL2_TREATED_TCELL_2H_UP |
| GSE3982_BASOPHIL_VS_TH2_UP | GSE13493_DP_VS_CD4INTCD8POS_THYMOCYTE_UP |
| GSE3982_MAC_VS_TH1_UP | MODULE_24 |
| GSE15930_NAIVE_VS_48H_IN_VITRO_STIM_IFNAB_CD8_TCELL_UP | WANG_SMARCE1_TARGETS_UP |
| MITSIADES_RESPONSE_TO_APLIDIN_UP | AMUNDSON_RESPONSE_TO_ARSENITE |
| GSE42021_CD24HI_TREG_VS_CD24HI_TCONV_THYMUS_DN | CREB_Q4 |
| KEEN_RESPONSE_TO_ROSIGLITAZONE_DN | GO_WOUND_HEALING |
| GAL_LEUKEMIC_STEM_CELL_DN | HADDAD_B_LYMPHOCYTE_PROGENITOR |
| AMIT_EGF_RESPONSE_40_HELA | SENGUPTA_NASOPHARYNGEAL_CARCINOMA_WITH_LMP1_UP |
| WELCSH_BRCA1_TARGETS_UP | GO_POSITIVE_REGULATION_OF_CELL_ADHESION |
| GSE32986_UNSTIM_VS_CURDLAN_HIGHDOSE_STIM_DC_DN | RODRIGUES_THYROID_CARCINOMA_ANAPLASTIC_DN |
| GSE18893_TCONV_VS_TREG_2H_TNF_STIM_DN | ACATTCC_MIR1_MIR206 |
| GEORGANTAS_HSC_MARKERS | BASAKI_YBX1_TARGETS_DN |
| GSE36476_CTRL_VS_TSST_ACT_40H_MEMORY_CD4_TCELL_YOUNG_UP | GO_REGULATION_OF_CELL_ACTIVATION |
| GSE27434_WT_VS_DNMT1_KO_TREG_DN | GO_RESPONSE_TO_OXIDATIVE_STRESS |
| KOBAYASHI_EGFR_SIGNALING_24HR_UP | GO_REGULATION_OF_BODY_FLUID_LEVELS |
| GSE24142_DN2_VS_DN3_THYMOCYTE_UP | AACTGGA_MIR145 |
| ODONNELL_TFRC_TARGETS_UP | GO_RESPONSE_TO_INORGANIC_SUBSTANCE |
| STK33_NOMO_UP | CAGTATT_MIR200B_MIR200C_MIR429 |
| PHONG_TNF_TARGETS_UP | WTTGKCTG_UNKNOWN |
| GSE26156_DOUBLE_POSITIVE_VS_CD4_SINGLE_POSITIVE_THYMOCYTE_UP | GSE22229_UNTREATED_VS_IMMUNOSUPP_THERAPY_RENAL_TRANSPLANT_PATIENT_PBMC_DN |
| GSE28726_ACT_CD4_TCELL_VS_ACT_VA24NEG_NKTCELL_DN | GSE21670_IL6_VS_TGFB_AND_IL6_TREATED_STAT3_KO_CD4_TCELL_DN |
| GSE22432_PDC_VS_TGFB1_TREATEDCOMMON_DC_PROGENITOR_DN | GNF2_PCAF |
| GERY_CEBP_TARGETS | GSE20366_EX_VIVO_VS_DEC205_CONVERSION_UP |
| GSE21360_SECONDARY_VS_QUATERNARY_MEMORY_CD8_TCELL_UP | WIERENGA_STAT5A_TARGETS_GROUP2 |
| GSE7219_WT_VS_NIK_NFKB2_KO_DC_UP | GSE23925_DARK_ZONE_VS_NAIVE_BCELL_DN |
| GSE29618_BCELL_VS_MDC_UP | GSE31082_CD4_VS_CD8_SP_THYMOCYTE_UP |
| MODULE_5 | GO_REGULATION_OF_EXTRINSIC_APOPTOTIC_SIGNALING_PATHWAY |
| GSE42021_TREG_PLN_VS_CD24HI_TREG_THYMUS_UP | TGAATGT_MIR181A_MIR181B_MIR181C_MIR181D |
| GSE369_PRE_VS_POST_IL6_INJECTION_SOCS3_KO_LIVER_UP | RAGHAVACHARI_PLATELET_SPECIFIC_GENES |
| GSE7460_CD8_TCELL_VS_TREG_ACT_UP | GSE37301_MULTIPOTENT_PROGENITOR_VS_PRO_BCELL_DN |
| GSE12366_GC_VS_MEMORY_BCELL_DN | GSE22886_UNSTIM_VS_STIM_MEMORY_TCELL_UP |
| ZHENG_BOUND_BY_FOXP3 | GSE22935_UNSTIM_VS_24H_MBOVIS_BCG_STIM_MYD88_KO_MACROPHAGE_UP |
| GSE24574_BCL6_HIGH_VS_LOW_TFH_CD4_TCELL_UP | PENG_RAPAMYCIN_RESPONSE_UP |

| | |
|---|---|
| HECKER_IFNB1_TARGETS | GSE21774_CD62L_POS_CD56_BRIGHT_VS_CD62L_NEG_CD56_DIM_NK_CELL_DN |
| GSE41867_NAIVE_VS_DAY8_LCMV_EFFECTOR_CD8_TCELL_DN | GO_RESPONSE_TO_ALCOHOL |
| GSE23505_UNTREATED_VS_4DAY_IL6_IL1_IL23_TREATED_CD4_TCELL_DN | GSE24634_NAIVE_CD4_TCELL_VS_DAY3_IL4_CONV_TREG_UP |
| GO_ENDOSOMAL_PART | GO_LEUKOCYTE_DIFFERENTIATION |
| HORIUCHI_WTAP_TARGETS_UP | DAZARD_RESPONSE_TO_UV_NHEK_DN |
| TIEN_INTESTINE_PROBIOTICS_24HR_DN | GSE32986_GMCSF_VS_GMCSF_AND_CURDLAN_LOWDOSE_STIM_DC_UP |
| GSE17974_CTRL_VS_ACT_IL4_AND_ANTI_IL12_48H_CD4_TCELL_UP | GSE21033_CTRL_VS_POLYIC_STIM_DC_12H_UP |
| AMIT_SERUM_RESPONSE_60_MCF10A | GSE9988_ANTI_TREM1_VS_LOW_LPS_MONOCYTE_DN |
| GSE28408_LY6G_POS_VS_NEG_DC_UP | PILON_KLF1_TARGETS_UP |
| WIERENGA_STAT5A_TARGETS_DN | ADDYA_ERYTHROID_DIFFERENTIATION_BY_HEMIN |
| GSE36392_TYPE_2_MYELOID_VS_NEUTROPHIL_IL25_TREATED_LUNG_UP | GSE17974_0H_VS_72H_IN_VITRO_ACT_CD4_TCELL_UP |
| GSE36476_CTRL_VS_TSST_ACT_16H_MEMORY_CD4_TCELL_YOUNG_UP | GSE7852_THYMUS_VS_FAT_TCONV_UP |
| CHYLA_CBFA2T3_TARGETS_UP | GSE40274_CTRL_VS_FOXP3_AND_EOS_TRANSDUCED_ACTIVATED_CD4_TCELL_UP |
| GSE40666_UNTREATED_VS_IFNA_STIM_STAT4_KO_EFFECTOR_CD8_TCELL_90MIN_DN | GO_RESPONSE_TO_METAL_ION |
| GSE2770_TGFB_AND_IL4_VS_TGFB_AND_IL12_TREATED_ACT_CD4_TCELL_6H_DN | LU_EZH2_TARGETS_DN |
| HELLER_HDAC_TARGETS_UP | GO_POSITIVE_REGULATION_OF_LOCOMOTION |
| HELLER_HDAC_TARGETS_SILENCED_BY_METHYLATION_UP | MODULE_64 |
| GSE24574_BCL6_HIGH_TFH_VS_TFH_CD4_TCELL_DN | TAATAAT_MIR126 |
| GSE39820_TGFBETA3_IL6_VS_TGFBETA3_IL6_IL23A_TREATED_CD4_TCELL_UP | GSE23502_WT_VS_HDC_KO_MYELOID_DERIVED_SUPPRESSOR_CELL_BM_UP |
| GO_VESICLE_MEMBRANE | PDGF_ERK_DN.V1_DN |
| GSE24726_WT_VS_E2-2_KO_PDC_DAY6_POST_DELETION_DN | GSE43956_WT_VS_SGK1_KO_TH17_DIFFERENTIATED_CD4_TCELL_DN |
| GSE25123_CTRL_VS_IL4_AND_ROSIGLITAZONE_STIM_PPARG_KO_MACROPHAGE_UP | SRF_Q4 |
| GSE14769_UNSTIM_VS_20MIN_LPS_BMDM_DN | GSE9509_LPS_VS_LPS_AND_IL10_STIM_IL10_KO_MACROPHAGE_30MIN_UP |
| DIRMEIER_LMP1_RESPONSE_EARLY | GSE17974_CTRL_VS_ACT_IL4_AND_ANTI_IL12_72H_CD4_TCELL_UP |
| GSE21546_UNSTIM_VS_ANTI_CD3_STIM_DP_THYMOCYTES_UP | MODULE_2 |
| GO_REGULATION_OF_RESPONSE_TO_WOUNDING | GSE9006_HEALTHY_VS_TYPE_1_DIABETES_PBMC_AT_DX_DN |
| FOSTER_TOLERANT_MACROPHAGE_DN | DEBIASI_APOPTOSIS_BY_REOVIRUS_INFECTION_UP |
| GSE42088_UNINF_VS_LEISHMANIA_INF_DC_4H_DN | MANALO_HYPOXIA_UP |
| GSE45739_NRAS_KO_VS_WT_UNSTIM_CD4_TCELL_DN | GSE4590_LARGE_PRE_BCELL_VS_VPREB_POS_LARGE_PRE_BCELL_UP |
| GSE36476_CTRL_VS_TSST_ACT_72H_MEMORY_CD4_TCELL_OLD_UP | GO_SIDE_OF_MEMBRANE |
| GSE23925_LIGHT_ZONE_VS_NAIVE_BCELL_UP | GO_ORGANELLE_LOCALIZATION |
| GSE46606_UNSTIM_VS_CD40L_IL2_IL5_1DAY_STIMULATED_IRF4HIGH_SORTED_BCELL_DN | GSE20727_H2O2_VS_ROS_INHIBITOR_TREATED_DC_UP |
| GSE21546_WT_VS_SAP1A_KO_AND_ELK1_KO_ANTI_CD3_STIM_DP_THYMOCYTES_UP | GSE2706_UNSTIM_VS_2H_R848_DC_DN |

| | |
|---|---|
| PHONG_TNF_RESPONSE_VIA_P38_PARTIAL | CREB_Q2 |
| GSE6259_BCELL_VS_CD8_TCELL_DN | GSE20727_DNFB_ALLERGEN_VS_ROS_INH_AND_DNFB_ALLERGEN_TREATED_DC_UP |
| HALLMARK_HEME_METABOLISM | DAZARD_RESPONSE_TO_UV_NHEK_UP |
| GSE16385_ROSIGLITAZONE_IL4_VS_IL4_ALONE_STIM_MACROPHAGE_12H_DN | MODULE_19 |
| GSE3039_CD4_TCELL_VS_B2_BCELL_UP | GSE16266_CTRL_VS_LPS_STIM_MEF_UP |
| IVANOVA_HEMATOPOIESIS_MATURE_CELL | GSE25123_WT_VS_PPARG_KO_MACROPHAGE_IL4_AND_ROSIGLITAZONE_STIM_DN |
| GO_NEGATIVE_REGULATION_OF_IMMUNE_SYSTEM_PROCESS | TAKEDA_TARGETS_OF_NUP98_HOXA9_FUSION_10D_DN |
| ZWANG_CLASS_1_TRANSIENTLY_INDUCED_BY_EGF | GSE12845_IGD_POS_BLOOD_VS_DARKZONE_GC_TONSIL_BCELL_UP |
| GARY_CD5_TARGETS_UP | KAECH_DAY8_EFF_VS_MEMORY_CD8_TCELL_DN |
| GSE27859_CD11C_INT_F480_HI_MACROPHAGE_VS_CD11C_ING_F480_INT_DC_DN | GSE8515_CTRL_VS_IL1_4H_STIM_MAC_UP |
| GO_ORGANELLE_SUBCOMPARTMENT | CHEN_HOXA5_TARGETS_9HR_UP |
| GSE21774_CD62L_POS_CD56_DIM_VS_CD62L_NEG_CD56_DIM_NK_CELL_UP | GO_DNA_PACKAGING_COMPLEX |
| MODULE_6 | GO_CELLULAR_RESPONSE_TO_OXIDATIVE_STRESS |
| OSWALD_HEMATOPOIETIC_STEM_CELL_IN_COLLAGEN_GEL_UP | GSE21546_WT_VS_SAP1A_KO_AND_ELK1_KO_DP_THYMOCYTES_DN |
| TBK1.DF_UP | GO_T_CELL_DIFFERENTIATION |
| GO_REGULATION_OF_IMMUNE_EFFECTOR_PROCESS | VERHAAK_AML_WITH_NPM1_MUTATED_DN |
| MODULE_1 | GNF2_BNIP3L |
| GSE40068_BCL6_POS_VS_NEG_CXCR5_POS_TFH_DN | VANHARANTA_UTERINE_FIBROID_DN |
| GSE9988_ANTI_TREM1_AND_LPS_VS_CTRL_TREATED_MONOCYTES_UP | GSE29617_CTRL_VS_DAY7_TIV_FLU_VACCINE_PBMC_2008_UP |
| GSE3039_NKT_CELL_VS_B2_BCELL_UP | WINTER_HYPOXIA_METAGENE |
| GSE16450_IMMATURE_VS_MATURE_NEURON_CELL_LINE_12H_IFNA_STIM_UP | CHEN_LVAD_SUPPORT_OF_FAILING_HEART_UP |
| GSE40277_EOS_AND_LEF1_TRANSDUCED_VS_CTRL_CD4_TCELL_UP | YAGI_AML_WITH_T_8_21_TRANSLOCATION |
| RIGGI_EWING_SARCOMA_PROGENITOR_UP | KIM_WT1_TARGETS_UP |
| CHR6P22 | ENK_UV_RESPONSE_EPIDERMIS_DN |
| WIERENGA_STAT5A_TARGETS_UP | DORN_ADENOVIRUS_INFECTION_24HR_DN |
| GSE6674_ANTI_IGM_VS_CPG_STIM_BCELL_UP | GSE9988_ANTI_TREM1_VS_CTRL_TREATED_MONOCYTES_UP |
| GO_RESPONSE_TO_PEPTIDE | GSE369_SOCS3_KO_VS_WT_LIVER_POST_IL6_INJECTION_DN |
| GSE17974_CTRL_VS_ACT_IL4_AND_ANTI_IL12_4H_CD4_TCELL_UP | GSE3982_BCELL_VS_TH2_UP |
| ZWANG_CLASS_3_TRANSIENTLY_INDUCED_BY_EGF | GO_REGULATION_OF_CYTOKINE_PRODUCTION_INVOLVED_IN_IMMUNE_RESPONSE |
| TTTGCAC_MIR19A_MIR19B | GO_REPRODUCTIVE_SYSTEM_DEVELOPMENT |
| GSE24972_MARGINAL_ZONE_BCELL_VS_FOLLICULAR_BCELL_IRF8_KO_DN | GSE4984_LPS_VS_VEHICLE_CTRL_TREATED_DC_DN |
| GSE3982_BASOPHIL_VS_EFF_MEMORY_CD4_TCELL_UP | YAGI_AML_FAB_MARKERS |

#### b. Negative correlated gene sets

| | |
|---|---|
| GO_ORGANELLAR_RIBOSOME | MORF_PTPN11 |
| MORF_DNMT1 | GO_NCRNA_PROCESSING |
| GO_MITOCHONDRIAL_TRANSLATION | MORF_GSPT1 |
| GO_TRANSLATIONAL_TERMINATION | MORF_HDAC1 |
| MORF_FBL | GSE22886_IL2_VS_IL15_STIM_NKCELL_DN |
| MORF_BUB3 | MORF_RAD21 |
| GO_CELLULAR_PROTEIN_COMPLEX_DISASSEMBLY | PENG_RAPAMYCIN_RESPONSE_DN |
| GO_TRANSLATIONAL_ELONGATION | MORF_DAP3 |
| MORF_RAN | GO_RIBONUCLEOPROTEIN_COMPLEX_BIOGENESIS |
| MORF_G22P1 | MORF_GNB1 |
| GSE21927_SPLEEN_C57BL6_VS_EL4_TUMOR_BALBC_MONOCYTES_DN | WELCSH_BRCA1_TARGETS_DN |
| MORF_EIF3S2 | GO_RNA_SPLICING_VIA_TRANSESTERIFICATION_REACTIONS |
| GSE24634_TREG_VS_TCONV_POST_DAY3_IL4_CONVERSION_UP | GSE27859_DC_VS_CD11C_INT_F480_INT_DC_DN |
| GSE16450_IMMATURE_VS_MATURE_NEURON_CELL_LINE_DN | GSE24210_CTRL_VS_IL35_TREATED_TCONV_CD4_TCELL_DN |
| MORF_SOD1 | MORF_UBE2I |
| MORF_RAD23A | MORF_RAD54L |
| HALLMARK_MYC_TARGETS_V2 | GSE24210_RESTING_TREG_VS_TCONV_DN |
| GSE43863_NAIVE_VS_MEMORY_LY6C_INT_CXCR5POS_CD4_TCELL_D150_LCMV_DN | STARK_PREFRONTAL_CORTEX_22Q11_DELETION_DN |
| MORF_PCNA | MORF_PSMC1 |
| MORF_CSNK2B | GSE40274_XBP1_VS_FOXP3_AND_XBP1_TRANSDUCED_ACTIVATED_CD4_TCELL_DN |
| MORF_DEK | GSE41867_DAY6_VS_DAY8_LCMV_ARMSTRONG_EFFECTOR_CD8_TCELL_DN |
| REACTOME_MRNA_SPLICING | GO_GENERATION_OF_PRECURSOR_METABOLITES_AND_ENERGY |
| MORF_ANP32B | MORF_CTBP1 |
| MORF_SMC1L1 | MORF_ACP1 |
| MORF_MSH2 | GO_RRNA_METABOLIC_PROCESS |
| GO_ORGANELLE_INNER_MEMBRANE | PENG_LEUCINE_DEPRIVATION_DN |
| GO_NUCLEOLAR_PART | GSE14415_TCONV_VS_FOXP3_KO_INDUCED_TREG_UP |
| MORF_HDAC2 | GO_CELLULAR_COMPONENT_DISASSEMBLY |
| GO_MITOCHONDRIAL_MATRIX | MORF_UBE2N |
| GSE37532_TREG_VS_TCONV_CD4_TCELL_FROM_LN_UP | MORF_SKP1A |
| MORF_AATF | MORF_GMPS |
| GSE40274_CTRL_VS_FOXP3_AND_XBP1_TRANSDUCED_ACTIVATED_CD4_TCELL_UP | GO_MATURATION_OF_5_8S_RRNA |
| GO_RIBOSOMAL_LARGE_SUBUNIT_BIOGENESIS | GO_RIBONUCLEOPROTEIN_COMPLEX_SUBUNIT_ORGANIZATION |
| MORF_HAT1 | GSE41867_LCMV_ARMSTRONG_VS_CLONE13_DAY6_EFFECTOR_CD8_TCELL_DN |
| YAO_TEMPORAL_RESPONSE_TO_PROGESTERONE_CLUSTER_14 | MONNIER_POSTRADIATION_TUMOR_ESCAPE_UP |
| GO_PRERIBOSOME | IVANOVA_HEMATOPOIESIS_INTERMEDIATE_PROGENITOR |
| GSE21927_BALBC_VS_C57BL6_MONOCYTE_TUMOR_UP | GSE30971_2H_VS_4H_LPS_STIM_MACROPHAGE_WBP7_HET_UP |
| MORF_XRCC5 | KIM_ALL_DISORDERS_DURATION_CORR_DN |
| MORF_MAP2K2 | GSE6674_ANTI_IGM_VS_CPG_STIM_BCELL_DN |

| | |
|---|---|
| MORF_PPP1CA | GSE43863_TH1_VS_LY6C_INT_CXCR5POS_MEMORY_CD4_TCELL_UP |
| GO_RIBOSOME_BIOGENESIS | GO_MATURATION_OF_5_8S_RRNA_FROM_TRICISTRONIC_RRNA_TRANSCRIPT_SSU_RRNA_5_8S_RRNA_LSU_RRNA_ |
| GO_90S_PRERIBOSOME | MORF_MTA1 |
| PENG_GLUTAMINE_DEPRIVATION_DN | MENSSEN_MYC_TARGETS |
| MORF_PRKDC | GO_RNA_SPLICING |
| REACTOME_MRNA_PROCESSING | CAMP_UP.V1_UP |
| GSE21360_SECONDARY_VS_TERTIARY_MEMORY_CD8_TCELL_UP | TONKS_TARGETS_OF_RUNX1_RUNX1T1_FUSION_MONOCYTE_UP |
| MOOTHA_MITOCHONDRIA | GSE41867_NAIVE_VS_EFFECTOR_CD8_TCELL_DN |

**Extended Data Table 7. AML patient sample information for scRNA-seq data before and after chemotherapy (GSE116256)**

| Patient ID | Age | Gender | Subtype | Somatic mutations | Sample collection post-therapy (days) | | |
|---|---|---|---|---|---|---|---|
| AML329 | 73 | F | Monocytic | FLT3, NPM1, NOTCH1, SMC3 | 0 | 20 | 37 |
| AML420B | 58 | M | Monocytic | TP53, IDH2, SH2B3 | 0 | 14 | 35 |
| AML556 | 70 | M | Monocytic | DNMT3A, NPM1, NRAS, TET2 | 0 | 15* | 31 |
| AML707B | 26 | M | Non-monocytic | BRCC3, KIT, RAD21, RUNX1-RUNX1T1 | 0 | 18 | 41 |
| AML870 | 32 | M | Non-monocytic | ZRSR2, MLL-MLLT3 | 0 | 14 | - |

**Extended Data Table 8. Plasmid map and sequences of doxycycline-inducible cDNA expressing vectors, V191. (e.g. TagBFP control)**

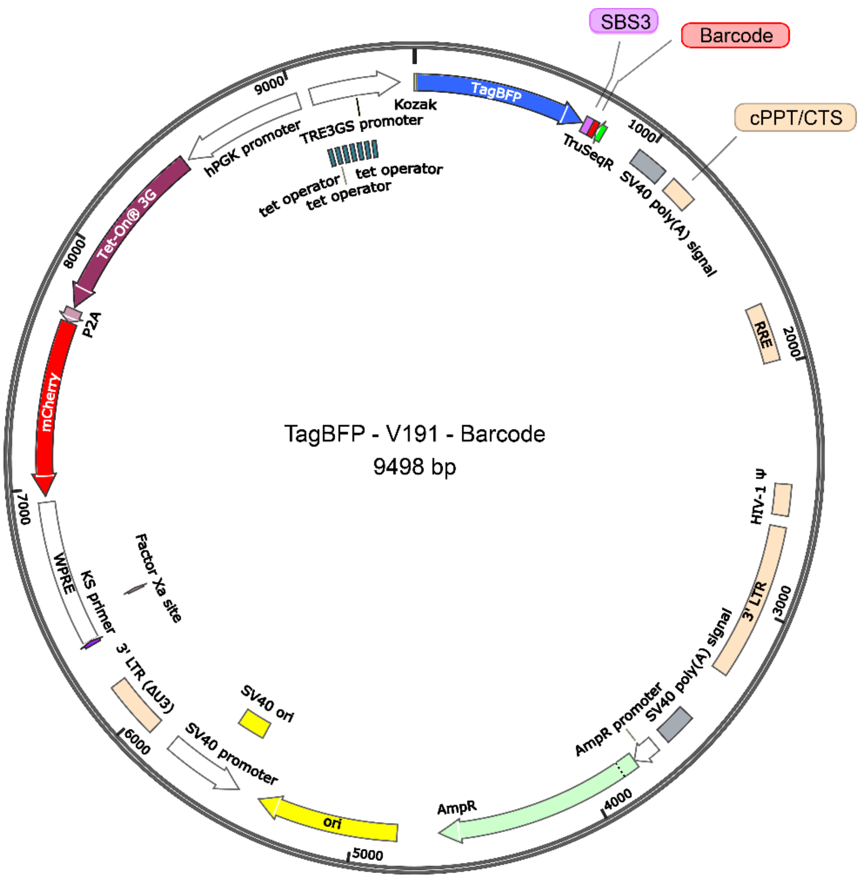


GCCGCCACCATGAGCGAGCTGATTAAGGAGAACATGCACATGAAGCTGTACATGGAGGGCACCGTG
GACAACCATCACTTCAAGTGCACATCCGAGGGCGAAGGCAAGCCCTACGAGGGCACCCAGACCATG
AGAATCAAGGTGGTCGAGGGCGGCCCTCTCCCCTTCGCCTTCGACATCCTGGCTACTAGCTTCCTC
TACGGCAGCAAGACCTTCATCAACCACACCCAGGGCATCCCCGACTTCTTCAAGCAGTCCTTCCCTG
AGGGCTTCACATGGGAGAGAGTCACCACATACGAAGACGGGGGCGTGCTGACCGCTACCCAGGAC
ACCAGCCTCCAGGACGGCTGCCTCATCTACAACGTCAAGATCAGAGGGGTGAACTTCACATCCAAC
GGCCCTGTGATGCAGAAGAAAACACTCGGCTGGGAGGCCTTCACCGAAACGCTGTACCCCGCTGAC
GGCGGCCTGGAAGGCAGAAACGACATGGCCCTGAAGCTCGTGGGCGGGAGCCATCTGATCGCAAA
CATCAAGACCACATATAGATCCAAGAAACCCGCTAAGAACCTCAAGATGCCTGGCGTCTACTATGTG
GACTACAGACTGGAAAGAATCAAGGAGGCCAACAACGAGACCTACGTCGAGCAGCACGAGGTGGCA
GTGGCCAGATACTGCGACCTCCCTAGCAAACTGGGGCACAAGCTTAATTGAACGCGTACACTCTTTC
CCTACACGACGCTCTTCCGATCTTTTCTATGATCGGTATTCTTGTGCCAGATCGGAAGAGCACACGT
CTGAACTCCAGTCACGCTAGCCTGGACAGCCAATGACGGGTAAGAGAGTGACATTTTTCACTAACCT
AAGACAGGAGGGCCGTCAGAGCTACTGCCTAATCCAAAGACGGGTAAAAGTGATAAAAATGTATCAC
TCCAACCTAAGACAGGCGCAGCTTCCGAGGGATTTGAGATCCAGACATGATAAGATACATTGATGAG
TTTGGACAAACCAAAACTAGAATGCAGTGAAAAAAATGCCTTATTTGTGAAATTTGTGATGCTATTGC
CTTATTTGTAACCATTATAAGCTGCAATAAACAAGTTAAGTCGATAAACTGGATCTCTGCTGTCCCTGT
AATAAACCCGAAAATTTTGAATTTTTGTAATTTGTTTTTGTAGTTCTTTAGTTTGTATGTCTGTTGCTAT
TATGTCTACTATTCTTTCCCCTGCACTGTACCCCCCAATCCCCCCTTTTCTTTTAAAGGCGATACCGT
CGAGATCCGTTCACTAATCGAATGGATCTGTCTCTGTCTCTCTCTCCACCTTCTTCTTCTATTCCTTCG
GGCCTGTCGGGTCCCCTCGGGGTTGGGAGGTGGGTCTGAAACGATAATGGTGAATATCCCTGCCTA

ACTCTATTCACTATAGAAAGTACAGCAAAAACTATTCTTAAACCTACCAAGCCTCCTACTATCATTATG
AATAATTTTATATACCACAGCCAATTTGTTATGTTAAACCAATTCCACAAACTTGCCCATTTATCTAATT
CCAATAATTCTTGTTCATTCTTTTCTTGCTGGTTTTGCGATTCTTCAATTAAGGAGTGTATTAAGCTTGT
GTAATTGTTAATTTCTCTGTCCCACTCCATCCAGGTCGTGTGATTCCAAATCTGTTCCAGAGATTTATT
ACTCCAACTAGCATTCCAAGGCACAGCAGTGGTGCAAATGAGTTTTCCAGAGCAACCCCAAATCCCC
AGGAGCTGTTGATCCTTTAGGTATCTTTCCACAGCCAGGATTCTTGCCTGGAGCTGCTTGATGCCCC
AGACTGTGAGTTGCAACAGATGCTGTTGCGCCTCAATAGCCCTCAGCAAATTGTTCTGCTGCTGCAC
TATACCAGACAATAATTGTCTGGCCTGTACCGTCAGCGTCATTGACGCTGCGCCCATAGTGCTTCCT
GCTGCTCCCAAGAACCCAAGGAACAAAGCTCCTATTCCCACTGCTCTTTTTTCTCTCTGCACCACTCT
TCTCTTTGCCTTGGTGGGTGCTACTCCTAATGGTTCAATTTTTACTACTTTATATTTATATAATTCACTT
CTCCAATTGTCCCTCATATCTCCTCCTCCAGGTCTGAAGATCAGCGGCCGGCCGCTTGCTGTGCGGT
GGTCTTACTTTTGTTTTGCTCTTCCTCTATCTTGTCTAAAGCTTCCTTGGTGTCTTTTATCTCTATCCTT
TGATGCACACAATAGAGGGTTGCTACTGTATTATATAATGATCTAAGTTCTTCTGATCCTGTCTGAAG
GGATGGTTGTAGCTGTCCCAGTATTTGTCTACAGCCTTCTGATGTTTCTAACAGGCCAGGATTAACTG
CGAATCGTTCTAGCTCCCTGCTTGCCCATACTATATGTTTTAATTTATATTTTTTCTTTCCCCCTGGCC
TTAACCGAATTTTTTCCCATCGCGATCTAATTCTCCCCCGCTTAATACTGACGCTCTCGCACCCATCT
CTCTCCTTCTAGCCTCCGCTAGTCAAAATTTTTGGCGTACTCACCAGTCGCCGCCCCTCGCCTCTTG
CCGTGCGCGCTTCAGCAAGCCGAGTCCTGCGTCGAGAGAGCTCCTCTGGTTTCCCTTTCGCTTTCA
AGTCCCTGTTCGGGCGCCACTGCTAGAGATTTTCCACACTGACTAAAAGGGTCTGAGGGATCTCTAG
TTACCAGAGTCACACAACAGACGGGCACACACTACTTGAAGCACTCAAGGCAAGCTTTATTGAGGCT
TAAGCAGTGGGTTCCCTAGTTAGCCAGAGAGCTCCCAGGCTCAGATCTGGTCTAACCAGAGAGACC
CAGTACAGGCAAAAAGCAGCTGCTTATATGCAGGATCTGAGGGCTCGCCACTCCCCAGTCCCGCCC
AGGCCACGCCTCCCTGGAAAGTCCCCAGCGGAAAGTCCCTTGTAGCAAGCTCGATATCAGCAGTTC
TTGAAGTACTCCGGATGCAGCTCTCGGGCCACGTGATGAAATGCTAGGCGGCTGTCAAACCTCCAC
TCTAACACTTCTCTCTCCGGGTCATCCATCCCATGCAGGCTCACAGGGTGTAACAAGCTGGTGTTCT
CTCCTTTATTGGCCTCTTCTACCTTATCTGGCTCAACTGGTACTAGCTTGTAGCACCATCCAAAGGTC
AGTGGATATCTGACCCCTGGCCCTGGTGTGTAGTTCTGCTAATCAGGGAAGTAGCCTTGTGTGTGGT
AGATCCACAGATCAAGGATATCTTGTCTTCTTTGGGAGTGAATTAGCCCTTCCAACTACTAAGTTTGT
AGTACATATTTAACAAATACAATTTCTTTAAAAACTACTAAGTTTGTAGTACATATTTAACAAATACAAT
TTCTTTAAAATGAAAATAATTCAGAGGAATCACAGGTTTAGAGTAAATGAAACCACTAGGTAATTGGC
AGTGGTAATAGGGTATGGGGTGGGAAGTTTGGGATGATTTTGGTTAGCTTGAGTTATCCAGTTGATC
CAGACATGATAAGATACATTGATGAGTTTGGACAAACCACAACTAGAATGCAGTGAAAAAAATGCTTT
ATTTGTGAAATTTGTGATGCTATTGCTTTATTTGTAACCATTATAAGCTGCAATAAACAAGTTGATCTC
CCGATCCGTCGACGTCAGGTGGCACTTTTCGGGGAAATGTGCGCGGAACCCCTATTTGTTTATTTTT
CTAAATACATTCAAATATGTATCCGCTCATGAGACAATAACCCTGATAAATGCTTCAATAATATTGAAA
AAGGAAGAGTATGAGTATTCAACATTTCCGTGTCGCCCTTATTCCCTTTTTTGCGGCATTTTGCCTTC
CTGTTTTTGCTCACCCAGAAACGCTGGTGAAAGTAAAAGATGCTGAAGATCAGTTGGGTGCACGAGT
GGGTTACATCGAACTGGATCTCAACAGCGGTAAGATCCTTGAGAGTTTTCGCCCCGAAGAACGTTTT
CCAATGATGAGCACTTTTAAAGTTCTGCTATGTGGCGCGGTATTATCCCGTATTGACGCCGGGCAAG
AGCAACTCGGTCGCCGCATACACTATTCTCAGAATGACTTGGTTGAGTACTCACCAGTCACAGAAAA
GCATCTTACGGATGGCATGACAGTAAGAGAATTATGCAGTGCTGCCATAACCATGAGTGATAACACT
GCGGCCAACTTACTTCTGACAACGATCGGAGGACCGAAGGAGCTAACCGCTTTTTTGCACAACATGG
GGGATCATGTAACTCGCCTTGATCGTTGGGAACCGGAGCTGAATGAAGCCATACCAAACGACGAGC
GTGACACCACGATGCCTGTAGCAATGGCAACAACGTTGCGCAAACTATTAACTGGCGAACTACTTAC
TCTAGCTTCCCGGCAACAATTAATAGACTGGATGGAGGCGGATAAAGTTGCAGGACCACTTCTGCGC
TCGGCCCTTCCGGCTGGCTGGTTTATTGCTGATAAATCTGGAGCCGGTGAGCGTGGGTCTCGCGGT
ATCATTGCAGCACTGGGGCCAGATGGTAAGCCCTCCCGTATCGTAGTTATCTACACGACGGGGAGT
CAGGCAACTATGGATGAACGAAATAGACAGATCGCTGAGATAGGTGCCTCACTGATTAAGCATTGGT
AACTGTCAGACCAAGTTTACTCATATATACTTTAGATTGATTTAAAACTTCATTTTTAATTTAAAAGGAT
CTAGGTGAAGATCCTTTTTGATAATCTCATGACCAAAATCCCTTAACGTGAGTTTTCGTTCCACTGAG
CGTCAGACCCCGTAGAAAAGATCAAAGGATCTTCTTGAGATCCTTTTTTTCTGCGCGTAATCTGCTGC
TTGCAAACAAAAAAACCGCTACCAGCGGTGGTTTGTTTGCCGGATCAAGAGCTACCAACTCTTTTTCC
GAAGGTAACTGGCTTCAGCAGAGCGCAGATACCAAATACTGTTCTTCTAGTGTAGCCGTAGTTAGGC
CACCACTTCAAGAACTCTGTAGCACCGCCTACATACCTCGCTCTGCTAATCCTGTTACCAGTGGCTG
CTGCCAGTGGCGATAAGTCGTGTCTTACCGGGTTGGACTCAAGACGATAGTTACCGGATAAGGCGC
AGCGGTCGGGCTGAACGGGGGGTTCGTGCACACAGCCCAGCTTGGAGCGAACGACCTACACCGAA

CTGAGATACCTACAGCGTGAGCTATGAGAAAGCGCCACGCTTCCCGAAGGGAGAAAGGCGGACAG
GTATCCGGTAAGCGGCAGGGTCGGAACAGGAGAGCGCACGAGGGAGCTTCCAGGGGGAAACGCCT
GGTATCTTTATAGTCCTGTCGGGTTTCGCCACCTCTGACTTGAGCGTCGATTTTTGTGATGCTCGTCA
GGGGGGCGGAGCCTATGGAAAAACGCCAGCAACGCGGCCTTTTTACGGTTCCTGGCCTTTTGCTGG
CCTTTTGCTCACATGTAATGGATATACAAGCTCCCGGGAGCTTTTTGCAAAAGCCTAGGCCTCCAAAA
AAGCCTCCTCACTACTTCTGGAATAGCTCAGAGGCAGAGGCGGCCTCGGCCTCTGCATAAATAAAAA
AAATTAGTCAGCCATGGGGCGGAGAATGGGCGGAACTGGGCGGAGTTAGGGGCGGGATGGGCGG
AGTTAGGGGCGGGACTATGGTTGCTGACTAATTGAGATGCATGCTTTGCATACTTCTGCCTGCTGGG
GAGCCTGGGGACTTTCCACACCTGGTTGCTGACTAATTGAGATGCATGCTTTGCATACTTCTGCCTG
CTGGGGAGCCTGGGGACTTTCCACACCCTAACTGACACACATTCCAGTTTAAACAATGTCAAGGCCT
CTCACTCTCTGATATTCATTTCTTTGCAAGTTATAAATACTGAATAATAAGATGACATGAACTACTACT
GCTAGAGATTTTCCACACTGACTAAAAGGGTCTGAGGGATCTCTAGTTACCAGAGTCACACAACAGA
CGGGCACACACTACTTGAAGCACTCAAGGCAAGCTTTATTGAGGCTTAAGCAGTGGGTTCCCTAGTT
AGCCAGAGAGCTCCCAGGCTCAGATCTGGTCTAACCAGAGAGACCCAGTACAAGCAAAAAGCAGAT
CTTGTCTTCGTTGGGAGTGAATTAGCCCTTCCAGTCCCCTCTTTTCTTTTAAAAAGTGGCTAAGATCT
ACAGCTGCCTTGTAAGTCATTGGTCTTAAAGGTACCTGAGGTGTGACTGGAAAACCCACCTCCTCCT
CCTCTTGTGCTTCTAGCCAGGCACAATCAGCATTGGTAGCTGCTGTATTGCTACTTGTGATTGCTCCA
TGTTTTTCTAGGTCTCGAGGTCGACGGTATCGATGCGGGGAGGCGGCCCAAAGGGAGATCCGACTC
GTCTGAGGGCGAAGGCGAAGACGCGGAAGAGGCCGCAGAGCCGGCAGCAGGCCGCGGGAAGGAA
GGTCCGCTGGATTGAGGGCCGAAGGGACGTAGCAGAAGGACGTCCCGCGCAGAATCCAGGTGGCA
ACACAGGCGAGCAGCCAAGGAAAGGACGATGATTTCCCCGACAACACCACGGAATTGTCAGTGCCC
AACAGCCGAGCCCCTGTCCAGCAGCGGGCAAGGCAGGCGGCGATGAGTTCCGCCGTGGCAATAGG
GAGGGGGAAAGCGAAAGTCCCGGAAAGGAGCTGACAGGTGGTGGCAATGCCCCAACCAGTGGGG
GTTGCGTCAGCAAACACAGTGCACACCACGCCACGTTGCCTGACAACGGGCCACAACTCCTCATAA
AGAGACAGCAACCAGGATTTATACAAGGAGGAGAAAATGAAAGCCATACGGGAAGCAATAGCATGAT
ACAAAGGCATTAAAGCAGCGTATCCACATAGCGTAAAAGGAGCAACATAGTTAAGAATACCAGTCAA
TCTTTCACAAATTTTGTAATCCAGAGGTTGATTACCGATAAGCTTGATATCGAATTCTTACTTGTACAG
CTCGTCCATGCCGCCGGTGGAGTGGCGGCCCTCGGCGCGTTCGTACTGTTCCACGATGGTGTAGT
CCTCGTTGTGGGAGGTGATGTCCAACTTGATGTTGACGTTGTAGGCGCCGGGCAGCTGCACGGGCT
TCTTGGCCTTGTAGGTGGTCTTGACCTCAGCGTCGTAGTGGCCGCCGTCCTTCAGCTTCAGCCTCTG
CTTGATCTCGCCCTTCAGGGCGCCGTCCTCGGGGTACATCCGCTCGGAGGAGGCCTCCCAGCCCA
TGGTCTTCTTCTGCATTACGGGGCCGTCGGAGGGGAAGTTGGTGCCGCGCAGCTTCACCTTGTAGA
TGAACTCGCCGTCCTGCAGGGAGGAGTCCTGGGTCACGGTCACCACGCCGCCGTCCTCGAAGTTC
ATCACGCGCTCCCACTTGAAGCCCTCGGGGAAGGACAGCTTCAAGTAGTCGGGGATGTCGGCGGG
GTGCTTCACGTAGGCCTTGGAGCCGTACATGAACTGAGGGGACAGGATGTCCCAGGCGAAGGGCA
GGGGGCCACCCTTGGTCACCTTCAGCTTGGCGGTCTGGGTGCCCTCGTAGGGGCGGCCCTCGCCC
TCGCCCTCGATCTCGAACTCGTGGCCGTTCACGGAGCCCTCCATGTGCACCTTGAAGCGCATGAAC
TCCTTGATGATGGCCATGTTATCCTCCTCGCCCTTGCTCACCATCGGTCCAGGATTCTCTTCGACATC
TCCGGCTTGTTTCAGCAGAGAGAAGTTTGTTGCGCCGGATCCCCCGGGGAGCATGTCAAGGTCAAA
ATCGTCAAGAGCGTCAGCAGGCAGCATATCAAGGTCAAAGTCGTCAAGGGCATCGGCTGGGAGCAT
GTCTAAGTCAAAATCGTCAAGGGCGTCGGTCGGCCCGCCGCTTTCGCACTTTAGCTGTTTCTCCAGG
CCACATATGATTAGTTCCAGGCCGAAAAGGAAGGCAGGTTCGGCTCCCTGCCGGTCGAACAGCTCA
ATTGCTTGTTTCAGAAGTGGGGGCATAGAATCGGTGGTAGGTGTCTCTCTTTCCTCTTTTGCTACTTG
ATGCTCCTGTTCCTCCAATACGCAGCCCAGTGTAAAGTGGCCCACGGCGGACAGAGCGTACAGTGC
GTTCTCCAGGGAGAAGCCTTGCTGACACAGGAACGCGAGCTGATTTTCCAGGGTTTCGTACTGTTTC
TCTGTTGGGCGGGTGCCGAGATGCACTTTAGCCCCGTCGCGATGTGAGAGGAGAGCACAGCGGTA
TGACTTGGCGTTGTTCCGCAGAAAGTCTTGCCATGACTCGCCTTCCAGGGGGCAGGAGTGGGTATG
ATGCCTGTCCAGCATCTCGATTGGCAGGGCATCGAGCAGGGCCCGCTTGTTCTTCACGTGCCAGTA
CAGGGTAGGCTGCTCAACTCCCAGCTTTTGAGCGAGTTTCCTTGTCGTCAGGCCTTCGATACCGACT
CCATTGAGTAATTCCAGAGCAGAGTTTATGACTTTGCTCTTGTCCAGTCTAGACATGGTAATTCGATG
ATCCCCCTGGGGAGAGAGGTCGGTGATTCGGTCAACGAGGGAGCCGACTGCCGACGTGCGCTCCG
GAGGCTTGCAGAATGCGGAACACCGCGCGGGCAGGAACAGGGCCCACACTACCGCCCCACACCCC
GCCTCCCGCACCGCCCCTTCCCGGCCGCTGCTCTCGGCGCGCCCTGCTGAGCAGCCGCTATTGGC
CACAGCCCATCGCGGTCGGCGCGCTGCCATTGCTCCCTGGCGCTGTCCGTCTGCGAGGGTACTAG
TGAGACGTGCGGCTTCCGTTTGTCACGTCCGGCACGCCGCGAACCGCAAGGAACCTTCCCGACTTA
GGGGCGGAGCAGGAAGCGTCGCCGGGGGGCCCACAAGGGTAGCGGCGAAGATCCGGGTGACGCT

GCGAACGGACGTGAAGAATGTGCGAGACCCAGGGTCGGCGCCGCTGCGTTTCCCGGAACCACGCC
CAGAGCAGCCGCGTCCCTGCGCAAACCCAGGGCTGCCTTGGAAAAGGCGCAACCCCAACCCCGTG
GAATTATCACCTCGATCGGCTAGAACCGGTTGGAATTATCACCTCGATCGAGTTTACTCCCTATCAGT
GATAGAGAACGTATGAAGAGTTTACTCCCTATCAGTGATAGAGAACGTATGCAGACTTTACTCCCTAT
CAGTGATAGAGAACGTATAAGGAGTTTACTCCCTATCAGTGATAGAGAACGTATGACCAGTTTACTCC
CTATCAGTGATAGAGAACGTATCTACAGTTTACTCCCTATCAGTGATAGAGAACGTATATCCAGTTTA
CTCCCTATCAGTGATAGAGAACGTATAAGCTTTGCTTATGTAAACCAGGGCGCCTATAAAAGAGTGCT
GATTTTTTGAGTAAACTTCAATTCCACAACACTTTTGTCTTATACCAACTTTCCGTACCACTTCCTACC
CTCGTAAAGCGATCGCACTAGTCCAGTGTGGTGGAATTCTGCAGATATCCAGCACAGTGGCGGCCG
C

**Extended Data Table 9. LRC regulator cDNA library.**

| Gene | RefSeq ID | CCDS ID |
|---|---|---|
| ERG | NM_182918 | CCDS13658.1 |
| FLI1 | NM_002017 | CCDS44768.1 |
| ETS1 | NM_005238 | CCDS8475.1 |
| KLF2 | NM_016270 | CCDS12343.1 |
| KLF4 | NM_004235 | CCDS6770.2 |
| KLF6 | NM_001300 | CCDS7060.1 |
| JUN | NM_002228 | CCDS610.1 |
| JUNB | NM_002229 | CCDS12280.1 |
| JUND | NM_001286968 | |
| FOSB | NM_001114171 | CCDS46113.1 |
| FOSL2 | NM_005253 | CCDS1766.1 |
| GATA2 | NM_032638 | CCDS3049.1 |
| RUNX1 | NM_001001890 | CCDS42922.1 |
| MYB | NM_005375 | CCDS5174.1 |
| LYL1 | NM_005583 | CCDS12292.1 |
| MEF2C | NM_002397 | CCDS47245.1 |
| ZFP36L1 | NM_001244701 | |
| BTG2 | NM_006763 | CCDS1437.1 |
| EGR1 | NM_001964 | CCDS4206.1 |
| TagBFP | control | |

**Extended Data Table 10. Primer lists for cDNA screen library preparation**

| Forward Primer | Sequence |
|---|---|
| ORF_P5-SBS3_FW | AATGATACGGCGACCACCGAGATCTACACTCTTTCCCTACACGACGCTCTTCCGATCT |

| Reverse Primer | Sequence |
|---|---|
| ORF_P7_BC1_TruSeq | CAAGCAGAAGACGGCATACGAGATCGTGATGTGACTGGAGTTCAGACGTGTGCTCTTCCGATCT |
| ORF_P7_BC2_TruSeq | CAAGCAGAAGACGGCATACGAGATACATCGGTGACTGGAGTTCAGACGTGTGCTCTTCCGATCT |
| ORF_P7_BC3_TruSeq | CAAGCAGAAGACGGCATACGAGATGCCTAAGTGACTGGAGTTCAGACGTGTGCTCTTCCGATCT |
| ORF_P7_BC4_TruSeq | CAAGCAGAAGACGGCATACGAGATTGGTCAGTGACTGGAGTTCAGACGTGTGCTCTTCCGATCT |
| ORF_P7_BC5_TruSeq | CAAGCAGAAGACGGCATACGAGATCACTGTGTGACTGGAGTTCAGACGTGTGCTCTTCCGATCT |
| ORF_P7_BC6_TruSeq | CAAGCAGAAGACGGCATACGAGATATTGGCGTGACTGGAGTTCAGACGTGTGCTCTTCCGATCT |
| ORF_P7_BC7_TruSeq | CAAGCAGAAGACGGCATACGAGATGATCTGGTGACTGGAGTTCAGACGTGTGCTCTTCCGATCT |
| ORF_P7_BC8_TruSeq | CAAGCAGAAGACGGCATACGAGATTCAAGTGTGACTGGAGTTCAGACGTGTGCTCTTCCGATCT |
| ORF_P7_BC9_TruSeq | CAAGCAGAAGACGGCATACGAGATCTGATCGTGACTGGAGTTCAGACGTGTGCTCTTCCGATCT |
| ORF_P7_BC10_TruSeq | CAAGCAGAAGACGGCATACGAGATAAGCTAGTGACTGGAGTTCAGACGTGTGCTCTTCCGATCT |
| ORF_P7_BC11_TruSeq | CAAGCAGAAGACGGCATACGAGATGTAGCCGTGACTGGAGTTCAGACGTGTGCTCTTCCGATCT |
| ORF_P7_BC12_TruSeq | CAAGCAGAAGACGGCATACGAGATTACAAGGTGACTGGAGTTCAGACGTGTGCTCTTCCGATCT |
| ORF_P7_BC13_TruSeq | CAAGCAGAAGACGGCATACGAGATTTGACTGTGACTGGAGTTCAGACGTGTGCTCTTCCGATCT |
| ORF_P7_BC14_TruSeq | CAAGCAGAAGACGGCATACGAGATGGAACTGTGACTGGAGTTCAGACGTGTGCTCTTCCGATCT |
| ORF_P7_BC15_TruSeq | CAAGCAGAAGACGGCATACGAGATTGACATGTGACTGGAGTTCAGACGTGTGCTCTTCCGATCT |

**Extended Data Table 11. gRNA and primer lists used for CRISPR genome editing**

**a. gRNA lists**

| gRNA | Sequence | Reference |
|---|---|---|
| JUN gRNA-1 | CCATAAGGTCCGCTCTCGGA | Hs.Cas9.JUN.1.AA (IDT) |
| JUN gRNA-2 | CTCCTCACCTCGCCCGACGT | Hs.Cas9.JUN.1.AB (IDT) |
| JUN gRNA-3 | TGATAATCCAGTCCAGCAAC | Hs.Cas9.JUN.1.AC (IDT) |
| JUN gRNA-4 | CTGAACCTGGCCGACCCAGT | Hs.Cas9.JUN.1.AD (IDT) |
| JUN gRNA-5 | GCTCATCTGTCACGTTCTTG | Hs.Cas9.JUN.1.AE (IDT) |
| JUN gRNA-6 | GCCCCACGTCGGGCGAGGTG | Brunello Human CRISPR Knockout Pooled Library (addgene #73178)* |
| JUN gRNA-7 | GCTCTCGGACGGGAGGAACG | |
| JUN gRNA-8 | GGCGGCGCAGCCGGTCAACG | |
| JUN gRNA-9 | TGAACCTGGCCGACCCAGTG | |
| AAVS1 | GGGGCCACTAGGGACAGGAT | addgene plasmid #70661** |

*Doench et al. Nat Biotechnol. 2016 doi: 10.1038/nbt.3437.
**Wang T et al. Science 2015 doi: 10.1126/science.aac7041.

**b. Primer lists for gRNA-targeting site PCR amplification**

| Target site | Primer | Sequence |
|---|---|---|
| JUN gRNA-1 | Forward | CCCGTTGCTGGACTGGATTA |
| | Reverse | CTTTTCAAAGCCGGGTAGCG |
| JUN gRNA-2 | Forward | TCATCTGTCACGTTCTTGGGG |
| | Reverse | GTGCGTGCGCTCTTAGAGAA |
| JUN gRNA-7 | Forward | CCCGTTGCTGGACTGGATTA |
| | Reverse | CTTTTCAAAGCCGGGTAGCG |
| AAVS1 | Forward | ATCCTCTCTGGCTCCATCGT |
| | Reverse | AGCCACCTCTCCATCCTCTT |
